# General moment closure for the neutral two-locus wright-fisher dynamics

**DOI:** 10.64898/2026.01.16.700021

**Authors:** Raunak Kundagrami, Sean Yetter, Matthias Steinrücken

## Abstract

The Wright-Fisher diffusion and its dual, the coalescent process, are at the core of many results and methods in population genetics. Approaches have been developed to study the dynamics of its moments under genetic drift, mutation, and recombination using ordinary differential equations. The dynamics of these moments can be used to study population genetic processes and are key building blocks of efficient methods to infer population genetic parameters, like demographic histories or fine-scale recombination rates. However, the system of equations does not close under recombination; that is, computing moments of a certain order requires knowledge of moments of higher order. By applying a coordinate transformation to the diffusion generator, we show that the canonical moments in these alternative coordinates yield a closed system, enabling more accurate numerical computations. Compared to previous approaches in the literature, we believe that this approach can be more readily extended to general scenarios. Through simulations, we verify that the derived system of differential equations can accurately capture the dynamics of the moments, and can be used to efficiently compute expected diversity and linkage statistics in population genetic samples.

## 1. Introduction

The Wright-Fisher model and its diffusion approximation are central models in population genetics that describe the dynamics of genetic types or alleles in a population. At its core, the diffusion is a model of genetic drift, the random fluctuations of allele frequencies due to randomness in reproductive success, but other genetic processes like mutation, selection, recombination, or gene-flow can be readily incorporated. It is the starting point for many fundamental results in population genetics, for example, the fixation probabilities (Kimura, 1962) and times (Kimura and Ohta, 1969) of neutral or beneficial alleles, and the distribution of allele frequencies at stationarity (Kimura, 1964). Further, the diffusion is a central building block in many population genetic methods, including methods for demographic inference (Gutenkunst et al., 2009; Jouganous et al., 2017) and for detecting selection from time-series genetic-data (Tataru et al., 2017; Fine and Steinrücken, 2025).

The Wright-Fisher diffusion is commonly presented as a partial differential equation (PDE), the Kolmogorov forward equation, that describes the evolution of the density of the population allele frequencies over time (Karlin and Taylor, 1981, Chapter 15), or equivalently, as a differential operator that is the generator for the corresponding Markov process (Durrett, 2008, Chapter 7), or as a stochastic differential equation (for example, Jenkins and Spanò, 2017; He et al., 2020). While these PDEs can be used to derive many important quantities that describe the dynamics of genetic variation in a population, no analytic solutions exist in general. Numerical approaches have been implemented in specific scenarios of interest to obtain approximate solutions, and these approaches range from general purpose finite-difference methods (Gutenkunst et al., 2009; Ragsdale and Gutenkunst, 2017) to more specifically tailored spectral approaches (Lukić and Hey, 2012; Song and Steinrücken, 2012).

However, in many cases, the density of the allele or haplotype frequencies in the population is not of primary interest, but rather the corresponding moments. These can be common statistics used to describe genetic variation, for example, heterozygosity 2E[*X*_1_(*t*)(1 − *X*_1_(*t*))] or linkage disequilibrium (LD) E[*X*_11_(*t*)*X*_22_(*t*) − *X*_12_(*t*)*X*_21_(*t*)], where *X*_*i*_(*t*) and *X*_*ij*_(*t*) are the random allele and haplotype frequencies at time *t*, respectively. Other moments of interest are the probability of observing a certain configuration of alleles or haplotypes in a sample of a given size from the respective population, referred to as sampling probabilities, or in some cases allele- or site-frequency-spectra (SFS). These can be obtained as multinomial moments of the underlying population distribution of allele or haplotype frequencies. These moments are studied in populations at equilibrium, or their dynamics over time, for example, when selection is acting or the size of the population changes. The sampling probabilities in particular have been used in many methods that infer population genetic parameters from genetic variation sampled from natural populations (for example, Gutenkunst et al., 2009; Steinrücken, Birkner, et al., 2013; Spence and Song, 2019).

Previous approaches to compute these moments proceed in two steps (Williamson et al., 2005; Gutenkunst et al., 2009; Ragsdale and Gutenkunst, 2017). First, they use numerical approaches to solve the associated PDE for the density of the population frequencies and then integrate this density numerically to obtain the requisite moments. However, recent methods use a different framework to compute the moments more directly, circumventing possible accumulation of numerical inaccuracies in the two-step approach (Jouganous et al., 2017; Ragsdale and Gravel, 2019; Friedlander and Steinrücken, 2022). The main insight for this direct approach is that the PDE or the generator of the Wright-Fisher diffusion can be used to derive a system of ordinary differential equations (ODEs) that describes the dynamics of the moments. Solving these ODEs directly then results in a more accurate and efficient way to compute the dynamics of the moments of interest.

However, solving these ODEs also presents challenges. While the dynamics for genetic drift and mutation can be solved efficiently, the dynamics for recombination and selection do not close; that is, computing the derivative of the moments of a certain order *n* (exponent of moment or sample size) at a certain time *t* requires the moments for order *n* + 1 at that time, which in turn requires *n* + 2 and so forth. Thus, no exact solution is possible. Some approaches to address this problem proceed as follows. Given that the moments of order *n* at a certain time *t* have been computed, they can be extrapolated to “guess” the requisite moments of order *n* + 1. Substituting these extrapolated values in the ODEs for the moments of order *n* can then be used to compute an approximate solution for their dynamics. Since the accuracy of the approximation for order *n* + 1 from order *n* increases as *n* increases, this approach does require *n* to be sizeable, which can be challenging, since the number of moments grows quickly with the order. In addition, these approaches become less accurate when the respective parameters, the recombination rates or selection coefficients, are large.

An alternative approach is to seek a different set of moments where the dynamics closes, but that still allows us to extract the moments of interest. Focusing on the neutral scenario with recombination between two loci, each with two alleles, Ragsdale and Gravel (2019) present such a system, based on the work of Hill and Robertson (1968). Hill and Robertson (1968) noticed that when considering expected values of combinations of marginal allele frequencies at each locus, combined with expected squared LD, then the joint dynamics closes under genetic drift and recombination. Extending this work, Ragsdale and Gravel (2019) present a closed system of moments in terms of powers of LD and functions of the marginal allele frequencies, and use moments up to order 4 within and between sub-populations to derive a powerful method to infer complex demographic histories from contemporary genomic data (see also Ragsdale et al., 2023). While it is possible to find a set of moments that yields a closed system of ODEs for the dynamics under recombination, no such solution has been presented for the general dynamics including selection, and current approaches rely on numerical closure (Jouganous et al., 2017; Ragsdale, 2022; Friedlander and Steinrücken, 2022). A system of moments for which the selection dynamics is closed might not exist, but a formal proof of this statement is beyond the scope of the current manuscript. Interestingly, using the set of moments presented by Ragsdale and Gravel (2019), Barroso and Ragsdale (2026) show that in scenarios with negative selection, the heterozygosity at a linked neutral locus (moment of order 2) can be well approximated by truncating the moment hierarchy at a high order.

In this manuscript we focus on the neutral case with two loci separated by an arbitrary recombination distance in a single well-mixing population. The number of alleles at each locus is arbitrary, with arbitrary mutation rates between the alleles. We allow the rate of genetic drift to depend on time *t*, which enables us to model population size changing over time. The dynamics of the standard moments of the haplotype frequencies do not close in this setting (Ragsdale and Gravel, 2019; Friedlander and Steinrücken, 2022). To derive a set of moments that yield a closed system of ODEs, we first introduce a new coordinate system and transform the generator of the corresponding Wright-Fisher diffusion into this coordinate system. We then define canonical moments in this new coordinate system, and use the transformed generator to derive a closed system of ODEs for the dynamics of these moments under genetic drift, mutation, and recombination. The set of moments we present is also minimal without redundancies, which improves numerical efficiency.

Besides allowing a better assessment of the dynamics of two-locus moments of interest, and how these dynamics are affected by variable populations size, computation of these moments in non-equilibrium scenarios also has practical applications. First, several methods have been developed that infer population size history from patterns of LD observed in genomic samples (Tenesa et al., 2007; Waples and Do, 2010; Santiago et al., 2020), and Novo et al. (2022) suggest that these approaches can be more robust to selection than approaches based on genomic likelihoods. The framework detailed in this manuscript can be used to compute LD and higher order linkage statistics for different population size histories accurately and efficiently, thus presenting a promising approach to improve these inference methods. Second, many methods have been presented in the literature to infer fine-scale recombination rates from genomic data of contemporary populations (Auton and McVean, 2007; Chan et al., 2012), even accounting for variable population size (Spence and Song, 2019). These methods are based on composite likelihoods multiplying two-locus sampling probabilities, and are designed such that these two-locus probabilities are provided as lookup tables that can easily be substituted. The framework presented in this manuscript could thus be readily used to efficiently provide accurate lookup tables for any of these methods. Lastly, methods that estimate the population size history from LD require knowledge of the recombination rates, whereas Spence and Song (2019) present a method that estimate fine-scale recombination rates conditional on a given population size history. Fast and accurate methods that can compute two-locus moments under variable population size as presented here open up the potential for simultaneous joint inference of the recombination rates and the population size history.

Before we present the details of our approach, we want to highlight several connections to related approaches that have been presented in the literature. Extending work of Ohta and Kimura (1969), Mano (2013) presents a set of moments for the two-locus two-allele case in terms of powers of marginal allele frequencies and haplotype frequencies that yield a closed system of ODEs under genetic drift and recombination. This system arises as a special case of our system when restricted to two alleles and no mutation. Moreover, Ragsdale and Gravel (2019) extend work by Hill and Robertson (1968) to present a different set of moments based on powers of marginal allele frequencies and LD that close under the dynamics of genetic drift and recombination. While our work extends the general moment closure to an arbitrary number of alleles at both loci, our closed moments are also structurally different from Ragsdale and Gravel (2019): we also include variables for the marginals, but no explicit LD variables. We believe that this more clearly highlights that the crucial step is not including LD variables per se, but rather a sufficient number of marginal variables. In addition, we believe that the approach we present here allows for a more convenient extension to multiple alleles, as well as potential future extensions to an arbitrary number of loci.

Both Jenkins and Song (2012) and Kamm et al. (2016) present methods to compute two-locus sampling probabilities, that is, the probabilities of observing a certain configuration of haplotypes in a population sample of a given number of individuals. The former does so in a population at equilibrium, the latter under a model of piecewise constant population size changes. To this end, Jenkins and Song (2012) also apply a coordinate transformation to the generator of the corresponding Wright-Fisher diffusion. As with the transformation we present in this manuscript, their transformation also includes the marginal frequencies, but they scale the LD coordinates by the recombination rate. While their transformed generator yields a more tractable system of equations for the sampling probabilities, the system does not close, and the authors use Padé approximations to approximate the requisite probabilities. In related work, Fearnhead et al. (2015) highlight connections between the diffusion process described by the transformed generator of Jenkins and Song (2012) and corresponding coalescent processes with large recombination rates. Kamm et al. (2016) on the other hand compute the sampling probabilities using a time-reversible particle system that mirrors the coalescent with recombination. This process includes lineages where the genetic variation is only specified at one locus, which resembles the inclusion of marginal allele frequencies.

Lastly, in related work, Baake and Baake (2003) investigate the deterministic dynamics under mutation and recombination for an arbitrary number of loci with an arbitrary number of alleles, but without genetic drift. To obtain an analytic solution of the dynamics in this case, they transform the dynamics into a new coordinate system, where the coordinates are multi-locus LDs. We believe that canonical moments defined in this coordinate system would not only close but also diagonalize the recombination dynamics. However, it is unclear which form the resulting dynamics under genetic drift would take. We leave this interesting inquiry for future work. Moreover, Baake and Hustedt (2011) investigate the stochastic dynamics with genetic drift in this setting. They conjecture that the arbitrary multi-locus dynamics cannot be closed, but we believe that this conjecture should be revisited with the techniques presented in the present manuscript.

The remainder of this paper is organized as follows. In Section 2, we introduce the Wright-Fisher diffusion with recombination in the two-locus setting more formally. We present the generator modeling genetic drift, mutation, and recombination, and present the ODEs for the standard moments of the haplotype frequencies. This will allow us to clearly exhibit the moment-closure problem. In Section 3, we then introduce the new coordinate system and present the generator of the diffusion transformed to this new system. We define canonical moments in this new coordinate system and present the respective system of ODEs for these moments, which is closed under genetic drift, mutation, and recombination. We also provide a more detailed comparison between our moment-closure approach and other approaches presented in the literature. We validate our system of ODEs in Section 4 by comparing numerical solutions of the ODEs with simulations, for different mutation and recombination parameters, as well as several variable population size histories. On the one hand, we show that the trajectories of low order moments capturing genetic diversity and LD computed from the ODE align well with the corresponding statistics estimated from simulated trajectories. On the other hand, we use genomic simulations with the tool `msprime` (Baumdicker et al., 2022), a state-of-the-art coalescent simulator, to obtain two-locus sampling probabilities, specifically the two-locus SFS, and demonstrate that solutions to our ODEs can be used to compute the same probabilities accurately and efficiently. We discuss possible applications and extensions of our approach to scenarios with multiple loci or selection in Section 5. All code to perform the simulations and numerical computations, as well as code to recreate the figures in this manuscript is available at: https://github.com/steinrue/two_locus_closure.

## 2. Model

### 2.1. Wright-Fisher diffusion

We consider the Wright-Fisher diffusion (Ewens, 2004, Chapter 5; Durrett, 2008, Chapter 7) for two linked loci, with an arbitrary number of alleles at each locus. We denote by *K* the number of alleles at the first locus and by *E*^(1)^ := {1, …, *K}* the set of possible alleles. Similarly, we denote by *L* the number of alleles at the second locus and by *E*^(2)^ := {1, …, *L}* the set of possible alleles. Thus, the set of the *K* × *L* possible two-locus haplotype configurations is given by 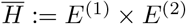. In what follows, we will also need to refer to the set of haplotype configurations that excludes, without loss of generality, the haplotype (*K, L*), and we denote this set by 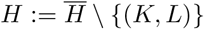.

Denote by 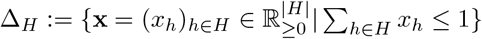 the simplex of non-negative haplotype frequencies *x*_*h*_ for h ∈ *H* whose sum is less than or equal to one. We use the convention that the frequency of the excluded haplotype (*K, L*) ∈*/ H* is given by

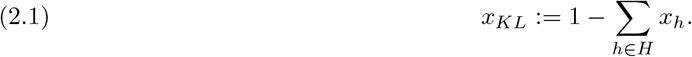

For given initial haplotype frequencies **x**^(0)^ ∈ Δ_*H*_, the Wright-Fisher diffusion is a Markov process {**X**(*t*)}_*t*≥0_, where **X**(*t*) ∈ Δ_*H*_ and **X**(0) = **x**^(0)^, that describes the temporal dynamics of the haplotype frequencies in the population subject to given population genetic processes.

The dynamics of the Wright-Fisher diffusion can be characterized in terms of the Kolmogorov backward equation, that is, for **x, y** ∈ Δ_*H*_, its transition density function

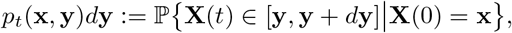

is the solution of the partial differential equation (PDE)

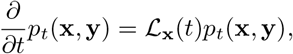

for a given second order differential operator ℒ_**x**_(*t*) in **x** that can depend on *t*. Here we use the notation **X**(*t*) ∈ [**y, y** + *d***y**] to refer to the event that **X**(*t*) is in an infinitesimal neighborhood around **y**. The domain of this operator is *C*_2_(Δ_*H*_), the twice continuously-differentiable real-valued functions on Δ_*H*_ . Note that common alternative but equivalent ways to characterize the Wright-Fisher diffusion are in terms of the corresponding Kolmogorov forward equation; using the differential operator ℒ_**x**_(*t*) as the generator of the Markov process; defining the process via the corresponding martingale problem; or by defining the corresponding stochastic differential equation.

The specific population genetic scenario modeled by the Wright-Fisher diffusion is determined by the exact form of ℒ_**x**_(*t*). Since we are interested in using the Wright-Fisher diffusion for the neutral dynamics of the frequencies of two-locus haplotypes under recurrent mutation and recombination, the operator takes the form

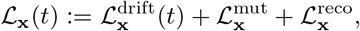

where the respective parts are defined as follows. Note that only the operator for drift depends on *t*, which allows modeling time-varying strength of drift, which is equivalent to time-varying effective population size. We will also frequently use the Kronecker delta *δ*_*hg*_, which is 1 if *h* = *g*, and 0 otherwise, as well as the indicator function **1**_*A*_, which is 1 if condition *A* is true and 0 otherwise. Moreover, we will denote the frequency of the haplotype *h* = (*i, j*) by *x*_*h*_ or *x*_*ij*_, depending on the context.

#### Definition 2.1

(Genetic drift). Denoting the effective number of diploid individuals in the population at time *t* by *N*_*e*_(*t*) *>* 0, the second order differential operator

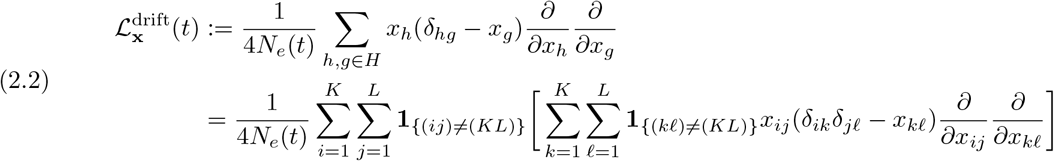

captures the random fluctuations of the haplotype frequencies due to genetic drift, see Equation (2) by Friedlander and Steinrücken (2022).

#### Definition 2.2

(Recurrent mutation). Denote by *m*_*h*,*g*_ ≥ 0 the rate of mutation from haplotype 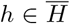 to haplotype 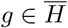. Then,

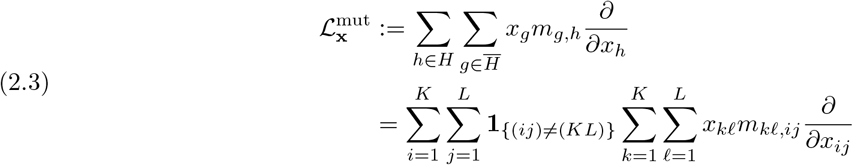

describes the change in the haplotype frequencies due to mutation, see Equation (3) by Friedlander and Steinrücken (2022). Here we used

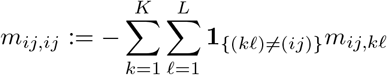

and equation (2.1). In equation (2.3), the change of the haplotype frequency *x*_*h*_ is given by the influx from all types that can mutate into *h*, represented by the terms of the inner sum with *g* ≠ *h*, and the outflux to all types that *h* can mutate into, given by the term with *g* = *h*.

Note that in Definition 2.2, we use a very general mutation model, where each two-locus haplotype can mutate into any other haplotype. A common assumption is that mutations are locus-specific and rare, so that at most one locus is subject to a mutation at a given time. Specifically, denote by 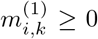 the rate that at the first locus, allele *i* mutates into allele *k* ≠*i*, and by 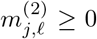 the rate of a *j* to *ℓ*≠ *j* mutation at the second locus. Assuming mutation to be locus-specific, and neglecting mutations at both loci at the same time, we get

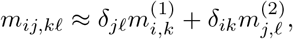

and particularly

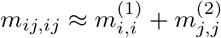

with

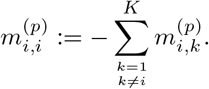

Substituting these approximations into the definition of the mutation operator in equation (2.3), we obtain

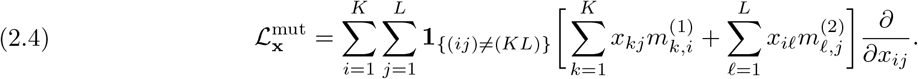

We will use this simplified version of the mutation operator in most of the remainder.

#### Definition 2.3

(Recombination). Denote by *r* ≥ 0 the rate that a recombination event occurs between the two loci considered here. Then the operator

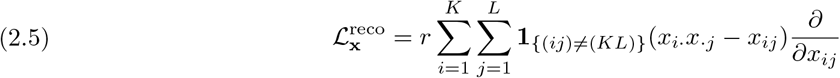

describes the change of the haplotype frequencies due to recombination, see Equation (5) by Friedlander and Steinrücken (2022). Here, 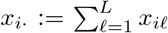 and 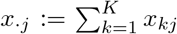 denote the marginal allele frequencies of *i* at the first locus and *j* at the second locus, respectively. These dynamics essentially decouple allelic associations between the two loci over time.

#### Remark 2.4.

We include the factor 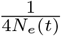 in Definition 2.1 for the operator representing genetic drift. Thus, time *t* is measured in terms of generations, and the mutation and recombination rates are equal to the per generation probabilities of the respective events.

Note that here, we introduced the Wright-Fisher diffusion as a stochastic process on Δ_*H*_, the positive haplotype frequencies for *h* ∈ *H* that sum to 1 or less, and we implicitly used *x*_*KL*_ = 1 − ∑_*h*∈*H*_ *x*_*h*_. Thus, the respective differential operators are acting on *C*_2_(Δ_*H*_), and consequently, the sums in the definitions are summing the differentials 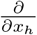 over *h* ∈ *H*. Alternatively, some references in the literature introduce the Wright-Fisher diffusion as a stochastic process on 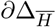, the boundary of the simplex 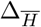; that is, the positive haplotype frequencies for all 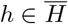 that sum to exactly 1. The associated differential operators are acting on 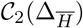, and thus, the summations over the differentials 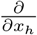 extend to 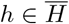.

Essentially, the question is whether the diffusion of dimension 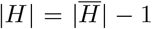 is embedded in a space of dimension 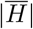, the possible number of genetic types, or whether it is embedded in a space of dimension |*H*|, the number of types minus 1. The most prominent case is the diffusion for a single locus with two alleles. This 1-dimensional diffusion is almost exclusively presented in the literature (Ewens, 2004, Chapter 5; Durrett, 2008, Chapter 7) embedded in a 1-dimensional space, that is, the number of alleles minus 1. In more general cases, Jenkins and Song (2012) and Friedlander and Steinrücken (2022) use the number of alleles, whereas Mano (2013) and Steinrücken, Wang, et al. (2013) use the number of alleles minus 1. In Section S.2.1, Section S.2.2, and Section S.2.3 in Supplemental Material, we show using *x*_*KL*_ = 1 ∑_*h* ∈ *H*_ *x*_*h*_ that the two respective formulations of the drift, the mutation, and the recombination operator are equivalent, and they can thus be used interchangeably. While this result is likely not surprising to readers familiar with this framework, we are unaware of an explicit statement of this result in the literature, and thus include it here for convenience. In the remainder, we will proceed with the operators on *C*_2_(Δ_*H*_), thus using number of alleles minus 1 as the dimension of the embedding space, since this version will be more convenient for the derivation of our results.

### 2.2. Moments of the Wright-Fisher diffusion

The Wright-Fisher diffusion models the random dynamics of the haplotype frequencies in the given population. However, our main interest here is to investigate the dynamics of the following moments of the diffusion.

#### Definition 2.5

(Moments *M*). With 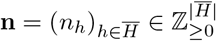 and **x**^(0)^, **y** ∈ Δ_*H*_, define

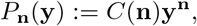

where

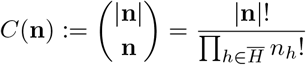

denotes the multinomial coefficient, with 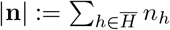, and

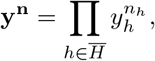

with the convention that *y*_*KL*_ = 1 − ∑ _*h*∈*H*_ *y*_*h*_. We then define the **n**-moment of the Wright-Fisher diffusion initialized at **x**^(0)^ as

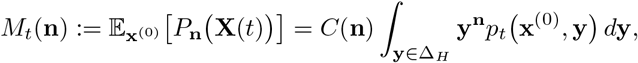

where we use the notation 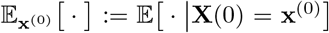 and omit the dependence of *M*_*t*_(**n**) on the initial condition **x**^(0)^ for brevity.

#### *Remarks* 2.6.

i. The moment *M*_*t*_(**n**) is the probability of observing *n*_*h*_ haplotypes of type *h* for all 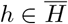 when sampling 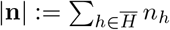 haplotypes from the population at time *t*, since this probability is given by multinomial sampling conditional on the underlying population haplotype frequencies.
ii. The definition of these canonical moments does not need to include the multinomial coefficient *C*(**n**), however, we include it here to make the connection to the sampling probabilities more explicit.

As discussed in Section 1, in many applications, the detailed dynamics of the population-level haplotype frequencies is not of primary interest, but rather the moments of the diffusion. Following Friedlander and Steinrücken (2022), a key step to compute these moments is Dynkin’s formula (Karlin and Taylor, 1981, Chapter 15, Section 11; Øksendal, 2003, Lemma 7.3.2). It states that for a given diffusion {**X**(*t*)}_*t*≥0_ with generator ℒ_**x**_(*t*) and a function *f* ∈ *C*_2_(Δ_*H*_), the relation

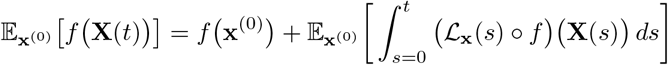

holds, where **x**^(0)^ ∈ Δ_*H*_ is the initial condition, which then yields the ODE

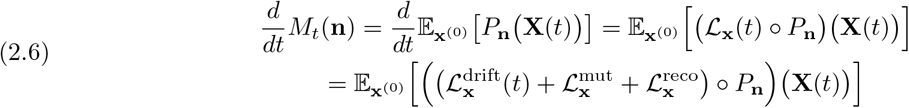

The following corollaries describing the respective contributions to the ODE for *M*_*t*_(**n**) from genetic drift, recurrent mutation, and recombination thus follow from Theorem 2 by Friedlander and Steinrücken (2022) and the results in Section S.2 in Supplemental Material about the equivalence of the different generator definitions used here and by Friedlander and Steinrücken (2022).

#### Corollary 2.7

(Genetic drift). *For given* **n** ∈ Z^|*H*|^, *the contribution of genetic drift to the ODE for M*_*t*_(**n**) *is given by*

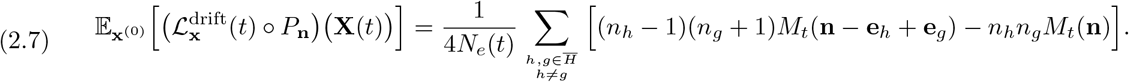

*Here*, **e**_*h*_ *denotes the unit vector of all 0, except 1 at index h*.

#### Corollary 2.8

(Recurrent mutation). *With* 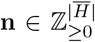, *the contribution of recurrent mutation to the ODE for M*_*t*_(**n**) *is given by*

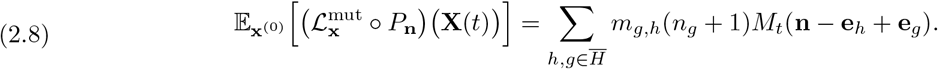

#### Corollary 2.9

(Recombination). *Let* 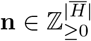, *then*

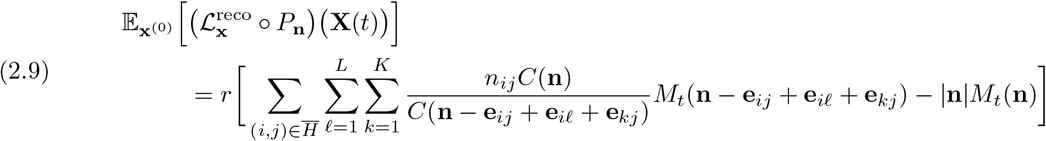

*is the contribution of recombination to the ODE for M*_*t*_(**n**).

#### Remark 2.10

In the expressions on the right-hand sides of equation (2.7), equation (2.8), and equation (2.9), some exponents of moments are decreased by 1. This could technically result in moments with negative exponents. However, here and in the remainder, we implicitly only include moments where all exponents are non-negative to keep the notation concise, and do not include explicit indicator functions. The reason that no terms with negative exponents have to be considered is that the derivations include taking derivatives of polynomials, and the derivative of a polynomial with exponent zero vanishes.

Corollary 2.7, Corollary 2.8, and Corollary 2.9 allow us to exhibit the problem of moment-closure explicitly. To this end, let *n* ∈ ℤ_≥0_ be a given moment order, and denote by

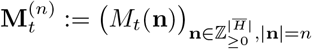

the vector of all moments *M*_*t*_(**n**) with |**n**| = *n*, that is, of order *n*. If we have suitable initial conditions 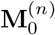 and want to compute 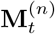 under the dynamics for genetic drift and recurrent mutation, we can implement a numerical scheme to solve the system of ODEs specified by equation (2.7) and equation (2.8), since all moments on the right-hand side of these equations are also of order *n*.

However, for recombination, this is not the case. Some moments on the right-hand side of equation (2.9) are of order *n* + 1. Thus the solution of the system of ODEs for order *n* depends on order *n* + 1, which in turn depends on order *n* + 2, and so forth. We therefore cannot readily solve this system using numerical approaches. In the next section, we will introduce alternative moments that yield a closed system.

## 3. Main results

To solve the moment-closure problem for the recombination dynamics, we proceed in two steps. We first introduce a new coordinate system and derive the representation of the differential operator ℒ_**x**_(*t*) in this coordinate system. We then define canonical moments based on this new coordinate system, and show that these moments close under genetic drift, recurrent mutation, and recombination.

### 3.1. Change of coordinate system

In Section 2.1, we introduced the Wright-Fisher diffusion **X**(*t*) ∈ Δ_*H*_ as a stochastic process in the coordinate system Δ_*H*_, the non-negative frequencies *x*_*h*_ of haplotypes *h* ∈ *H* summing to 1 or less, and with *x*_*KL*_ = 1 − ∑_*h*∈*H*_ *x*_*h*_. We now introduce a different coordinate system to describe these frequencies, in which the structure of genetic drift and recurrent mutation is not substantially changed, but recombination is more amenable to derive moments that yield a closed system of ODEs. To this end, note that the marginal allele frequencies 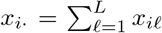 and 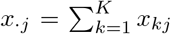 occur in the differential operator describing recombination in equation (2.5). We thus represent these marginal frequencies as explicit coordinates in our new coordinate system.

Formally, using the notation [*K* − 1] := {1, …, *K* − 1}, each element **x** ∈ Δ_*H*_ is given as a vector

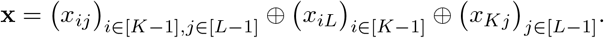

For given *K* and *L*, elements **z** in the new coordinate system are given by a vector

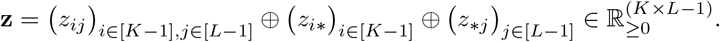

In the coordinates **z**, the *z*_*ij*_ represent haplotype frequencies, but for fewer combinations of alleles than *x*_*ij*_. The additional coordinates *z*_*i*∗_ and *z*_∗*j*_ represent an appropriate number of marginal frequencies that still allow the state of the diffusion originally represented in **x** to be uniquely represented in **z**. Note that the notations *x*_*i*·_ and *x*_·*j*_ represent the function of computing marginal frequencies from haplotype frequencies,

whereas *z*_*i*∗_ and *z*_∗*j*_ represent explicit coordinate in the **z**-system (not functions). The components of this vector have to satisfy

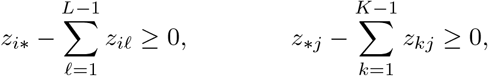

and

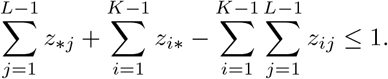

We denote the set of vectors satisfying these conditions by Ƶ_*K*,*L*_. The mapping **z**(**x**) from **x** ∈ Δ_*H*_ to **z** ∈ Ƶ_*K*,*L*_ is given by

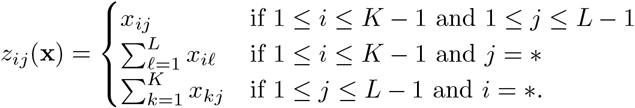

Furthermore, the mapping **x**(**z**) from **z** ∈ Ƶ_*K*,*L*_ to **x** ∈ Δ_*H*_ is given by

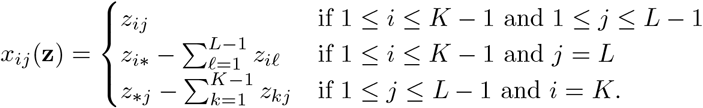

Figure 1 shows a schematic of this coordinate transform.

**FIGURE 1.**
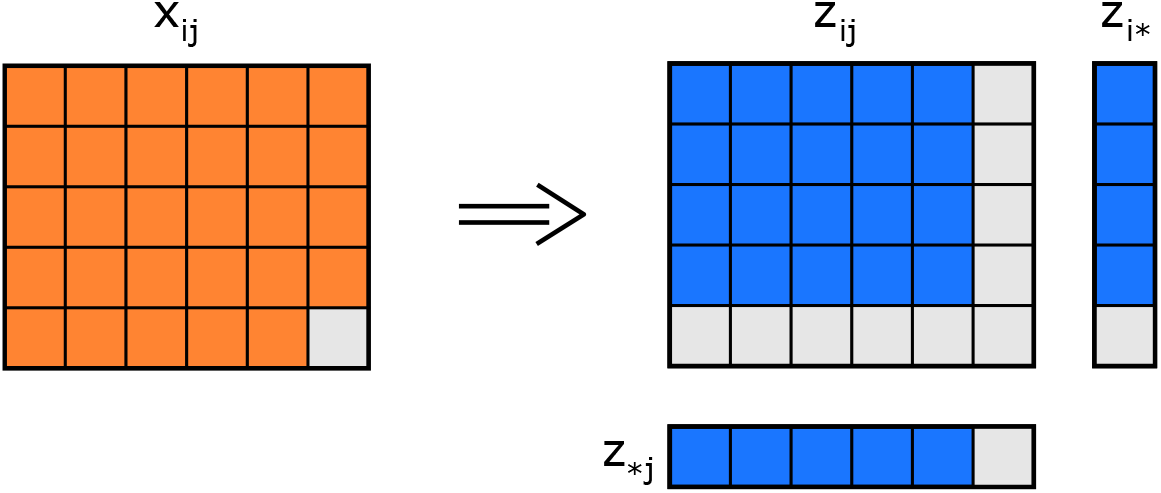
Schematic of the coordinate transformation. In the **x** coordinates, all *x*_*ij*_ are included (orange squares), except the highest index *x*_*KL*_ (gray square). The **z** coordinates include all *z*_*ij*_, all *z*_*i*∗_, and all *z*_∗*j*_ with *i < K, j < L* (blue squares). The coordinates with *i* = *K* or *j* = *L* are excluded (gray squares).

Note that **x**(**z**(**x**)) = **x** for all *x* ∈ Δ_*H*_ and **z**(**x**(**z**)) = **z** for all **z** ∈ Ƶ_*K*,*L*_, thus the two mappings are inverse to each other. Moreover, both mappings are continuously differentiable bijections, and thus they are diffeomorphisms. The differential operator ℒ_**x**_(*t*) in **x** defines the Wright-Fisher diffusion **X**(*t*), and we now derive the differential operator ℒ_**z**_(*t*) in **z** that describes the diffusion **z**(**X**(*t*)). To this end, note that using the total derivative with the chain rule, we obtain

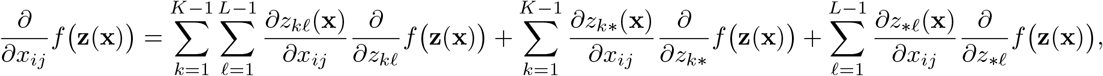

which yields

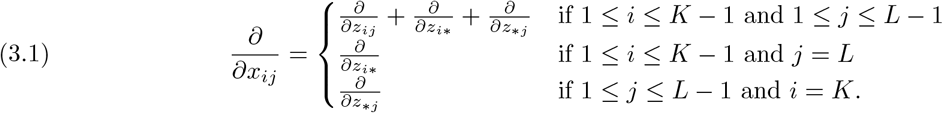

Substituting these differentials into the differential operators 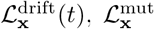, and 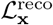, we obtain the following differential operators that characterize the Wright-Fisher diffusion in the coordinates **z**.

#### Lemma 3.1

(Genetic drift). *Denoting by N*_*e*_(*t*) *the effective diploid population size at time t, we obtain the second order differential operator*

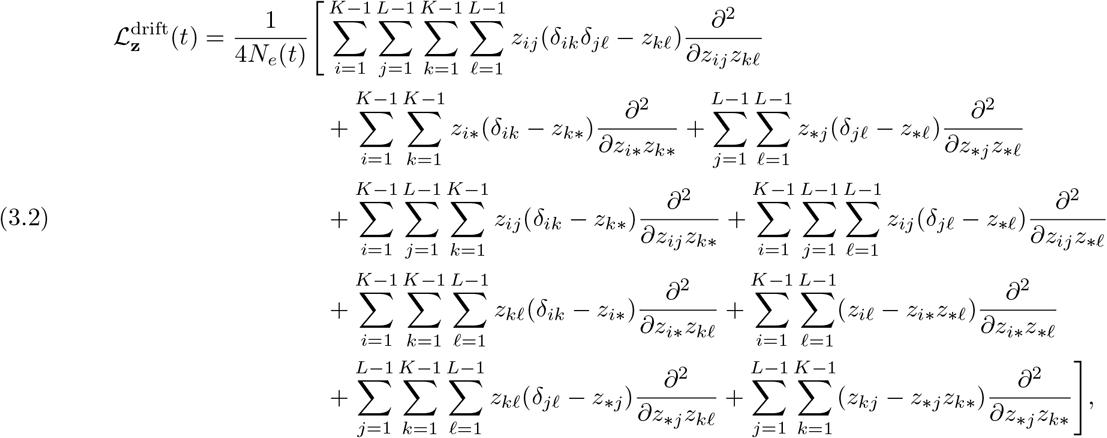

*which describes the change in haplotype frequencies due to genetic drift, where the frequencies are described using the coordinate system* **z**(**x**).

#### Lemma 3.2

(Recurrent mutation). *Denoting by* 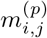 *the rate that allele i mutates to allele j*≠ *i at locus p, the generator*

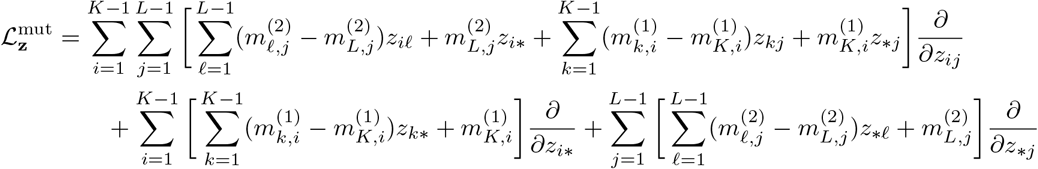

*describes the change in haplotype frequencies due to locus-specific recurrent mutation, where the frequencies are described using the coordinate system* **z**(**x**).

#### Lemma 3.3

(Recombination). *With r the rate of a recombination event occurring between the two loci of interest, we can write*

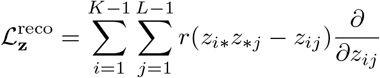

*to describe the change in haplotype frequencies due to recombination, where the frequencies are described using the coordinate system* **z**(**x**).

We provide proofs of Lemma 3.1, Lemma 3.2, and Lemma 3.3 in Section S.1.1 of Supplemental Material.

### 3.2. Closed moments

We now introduce the canonical set of moments in the coordinate system **z** and show that the resulting system of ODEs under drift, mutation, and recombination is closed.

#### Definition 3.4

(Moments *G*). Denote by {**Z**(*t*)}_*t*≥0_ the Wright-Fisher diffusion in the **z** coordinate system and by **z**^(0)^ ∈ Ƶ_*K*,*L*_ the initial configuration. With 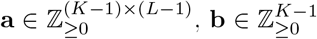, and 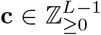, define the moments of the diffusion as

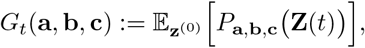

where

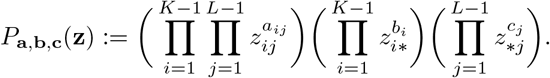

To derive the ODEs for the moments *G*_*t*_(**a, b, c**), we use the differential operator

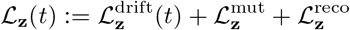

in Dynkin’s formula (2.6) and obtain

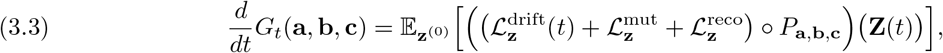

The following theorems describe the respective contributions to the ODE for *G*_*t*_(**a, b, c**) from genetic drift, recurrent mutation, and recombination.

#### Theorem 3.5

(Genetic drift). *For given* 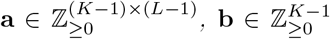, *and* 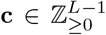, *the contribution from genetic drift to the ODE for G*_*t*_(**a, b, c**) *in the* **z** *coordinate system is*

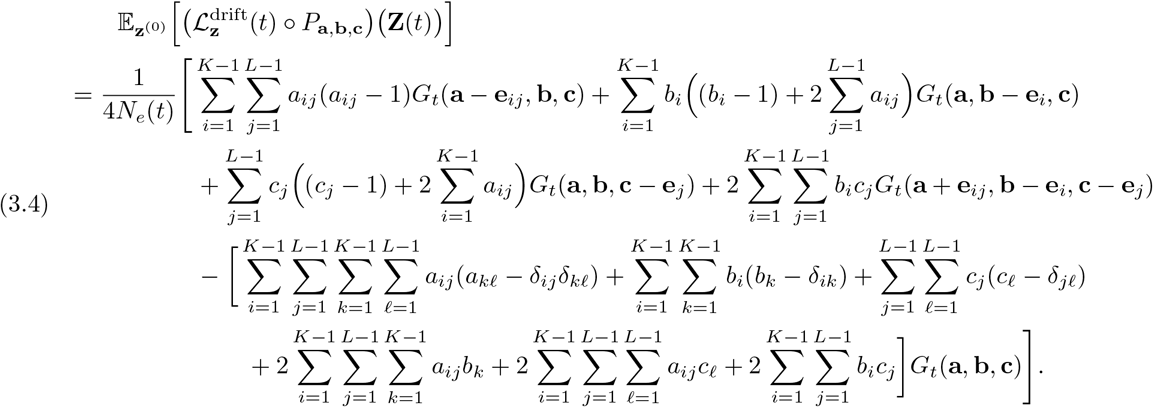

#### Theorem 3.6

(Recurrent mutation). *For given* 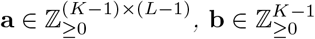, *and* 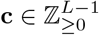, *the contribution from recurrent mutation to the ODE for G*_*t*_(**a, b, c**) *is given by*

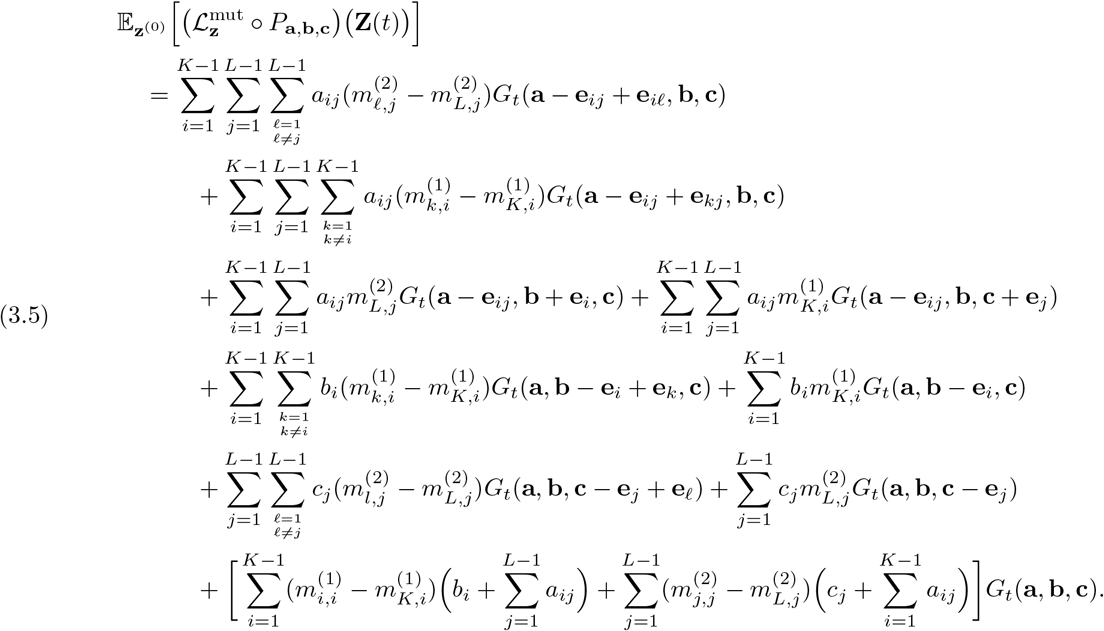

#### Corollary 3.7

(Parent-independent recurrent mutation). *The result in Theorem 3.6 can be further simplified in the case of parent-independent recurrent mutation (Durrett, 2008, Chapter 8.1.2). Under this assumption*, 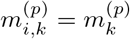 *for all i, k* ∈ {1, …, *K*} *with i*≠ *k, and we can rewrite equation 3.5 as*

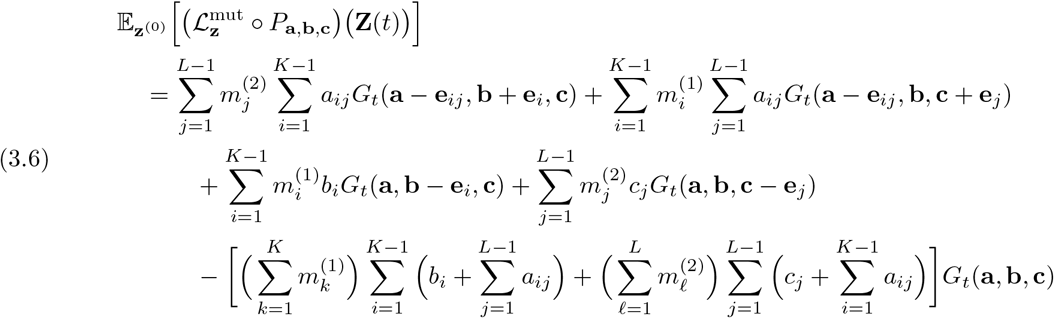

*to obtain the contribution from parent-independent mutation to the ODE for G*_*t*_(**a, b, c**).

The proofs for Theorem 3.5, Theorem 3.6, and Corollary 3.7 are provided in Section S.1.2 of Supplemetal Material.

#### Theorem 3.8

(Recombination). *The contribution to the ODE for G*_*t*_(**a, b, c**) *from recombination is given by*

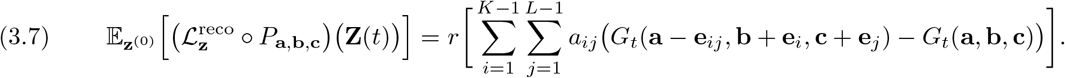

*Proof*. Applying the differential operator 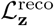 to the moment polynomial *P*_**a**,**b**,**c**_(**z**) and taking the expectation with respect to **Z**(*t*) yields the right-hand side of equation (3.7). The statement of the theorem follows from Dynkin’s formula. □

#### Remark 3.9

Similar to Remark 2.10, we implicitly only include moments where all exponents are non-negative on the right-hand side of equation (3.4), equation (3.5), equation (3.6), and equation (3.7). Note that here the corresponding multiplicative factors of the respective terms are actually zero in the correct cases.

The following lemma states the closure result for the system of ODEs defined by *G*_*t*_(**a, b, c**) explicitly.

#### Lemma 3.10

(Moment-closure). *Let the moment-order of G*_*t*_(**a, b, c**) *be* (*a, b, c*) = (|**a**|, |**b**|, |**c**|), *and define*

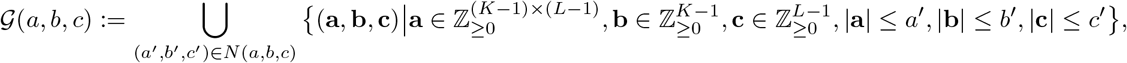

*where*

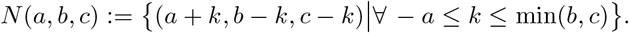

*Define*

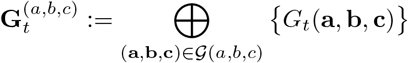

*as the vector of all moments whose orders are from the set G*(*a, b, c*). *Then, the system of ODEs for* 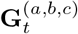 *defined by the dynamics* (3.3) *is closed*.

*Proof*. Observe that for any moment with exponents (**a, b, c**) ∈ *G*(*a, b, c*), the exponents in the sums on the corresponding right-hand sides of equations (3.4), (3.5), (3.6), and (3.7) are also in the set *G*(*a, b, c*), and thus the system of ODEs is closed. A critical property here is that, while some terms on the right-hand side increase certain exponents, they also decrease other exponents by the same amount. Thus, these terms can only grow the former exponents until the latter exponents reach 0, and the system cannot grow unboundedly. Specifically, genetic drift does not result in a net increase of exponents, except in one case, where it can increase **a**, but then has to decrease **b** and **c** at the same time. Mutation does not result in a net increase of exponents. Recombination can decrease **a**, but has to increase **b** and **c** at the same time (reversing the action of drift in this case). □

The closure of the moments *G*_*t*_(**a, b, c**) allows us to characterize the stationary solution of the system more explicitly.

#### Corollary 3.11

(Moments at stationarity). *If N*_*e*_(*t*) = *N*_*e*_ *>* 0 *and* 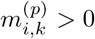 *for all p, i, k, then the system has a stationary distribution, which we obtain by setting the derivatives to zero, that is*,

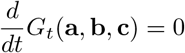

*for all* (**a, b, c**) ∈ *G*(*a, b, c*). *This is equivalent to solving the linear system*

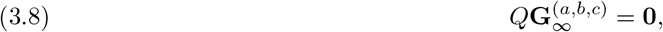

*where* 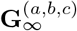 *denotes the moments at stationarity and the coefficients in the matrix Q are given by the coefficients in the summations of equations* (3.4), (3.5), (3.6), *and* (3.7).

#### Remark 3.12

Note that for the case of parent-independent mutation, equation (3.8) recovers recursions presented by Golding (1984, Appendix 1) and Ethier and Griffiths (1990, Equation (2.2)).

### 3.3 Comparison to related approaches in the literature

Many approaches in the literature consider the case of two loci with two alleles each. In our notation, this corresponds to *K* = 2, *L* = 2, and 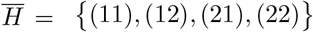. In this case we have **a** = (*a*_11_), **b** = (*b*_1_), and **c** = (*c*_1_); that is, the collections of exponents are given by a single number ∈ ℤ_≥0_ each. Thus, we can write

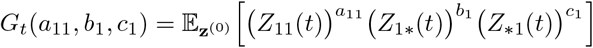

for the closed moments of the Wright-Fisher diffusion. Furthermore the ODEs for the closed moments *G*_*t*_(*a*_11_, *b*_1_, *c*_1_) are given by

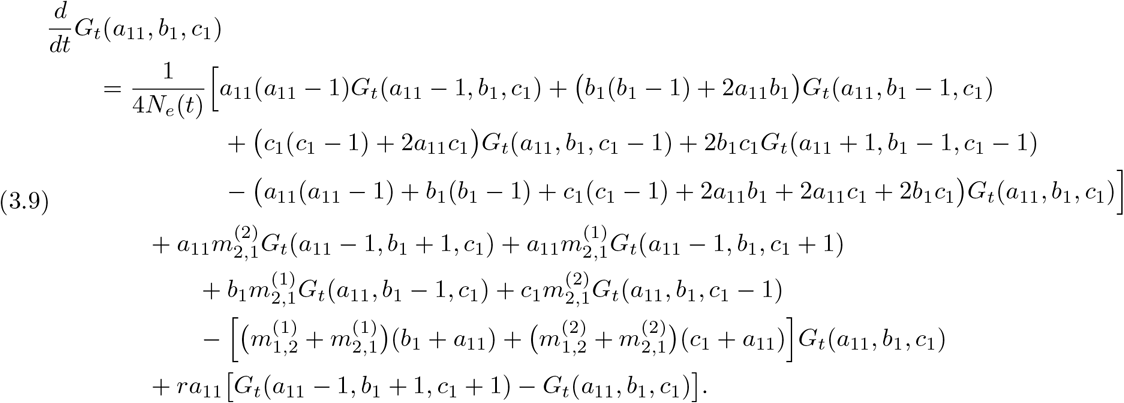

The first three lines on the right-hand side of equation (3.9) represent genetic drift, the next three lines recurrent mutation, and the last line represents recombination. Note that, without mutation 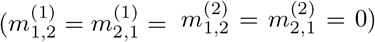, equation (3.9) is equal to Equation (3.2) by Mano (2013), multiplied by 4*N*_*e*_(*t*), where *n* = *a*_11_, *l* = *b*_1_, *m* = *c*_1_, *ξ*_*l*,*m*,*n*_ = *G*_*t*_(*n, l, m*), and *ρ* = 4*N*_*e*_(*t*)*r*.

Extending the approach of Hill and Robertson (1968), Ragsdale and Gravel (2019, S1 Appendix, Section S1.4.1) present an alternative collection of moments

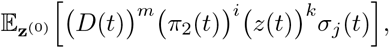

for *m, i, k, j* ∈ ℤ_≥0_, where

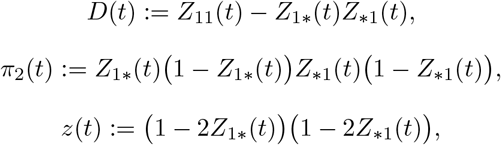

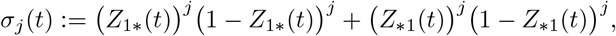

and show that these moments closes under genetic drift and recombination. In addition, the authors conjecture that moments of the form

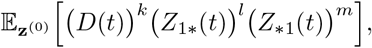

with *k, l, m* ∈ ℤ_≥0_ will result in a closed system of ODEs as well, which we also believe to be correct.

A detailed comparison of these different collections of moments is beyond the scope of this manuscript. However, since the different collections close the dynamics of genetic drift and recombination, they can likely be related to each other through linear transformations. Moreover, it should be possible to reduce each collection of moments to a minimal set. By this we refer to the fact that, for example, our moments *G*_*t*_(*a*_11_, *b*_1_, *c*_1_) do not include explicit powers of *Z*_22_(*t*), since they can be expressed as linear combinations of powers of *Z*_11_(*t*), *Z*_1∗_(*t*), and *Z*_∗1_(*t*), which are used to define *G*_*t*_(*a*_11_, *b*_1_, *c*_1_). We believe that each collection of moments that closes the dynamics can be reduced to such a minimal set, which will be of equal dimension for each collection. Thus, all sets will likely have comparable computational efficiency when solving the dynamics. However, in the approach of Ragsdale and Gravel (2019), the ODE for 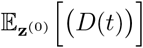, for example, only depends on this moment. This example shows that the system of ODEs for the different collections of moments could potentially be more or less sparse, and thus result in better numerical properties. Another aspect that might influence the choice of which moments to use are the specific properties of the solution needed for a certain application. The moments presented here are the only moments that can accommodate mutation with more than two alleles. Despite this, the moments of Ragsdale and Gravel (2019) include heterozygosity and LD as explicit variables, and thus might be better suited when studying these statistics. In contrast, the moments presented here more readily yield probabilities of configurations of sampled haplotypes, see Section 4.2.

Lastly, note that in the case without mutation, the coefficients on the right-hand side of equation (3.9) resemble the transition rates presented by Simonsen and Churchill (1997) for the two-locus, two-allele coalescent with recombination. In their manuscript, the authors define a backward-in-time Markov chain for the coalescent with recombination, with three different types of lineages: (1) alleles specified at both loci, (2) allele specified at first locus, and (3) allele specified at second locus. These partially specified lineages mirror the fact that moments with *b*_1_ marginalize over the second locus, and moments with *c*_1_ marginalize over the first. The coefficients of the moments in the first two lines for drift in equation (3.9) thus agree with the coalescent rates given by Simonsen and Churchill (1997) for the respective coalescent events of certain types of lineages, and the coefficient of the first recombination term in equation (3.9) agrees with the rate of splitting a lineage of type (1) into one lineage of type (2) and one lineage of type (3), when we account for the rescaling of the coalescent time used by Simonsen and Churchill (1997).

The terms representing mutation on the right-hand side of equation (3.9) resemble the dynamics of the coalescent with “killing”, defined by Durrett (2008, Ch. 1.3.1) or Kamm et al. (2017, Section 3.4). In the first two mutation terms, a type (1) lineage (allele is specified at both loci) is affected by a mutation at one of the two loci, resulting in the loss of a type (1) lineage and the gain of a partially specified type (2) or (3) lineage. In the next two terms, a type (2) or (3) lineage is affected by a mutation event, resulting in the loss of that lineage. However, note that such interpretations are only valid for the parent-independent mutation models in Corollary 3.7 and equation (3.9). In the general case of recurrent mutation in Theorem 3.6, mutation events do not “kill” lineages, but rather change the genetic type of a lineage. The coefficients of the terms *G*_*t*_(**a, b, c**) in equation (3.9) are the total sum of all rates. These properties are a manifestation of the well-established moment duality between diffusion and coalescent processes.

## 4. Numerical results

For the numerical results in this section, we again consider the case of two loci with two alleles each, that is, *K* = 2, *L* = 2, and 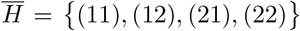, since many population genetic applications focus on analyzing Single Nucleotide Polymorphisms (SNPs), which are often considered biallelic. As detailed in Section 3.3, in this case we have **a** = (*a*_11_), **b** = (*b*_1_), and **c** = (*c*_1_) with *a*_11_, *b*_1_, *c*_1_ ∈ ℤ_≥0_. We can write

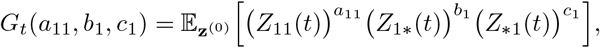

and the dynamics of these moments is given by the ODE (3.9). In the remainder of this section, we will also only consider scenarios where the mutation rates at both loci and between both alleles are the same, that is, 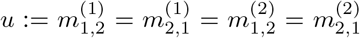. All numerical solutions of the ODEs presented here are implemented in `python` using the function `scipy.integrate.odeint()` (v1.16.2) with default parameters, except we set `hmax` to 1, so that the solver stops at the designated final time. The linear system to obtain the stationary distribution is solved using the function `numpy.linalg.solve()` (v2.3.3) with default parameters.

### 4.1. Comparison against simulated trajectories

In Figures 3, 4, and 5 we compare trajectories simulated using the Wright-Fisher model to solutions of the system of ODEs for *G*_*t*_(*a*_11_, *b*_1_, *c*_1_) for 2,000 generations with six different initial conditions. The initial conditions are in pairs, where the minor allele at either both, one, or neither of the two loci has low initial frequency. Then, for each pair, one initial condition exhibits LD and the other does not. The haplotype frequencies for each of these conditions are listed in the order (*x*_11_, *x*_12_, *x*_21_, *x*_22_), with the first digit in the subscript referring to the allele at the first locus and the second digit to the allele at the second locus. The specific marginal allele frequencies and haplotype frequencies are given in Table 1.

**TABLE 1.** The six initial conditions x^(0)^ used for the simulations. We indicate the respective marginal initial frequencies and whether LD is zero (without LD) or not (with LD).

| Marginal Initial Frequencies<br>$(X_{1\cdot}(0), X_{\cdot 1}(0))$ | with LD? | $\mathbf{x}^{(0)} = (x_{11}, x_{12}, x_{21}, x_{22})$ |
| --- | --- | --- |
| (0.5, 0.5) | no | (0.25, 0.25, 0.25, 0.25) |
| (0.5, 0.5) | yes | (0.4, 0.1, 0.1, 0.4) |
| (0.05, 0.5) | no | (0.025, 0.025, 0.475, 0.475) |
| (0.05, 0.5) | yes | (0.05, 0, 0.45, 0.5) |
| (0.05, 0.05) | no | (0.0025, 0.0475, 0.0475, 0.9025) |
| (0.05, 0.05) | yes | (0.05, 0, 0, 0.95) |

**FIGURE 2.**
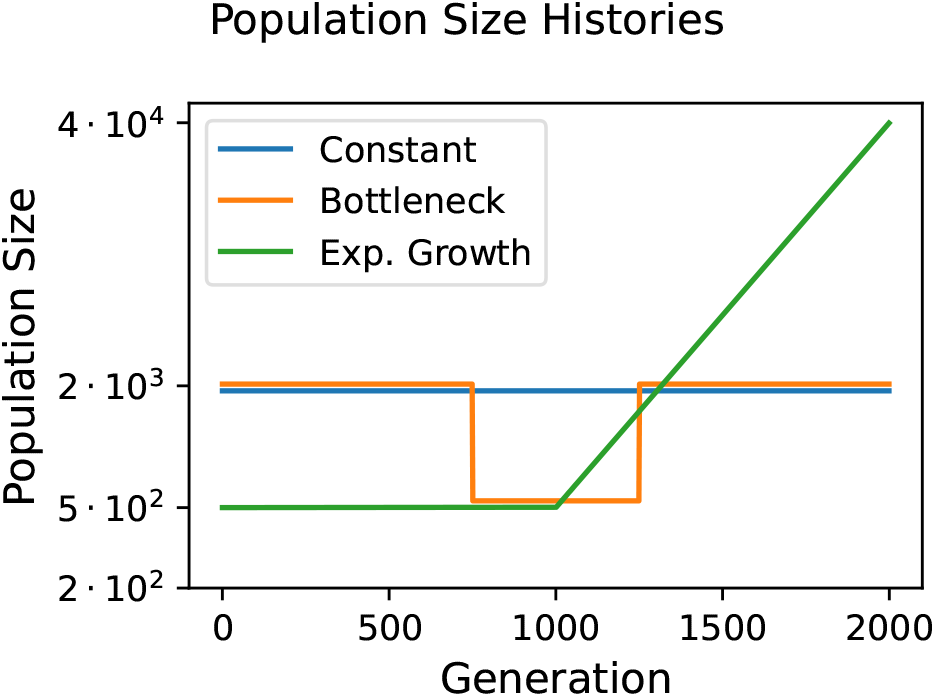
The three population size histories considered in our simulations: constant, bottleneck, and exponential growth. Note that the y-axis is on a logarithmic scale and the size histories are slightly shifted for visibility.

**FIGURE 3.**
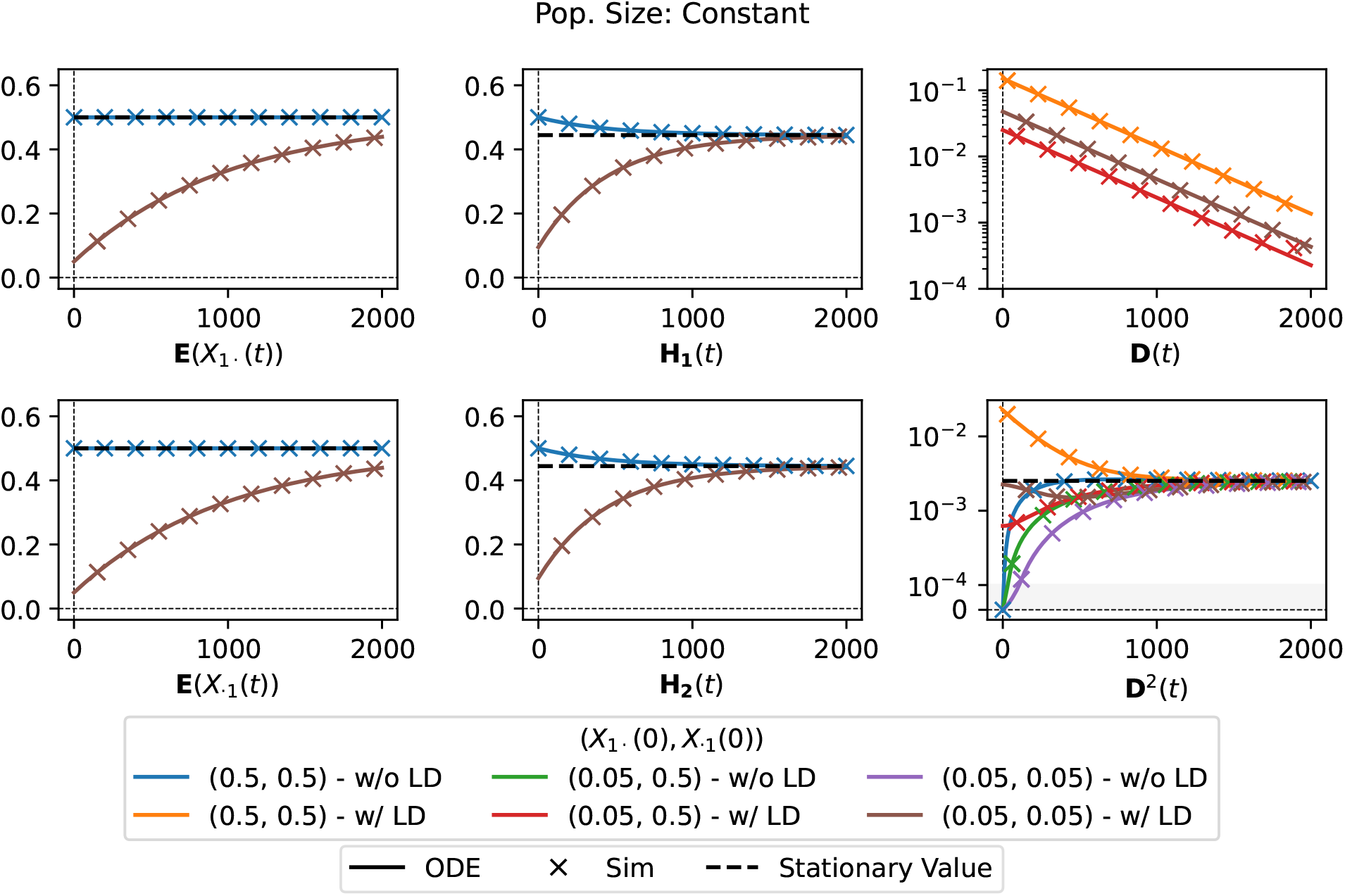
Trajectories of the expected allele frequencies E[*X*_1·_(*t*)] and E[*X*_·1_(*t*)], expected heterozygosities **H**_1_(*t*) and **H**_2_(*t*), and both expected LDs **D**(*t*) and **D**^2^(*t*) under the model of constant population size for different initial conditions. Estimated using 2^18^ simulated replicates and directly computed from the solution of the ODEs for *r* = 10^−4^ and *u* = 5·10^−4^. For E[*X*_1·_(*t*)], E[*X*_·1_(*t*)], **H**_1_(*t*), and **H**_2_(*t*), the six initial conditions have redundancies, so we only plot two. For **D**(*t*), we only plot non-zero trajectories. The gray shading in the plot for **D**^2^(*t*) indicates linear scaling.

**FIGURE 4.**
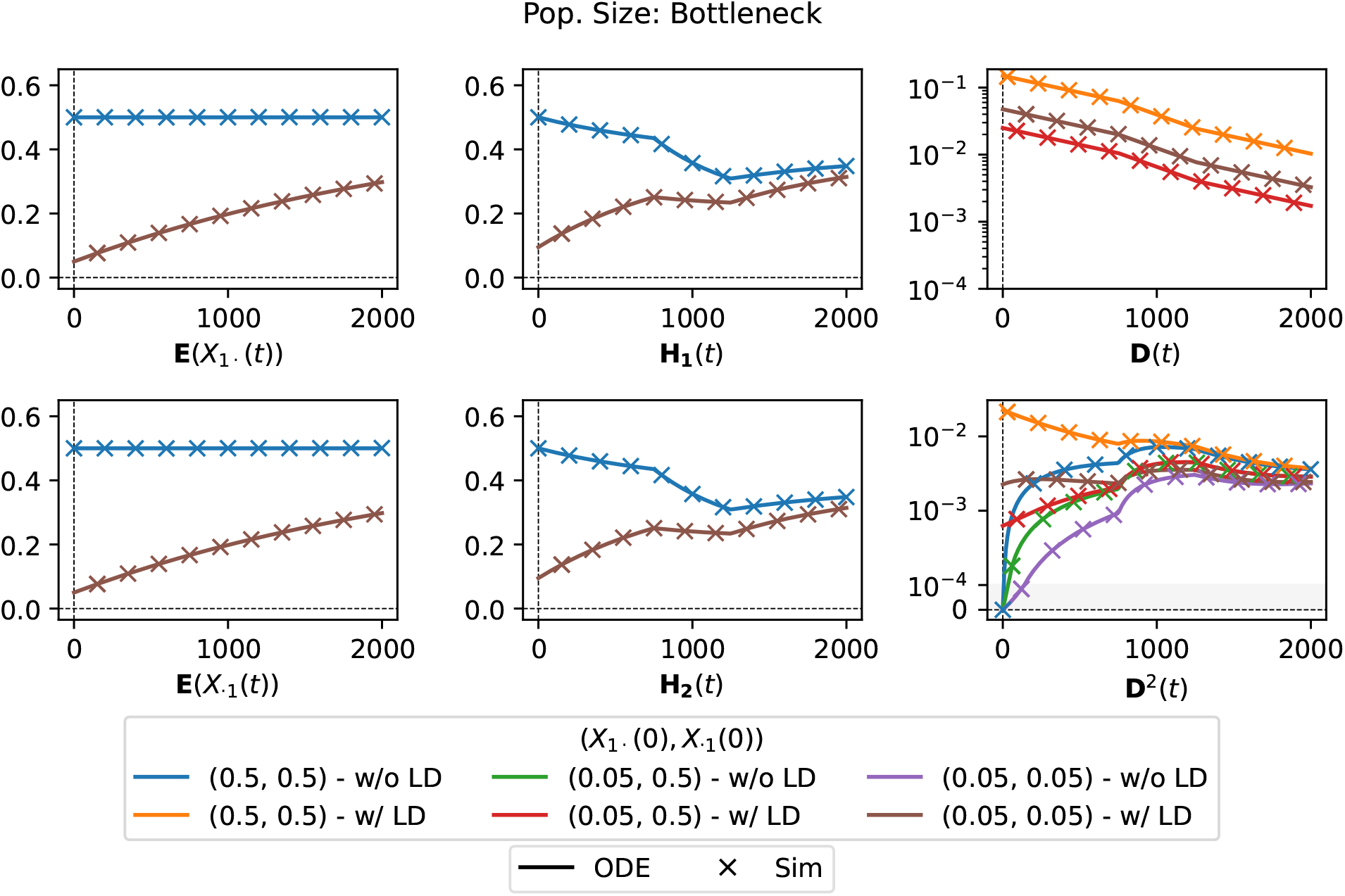
Trajectories of the expected allele frequencies E[*X*_1·_(*t*)] and E[*X*_·1_(*t*)], expected heterozygosities **H**_1_(*t*) and **H**_2_(*t*), and both expected LDs **D**(*t*) and **D**^2^(*t*) under the bottle-neck population size history for different initial conditions. Estimated using 2^18^ simulated replicates and directly computed from the solution of the ODEs for *r* = 10^−4^ and *u* = 2·10^−4^. For E[*X*_1·_(*t*)], E[*X*_·1_(*t*)], **H**_1_(*t*), and **H**_2_(*t*), the six initial conditions have redundancies, so we only plot two. For **D**(*t*), we only plot non-zero trajectories. The gray shading in the plot for **D**^2^(*t*) indicates linear scaling.

**FIGURE 5.**
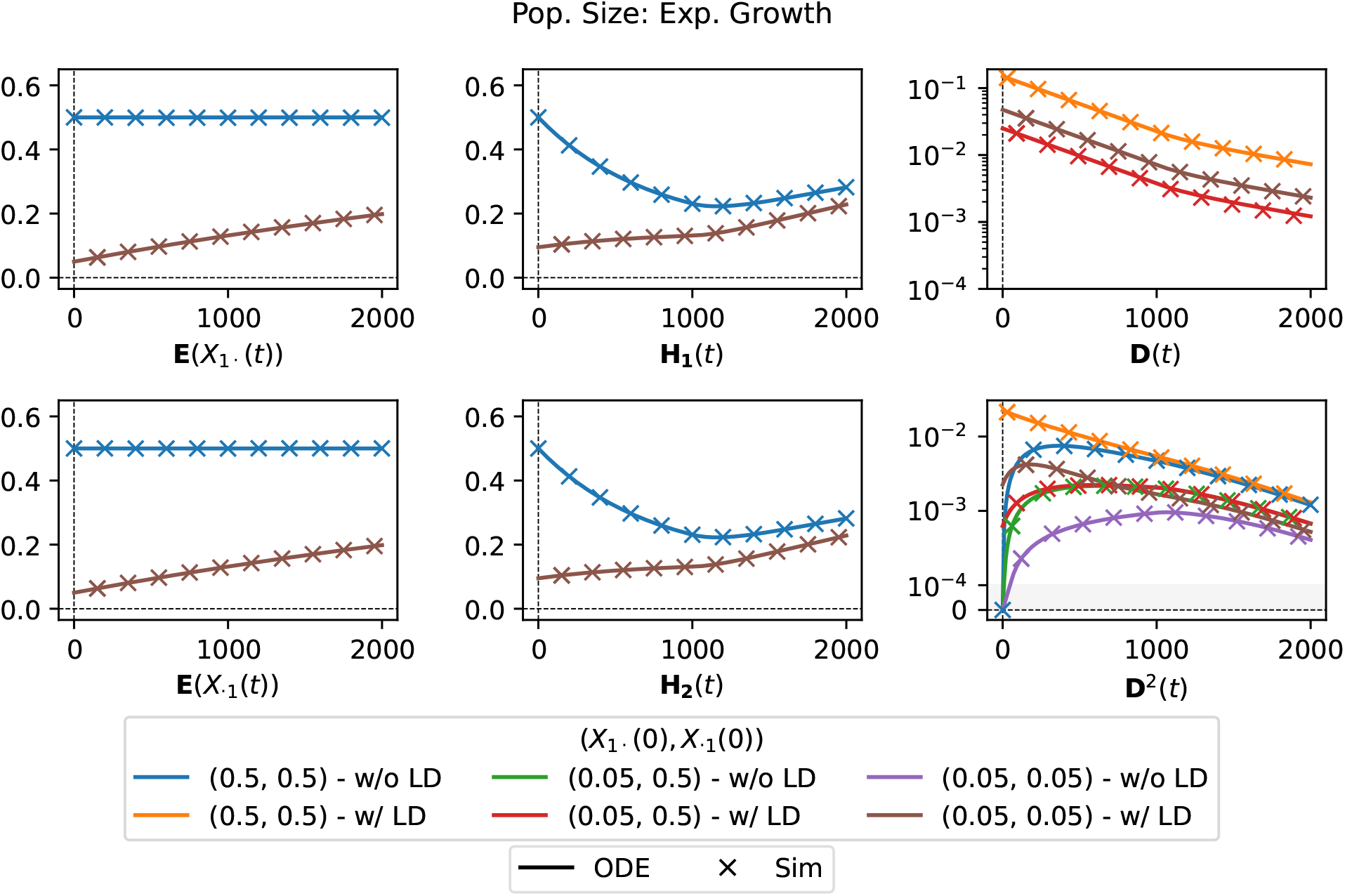
Trajectories of the expected allele frequencies E[*X*_1·_(*t*)] and E[*X*_·1_(*t*)], expected heterozygosities **H**_1_(*t*) and **H**_2_(*t*), and both expected LDs **D**(*t*) and **D**^2^(*t*) for the population with exponential growth under different initial conditions. Estimated using 2^18^ simulated replicates and directly computed from the solution of the ODEs for *r* = 5·10^−4^ and *u* = 10^−4^. For E[*X*_1·_(*t*)], E[*X*_·1_(*t*)], **H**_1_(*t*), and **H**_2_(*t*), the six initial conditions have redundancies, so we only plot two. For **D**(*t*), we only plot non-zero trajectories. The gray shading in the plot for **D**^2^(*t*) indicates linear scaling.

These six conditions are used with three different population size histories: a population with a constant size of *N*_*e*_ = 2000; a population of size *N*_*e*_ = 2000 that experienced a bottleneck of size *N*_*e*_ = 500 between generation 750 and 1,250; and a population initially of size *N*_*e*_ = 500 that starts to grow exponentially at generation 1,000 until it reaches size 40,000 at generation 2,000. Figure 2 shows the population size trajectories for each of these scenarios.

For the simulated trajectories and the solutions of the ODEs, Figures 3, 4, and 5 show expected marginal allele frequencies 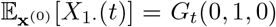 and 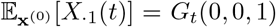, as well as other expected values for statistics of interest: the expected marginal heterozygosity:

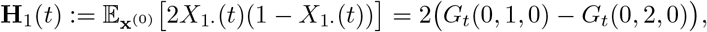

and **H**_2_(*t*) analogously, expected LD:

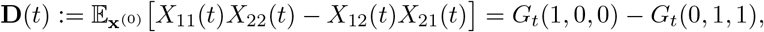

and expected squared LD:

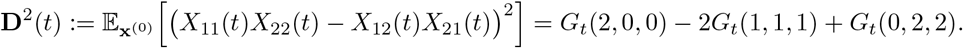

For the simulated trajectories, we simulated 2^18^ = 262, 144 replicates and estimate the expected values by averaging the respective quantities over the replicates. For the ODEs, these expected values are simply linear combinations of the respective moments *G*_*t*_(*a*_11_, *b*_1_, *c*_1_).

This comparison to simulations serves as validation that the solution to the system of ODEs for the closed moments *G*_*t*_(*a*_11_, *b*_1_, *c*_1_) accurately recapitulates the dynamics of the expected allele frequencies, heterozygosities and LD statistics. Our system of ODEs matches simulations in scenarios with a variety of population size histories and initial conditions, which allows efficient computation of these and other statistics in scenarios of interest. Specifically in the scenarios investigated, we observe that in the case of constant population size, the expected values approach their respective values at stationarity over time, whereas in the bottleneck and exponential growth scenario, the trajectories change shape at times when the population size changes. As has been observed previously (Hill and Robertson, 1968), expected LD decreases monotonically to zero, whereas expected squared LD, even if initially at zero, converges to a non-zero stationary value

### 4.2. Two-locus site-frequency-spectrum

As we will detail below, we can compute the standard moments of the haplotype frequencies *M*_*t*_(**n**) from the closed moments *G*_*t*_(*a*_11_, *b*_1_, *c*_1_). However, matching the standard moments or the closed moments to data requires being able to assess phased haplotypes. In many population genetic applications, phased haplotype data is not readily available. We thus aim at using our system of ODEs to derive quantities that can be assessed from just the allele frequencies in the observed data. Such a quantity is given by the two-locus site-frequency spectrum (SFS), 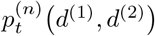. For a given sample size *n*, the *d*^(1)^, *d*^(2)^ -entries of the two-locus SFS with *d*^(1)^ ∈ {0, 1, …, *n*} and *d*^(2)^ ∈ {0, 1, …, *n*} are given by the probability that, in a sample of size *n*, we observe both *d*^(1)^ alleles of type 1 ∈ *E*^(1)^ at the first locus, and *d*^(2)^ alleles of type 1 ∈ *E*^(2)^ at the second locus. In terms of the standard moments *M*_*t*_(*n*) of the Wright-Fisher diffusion, these probabilities are given by

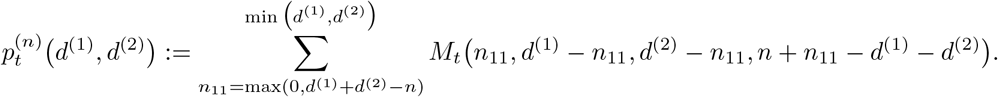

To compute the two-locus SFS, we thus need to compute the standard moments *M*_*t*_(**n**) from the closed moments *G*_*t*_(**a, b, c**). To this end, in Appendix A, we derive a bijection between the closed moments and the standard moments *M*_*t*_(**n**) with *n*_22_ = 0. This relation can be written in the form of a matrix equation

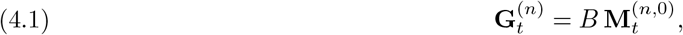

where 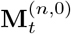 is the vector of all moments *M*_*t*_(**n**) with |**n**| = *n* and 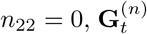 is the vector of all moments *G*_*t*_(*a*_11_, *b*_1_, *c*_1_) with *a*_11_ +*b*_1_ +*c*_1_ = *n*, and *B* is the matrix corresponding to the bijection. In order to compute the two-locus SFS from the closed moments *G*_*t*_(*a*_11_, *b*_1_, *c*_1_), we rely on this matrix equation and the following lemma, which we prove in Appendix A:

#### Lemma 4.1

*With* 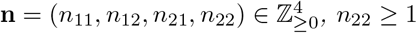, *and* 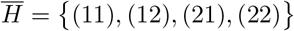, *the relation*

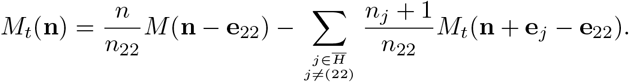

*holds*.

Lemma 4.1 can be used to reduce the last index in the standard moments, and thus we can write any moment *M*_*t*_(**n**) as a linear combination of moments of the form *M*_*t*_(**n**^′^) with 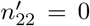. We combine this observation with the bijection (4.1) and define Algorithm 4.1 to compute 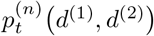 from the closed moments *G*_*t*_(*a*_11_, *b*_1_, *c*_1_). We provide associated proofs and further discussion regarding Algorithm 4.1 in Appendix A.

#### Algorithm 4.1

Compute 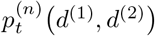 using *G*_*t*_(*a*_11_, *b*_1_, *c*_1_)

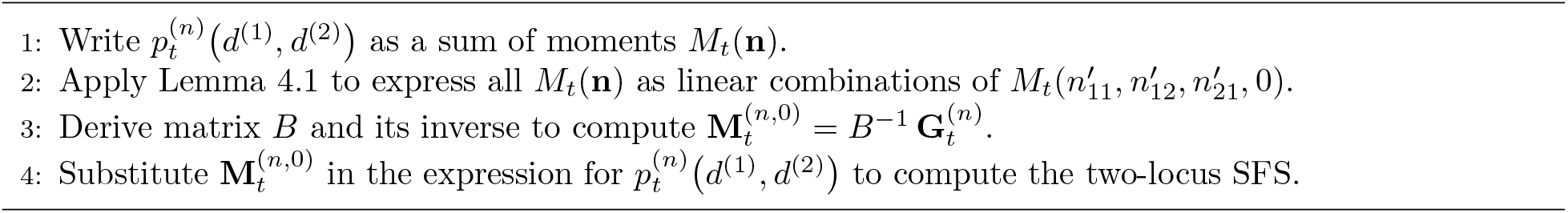

In Figures 6, 7, and 8 we present the temporal dynamics of 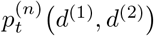 computed using Algorithm 4.1 under the three different population size histories for certain pairs *d*^(1)^, *d*^(2)^ . Here we used different mutation rates, recombination rate *r* = 10^−4^, and initial condition (*X*_1_ (*t*), *X*_1_ (*t*)) = (0.05, 0.5) with LD. For each population size history, these dynamics are shown alongside a heatmap showing all entries of the two-locus SFS at generation 2,000. In this section, all two-locus SFS are computed for a sample of size *n* = 8. We compare the trajectories of 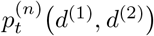 computed using our system of ODEs with estimates from simulations and observe that the ODEs are able to accurately capture the dynamics. For the bottleneck or exponential growth scenario, the dynamics exhibit inflection points at the time of the population size changes.

**FIGURE 6.**
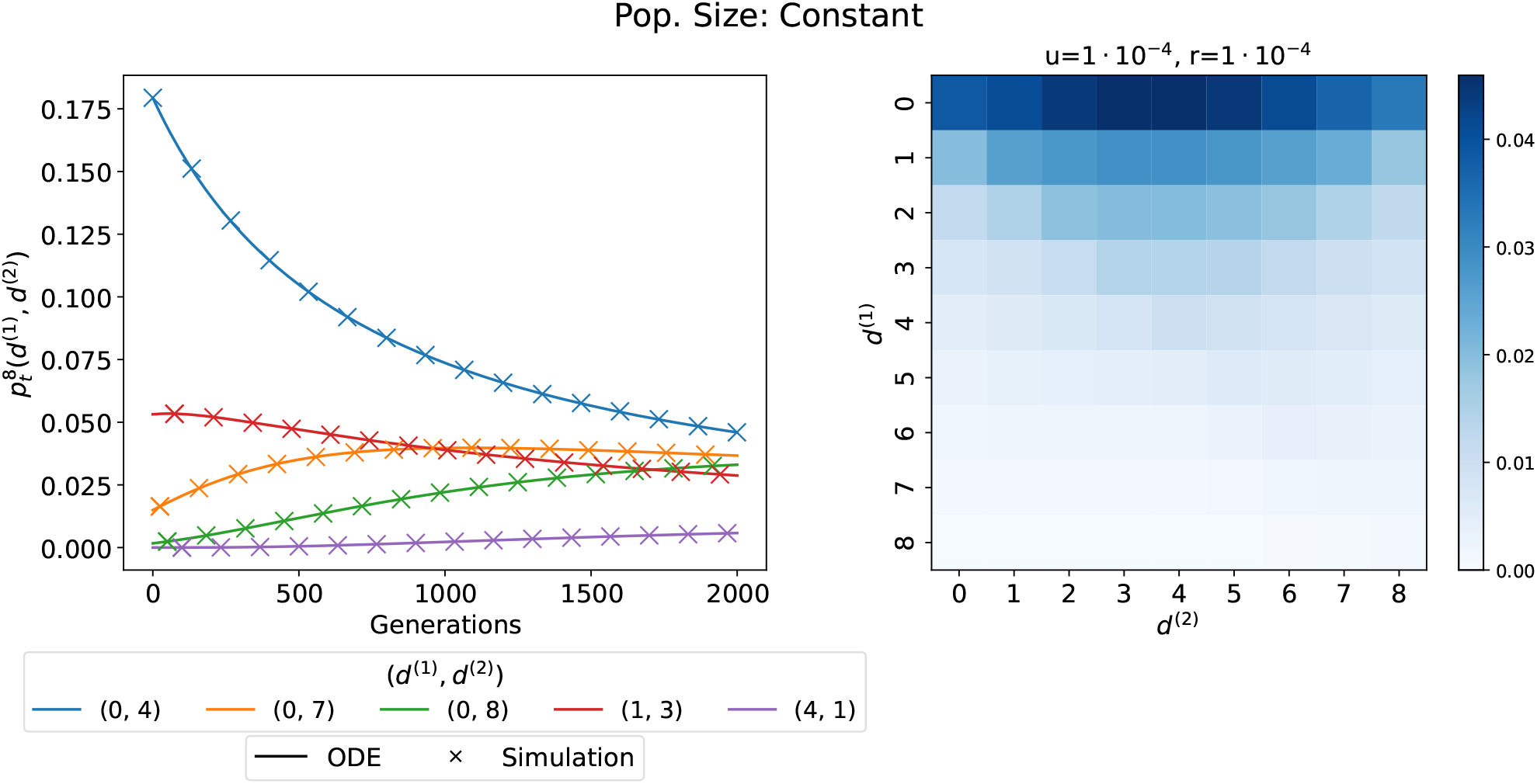
Left panel shows the dynamics of select entries of the two-locus SFS 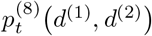 computed from solutions of the ODEs or estimated from 2^18^ simulated replicates. Right panel shows the full SFS at generation 2,000 computed from the ODEs. Constant population size scenario with *r* = 10^−4^, *u* = 10^−4^, and initial condition *X*_1·_(*t*), *X*_·1_(*t*) = (0.05, 0.5) with LD.

**FIGURE 7.**
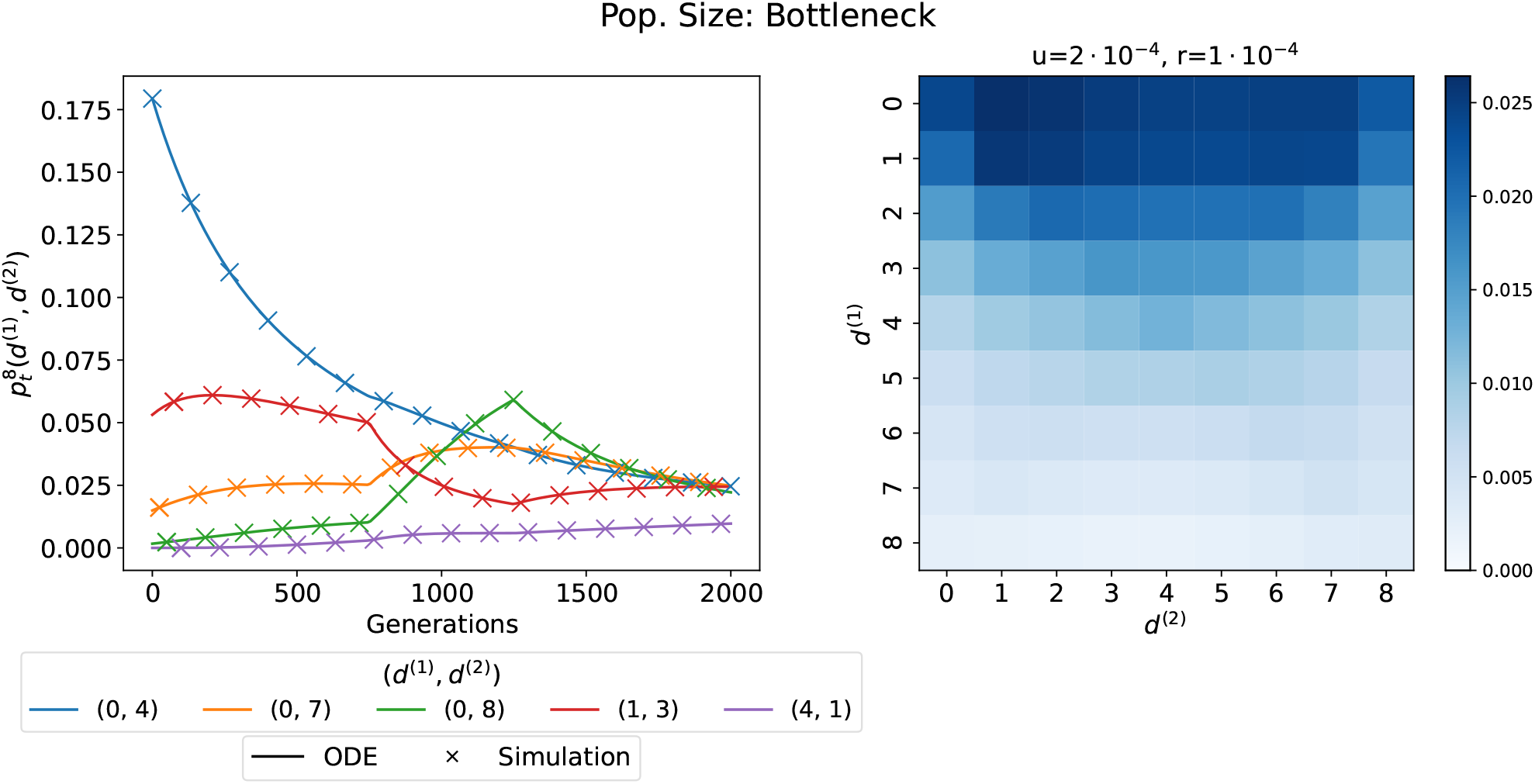
Left panel shows the dynamics of select entries of the two-locus SFS 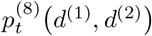 computed from solutions of the ODEs or estimated from 2^18^ simulated replicates. Right panel shows the full SFS at generation 2,000 computed from the ODEs. Bottle-neck scenario with *r* = 10^−4^, *u* = 2 10^−4^, and initial condition *X*_1·_(*t*), *X*_·1_(*t*) = (0.05, 0.5) with LD.

**FIGURE 8.**
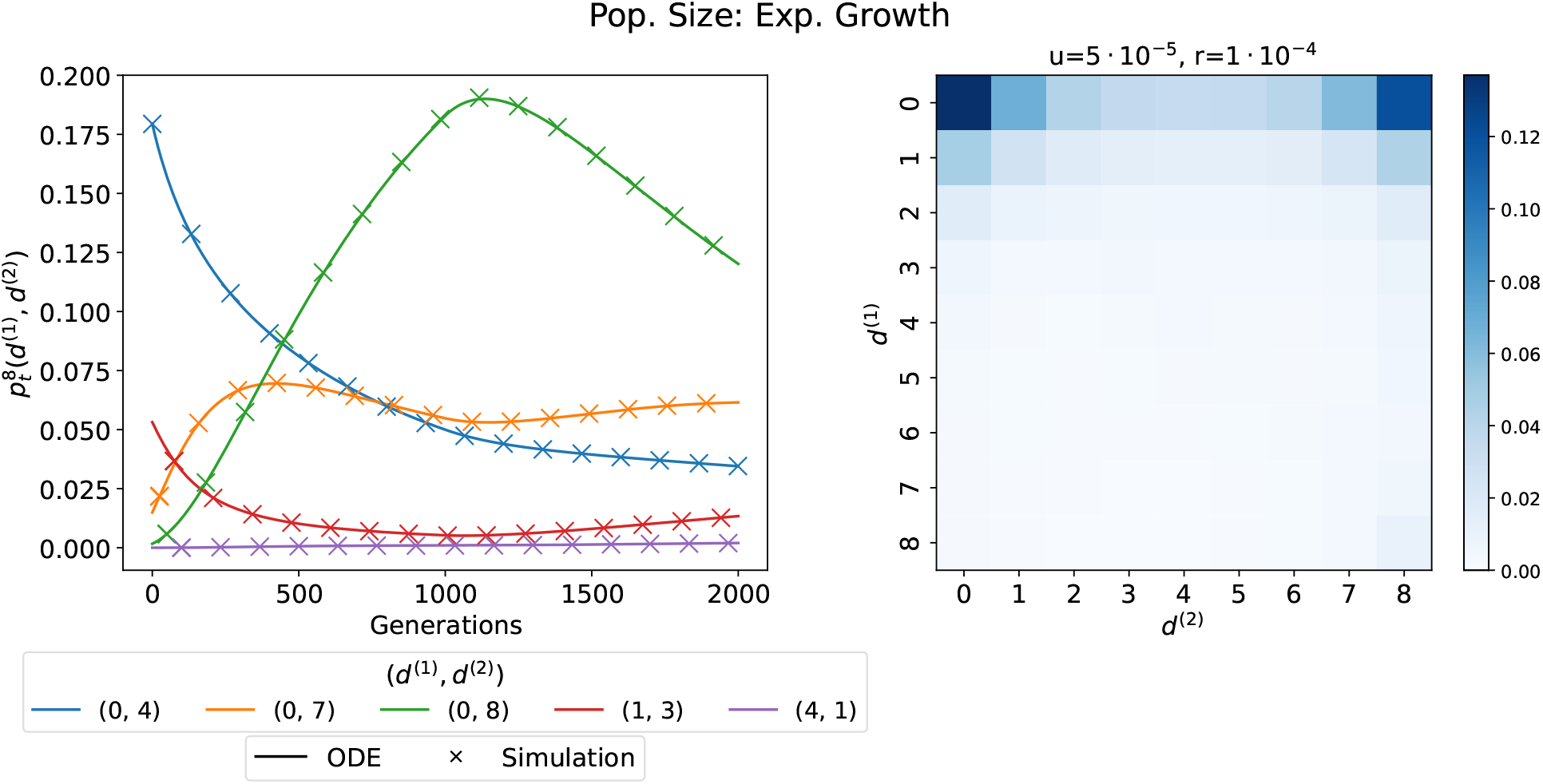
Left panel shows the dynamics of select entries of the two-locus SFS 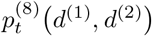 computed from solutions of the ODEs or estimated from 2^18^ simulated replicates. Right panel shows the full SFS at generation 2,000 computed from the ODEs. Scen ario with expo nentially growing population size, *r* = 10^−4^, *u* = 5·10^−5^, and initial condition *X*_1·_(*t*), *X*_·1_(*t*) = (0.05, 0.5) with LD.

As stated in Corollary 3.11, the closed system of ODEs can be conveniently used to compute moments *G*_*t*_(*a*_11_, *b*_1_, *c*_1_) at stationarity. This in turn allows us to compute the two-locus SFS 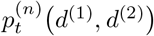 at stationarity. We use this as follows to model genetic variation in natural populations that experienced recent population size changes. In the three demographic models considered here, we initialize the moments at generation 0 to the stationary moments assuming a constant population size equal to the size at generation 0. We can then use the system of ODEs to compute the dynamics for the moments between generation 0 and 2,000, accounting for the population size changes in this period, and ultimately arrive at a two-locus SFS at generation 2,000. Note that in the case of constant population size, if the moments at generation 0 are stationary, then the moments at generation 2,000 are also stationary and equal to those at generation 0.

In Figure 9, we present heatmaps of the two-locus SFS in each demographic scenario, either initialized with the stationary moments, or using the initial condition *X*_1·_(*t*), *X*_·1_(*t*) = (0.05, 0.5) with LD, using mutation rate *u* = 2 · 10^−4^ and recombination rate *r* = 10^−4^. In the cases when we initialize with given frequencies, *d*^(1)^ remains low and *d*^(2)^ intermediate with high probability. As expected, the frequencies are less concentrated under the bottleneck, and more concentrated around low frequencies at the first locus in the case of exponential growth.

**FIGURE 9.**
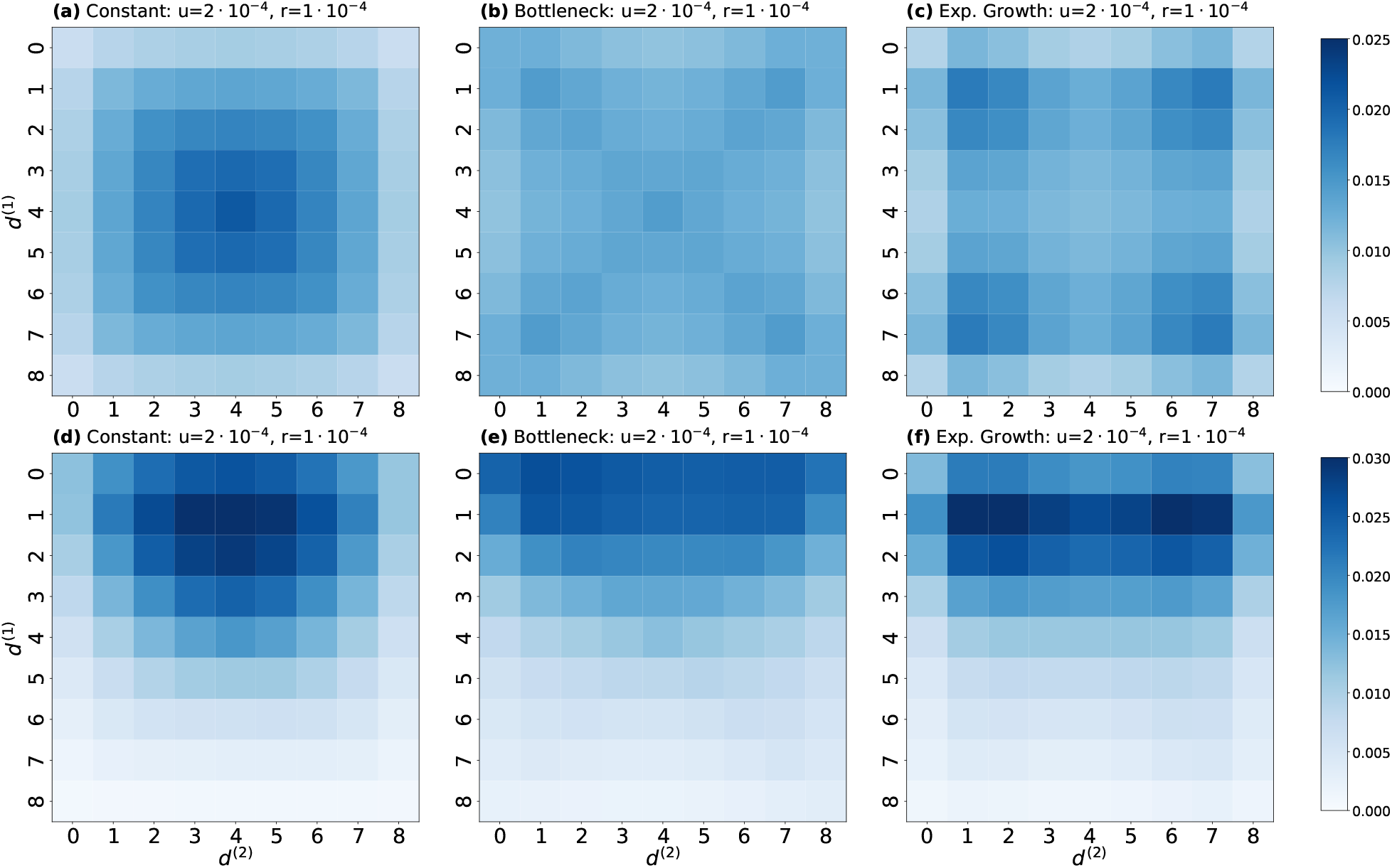
The two-locus SFS 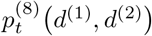 at generation 2,000 computed using the ODEs, under a model of: **(a) & (d)** constant population size, **(b) & (e)** bottleneck, and **(c) & (f)** exponential growth, using *r* = 10^−4^ and *u* = 2 · 10^−4^. The initial conditions are: **(a), (b) & (c)** the stationary moments and **(d), (e) & (f)** *X*_1·_(*t*), *X*_·1_(*t*) = (0.05, 0.5) with LD.

The heatmaps for the two-locus SFS when initialized with the stationary moments exhibit strong symmetries, which are a result of the mutation rates we used in these scenarios: Since we set the mutation rates to the same values at both loci, we have 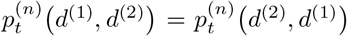. Moreover, from 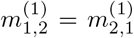 it follows that 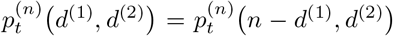. The heatmaps are therefore mirrored along the vertical, horizontal, and diagonal axes, and the 9 × 9 = 81 entries are essentially determined by 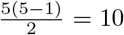 entries. Besides these symmetries, the same patterns as before hold. For the bottleneck, allele counts are more evenly distributed, and in the case of exponential growth, we observe more probability for lower allele counts, since new alleles enter the population at lower frequencies in a larger population and genetic drift is weaker.

In Figures 6, 7, and 8, we demonstrate that the system of ODEs matches the dynamics of the two-locus SFS when compared to forward-in-time simulations of the corresponding Wright-Fisher model. For further validation, we compare solutions obtained from the ODEs to simulations performed with the widely used backward-in-time coalescent simulator `msprime` (Baumdicker et al., 2022) in Figure 10. The first column in Figure 10 shows heatmaps for the two-locus SFS computed using Algorithm 4.1 in the three demographic scenarios with recombination rate *r* = 10^−4^ and mutation rate *u* = 22 · 10^−4^, initialized with the stationary moments.

**FIGURE 10.**
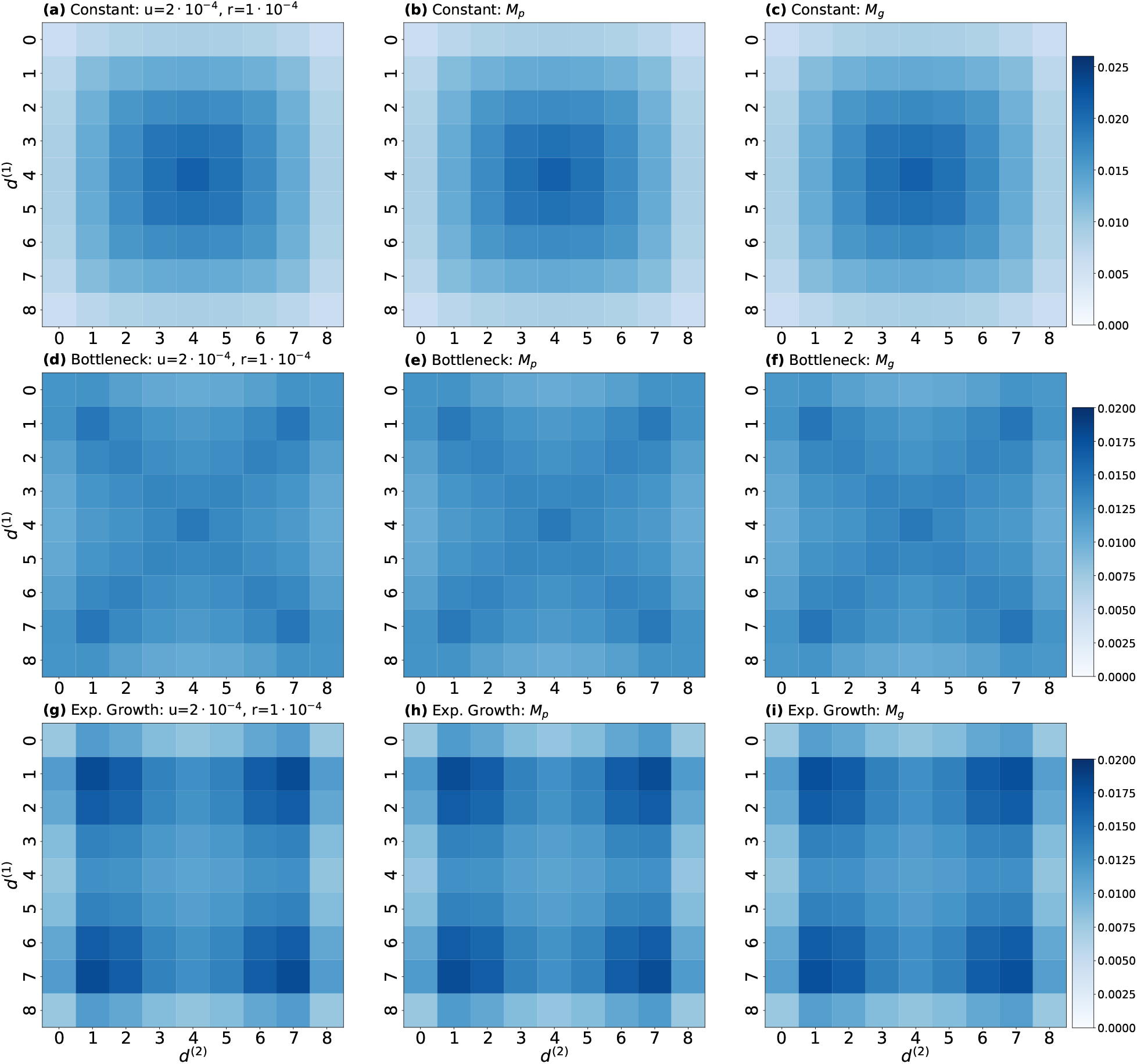
The two-locus SFS 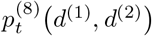 at generation 2,000 under a model of: **(a), (b) & (c)** constant population size, **(d), (e) & (f)** bottleneck, and **(g), (h) & (i)** exponential growth, using *r* = 10^−4^ and *u* = 2 · 10^−4^. The SFS is computed using: **(a), (d) & (g)** the ODEs, **(b), (e) & (h)** pairwise `msprime` simulations *M*_*p*_, and **(c), (f) & (i)** genomic `msprime` simulations *M*_*g*_.

We implemented two different ways to obtain the two-locus SFS from simulations using `msprime`. The first approach is to simulate a coalescent with exactly two loci separated by the given recombination distance *r*, and where the mutation rate at each locus is given by *u*. We used the `BinaryMutationModel` in `msprime` to implement a recurrent mutation model. It is important to note that in scenarios with recent population size changes, `msprime` implicitly assumes that the population is at equilibrium in the ancient times before the changes, which corresponds to initializing our system of ODEs with the stationary moments. In the `msprime` simulations with exactly two loci, each simulated replicate yields a pair of allele counts *d*^(1)^, *d*^(2)^ . We can thus obtain the probabilities for each entry in the two-locus SFS using Monte Carlo estimation from a large number of replicates. We denote this approach by *M*_*p*_ and show the respective two-locus SFS using 2^20^ replicates in the second column of Figure 10. The number of replicates ensured that the relative error of the Monte Carlo estimates for each entry of the two-locus SFS was below 1%.

Alternatively, rather than simulating a large number of two-locus replicates explicitly, we can simulate one sample of size *n* for a long genomic sequence using `msprime`, and tally all observed pairs of segregating sites to obtain a two-locus SFS from one simulated replicate. This interpretation mirrors how two-locus statistics computed from genomic data have been used to infer demographic parameters in past work (Ragsdale and Gravel, 2019; Novo et al., 2022). In particular, we simulate using a per locus mutation rate of *u* = 2 · 10^−4^, a recombination rate of *r* = 10^−6^ between adjacent sites, and a genome of length 4 10^4^ base pairs. The resulting pairs of segregating sites will be at random recombination distances. To compare the two-locus SFS obtained by solving the ODEs for a fixed recombination rate faithfully to the genomic simulations, we thus have to bin the pairs of segregating sites into bins of similar recombination rates. Using these binned pairs, we then compute Monte Carlo estimates for the entries of the two-locus SFS for a given recombination distance. To compare to the ODEs solved with *r* = 10^−4^, we group all pairs of segregating sites that have a recombination distance between 75 and 125 together to obtain the SFS. We label this second approach as *M*_*g*_ and present the corresponding two-locus SFS obtained as the average of 64 genomic replicates in the third column of Figure 10. Again, the relative error of the estimates for each entry was below 1%. We observe that the two-locus SFS obtained using the system of ODEs visually match the corresponding SFS obtained from the `msprime` simulations well. In Appendix B, we compute explicit distance and divergence measures between these SFS to provide further support that our ODEs match the simulations well.

In addition to the symmetry demonstrated in Figure 9, we highlight some other features of the two-locus SFS when initialized with stationary moments. When considering two unlinked loci with free recombination between them, the stationary two-locus SFS is given by the product of the probabilities of the marginal SFS, that is,

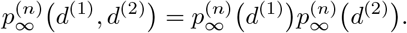

Consequently, the heatmaps are given by the outer product of the marginal probabilities. If recombination is not free, but occurs at a high rate, this independence still essentially holds. However, if the recombination rate is low or zero, the genetic variation at the two loci is not independent, even at stationarity. We found it most convenient to reason about this using the dual process, the coalescent with recombination. If there is low recombination between the two loci, the genealogies at both loci are nearly identical. Mutations are then superimposed on the genealogies to obtain the allele counts in the sample. The allele counts at both loci are not strongly correlated, since mutation at both loci is by definition independent, but some correlation is present because of the underlying identical genealogies.

To demonstrate this effect, Figures 11 and 12 each display six heatmaps corresponding to two-locus SFS. Both sets have a pair of heatmaps for each population size history scenario, and each scenario has a heatmap with recombination rate *r*_linked_ = 0 (fully linked) and recombination rate *r*_free_ = 0.5 (free recombination). Figure 11 shows these scenarios with a low mutation rate of *u*_low_ = 5 10^−5^, and Figure 12 with a high mutation rate of *u*_high_ = 2 10^−4^. We observe, more clearly in Figure 11, elevated values along the diagonals in Figures 11(a), 11(b), and 11(c), which indicate this dependence, whereas no elevated values are present in the corresponding free recombination cases, indicating independence between the marginals. The same features are present in Figure 12, but visually less prominent.

**FIGURE 11.**
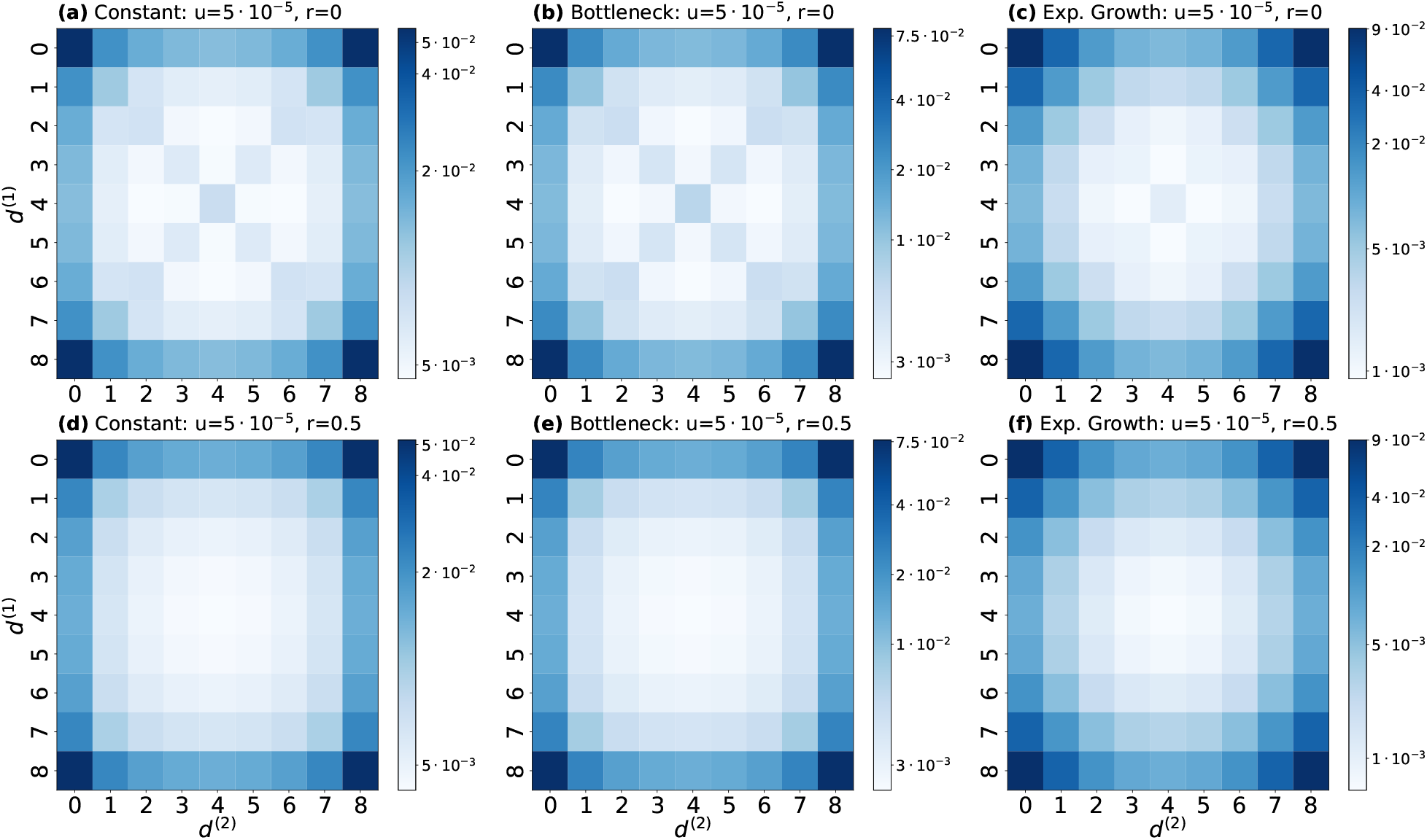
The two-locus SFS 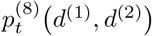 at generation 2,000 initialized from stationarity under a model of: **(a) & (d)** constant population size, **(b) & (e)** bottleneck, and **(c) & (f)** exponential growth. Computed for **(a), (b) & (c)** no recombination *r*_low_ = 0 and **(d), (e) & (f)** free recombination *r*_high_ = 0.5, using a mutation rate of *u* = 5 · 10^−5^.

**FIGURE 12.**
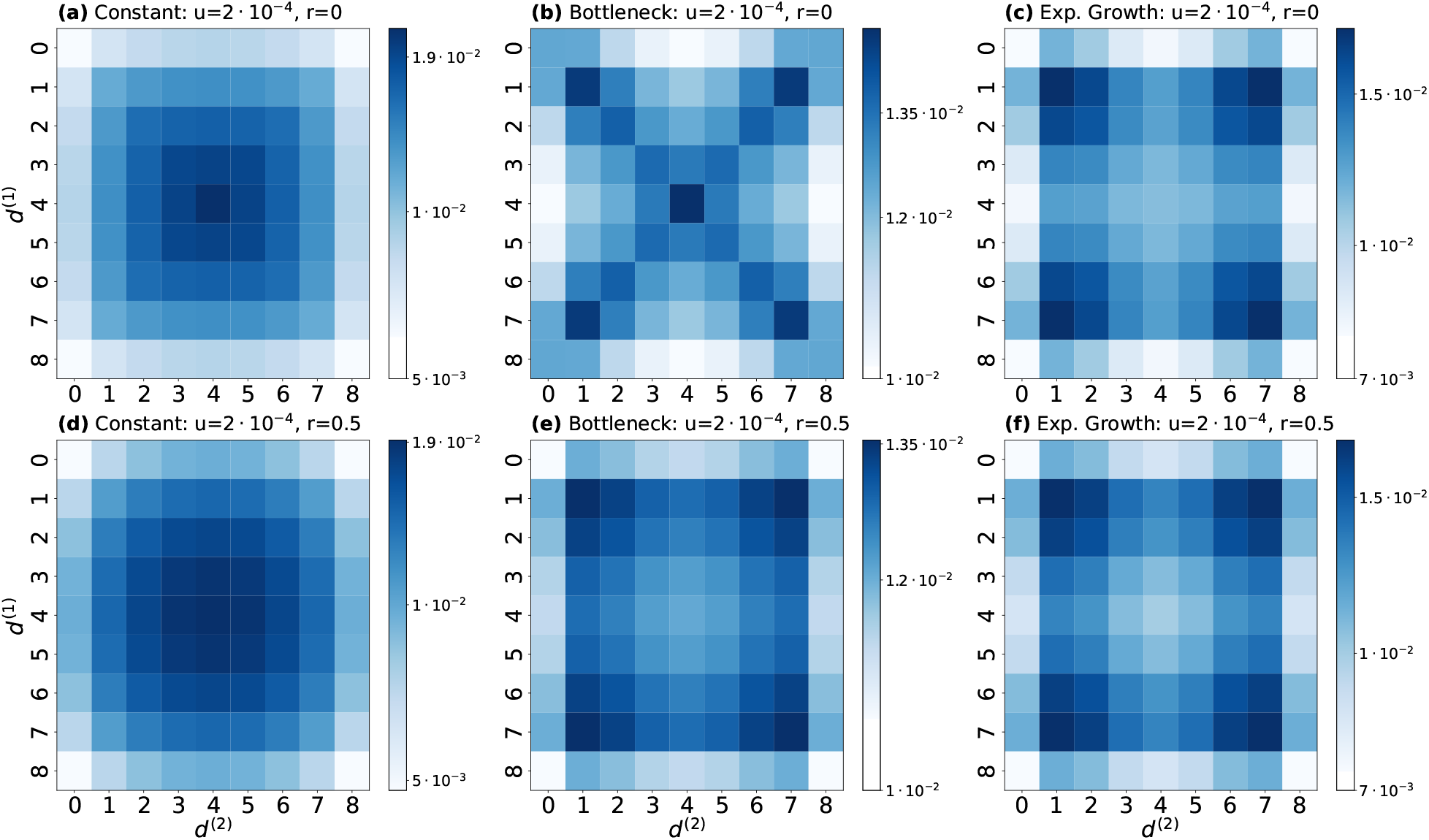
The two-locus SFS 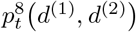 at generation 2,000 initialized from stationarity under a model of: **(a) & (d)** constant population size, **(b) & (e)** bottleneck, and **(c) & (f)** exponential growth. Computed for **(a), (b) & (c)** no recombination *r*_low_ = 0 and **(d), (e) & (f)** free recombination *r*_free_ = 0.5, using a mutation rate of *u* = 2 · 10^−4^.

To quantify this dependence further, we measured the mutual information (MI) between the marginal allele counts. MI is an information theoretic metric derived from the Kullback–Leibler (KL) Divergence for joint distributions, which in turn is the expected logarithm of the likelihood ratio between distributions. MI is the KL Divergence between a joint distribution and the product of its marginal distributions. This measure expresses the reduction in uncertainty in one variable due to the knowledge of another, or the amount of information one variable contains about another. Thus, MI measures the dependence present in a joint distribution relative to the distribution obtained from the product of the marginals. A value of 0 indicates complete independence between the marginal distributions.

In Table 2, we present the MI values corresponding to the different scenarios in Figures 11 and 12. For each demographic history and mutation rate, the MI value for free recombination is multiple orders of magnitude lower than the MI value for recombination rate *r*_linked_ = 0, and numerically indistinguishable from zero. This again demonstrates that the marginal allele counts are independent for free recombination, whereas they are dependent in the fully linked case. In addition, we observe a trend in the fully linked case that dependence is higher for lower mutation rates. Again, this is expected, since higher rates for independent mutation at both sites mask the correlation resulting from the shared genealogy stronger. We also observe trends between scenarios with different population size histories. Populations that experienced a bottleneck exhibit higher MI values than constant populations, while populations experiencing exponential growth exhibit lower MI. This suggests that bottleneck events, regardless of recombination rate or mutation rate, increase dependence between marginal allele frequencies in the population.

**TABLE 2.** Values of the Mutual Information for the two-locus SFS in the different population size scenarios: constant, Bottleneck, and Exponential Growth. Each computed with mutation rates *u*_low_ = 5 · 10^−5^ and *u*_high_ = 2 · 10^−4^, as well as recombination rates *r*_linked_ = 0 and *r*_free_ = 0.5.

|  | Constant |  | Bottleneck |  | Exp. Growth |  |
| --- | --- | --- | --- | --- | --- | --- |
| | $u_{\text{low}}$ | $u_{\text{high}}$ | $u_{\text{low}}$ | $u_{\text{high}}$ | $u_{\text{low}}$ | $u_{\text{high}}$ |
| $r_{\text{linked}}$ | $6.38 \cdot 10^{-3}$ | $1.71 \cdot 10^{-3}$ | $1.13 \cdot 10^{-2}$ | $2.77 \cdot 10^{-3}$ | $1.18 \cdot 10^{-3}$ | $6.22 \cdot 10^{-4}$ |
| $r_{\text{free}}$ | $2.18 \cdot 10^{-10}$ | $8.89 \cdot 10^{-10}$ | $2.27 \cdot 10^{-10}$ | $6.87 \cdot 10^{-10}$ | $4.05 \cdot 10^{-12}$ | $4.16 \cdot 10^{-12}$ |

Of further interest is the change in shape of the two-locus SFS when the mutation rate is increased from *u*_low_ in Figure 11 to *u*_high_ in Figure 12. All six heatmaps exhibit a shift of mass toward the center of the distribution. Again, this is expected, as higher recurrent mutation rates increase the likelihood to observe intermediate allele counts. In the case of the constant population size, we can be more specific. Here, the marginal allele frequencies can be described by a beta-binomial distribution. For low mutation rates this distribution is convex, but as the mutation rate increases it becomes concave. This change is reflected in the joint distribution of the allele counts. Again, in Figures 11 and 12, we demonstrate the that the system of ODEs can capture nuances in the dynamics of allele frequencies for two loci. The system is able to inform situations which would be inefficient to fully explore using simulations.

Lastly, Kamm et al. (2016) present an exact algorithm to compute 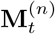 in the two-locus two-allele case under a piecewise constant population size history with complexity *O*(*Tn*^6^). Briefly, this complexity results from evaluating a chain of exponentials of sparse matrices with dimension *O*(*n*^6^) to compute the dynamics of their system, where the variable *T* reflects the computation of the stationary distribution and the sparse matrix exponentials, which can be efficiently implemented using techniques described by Al-Mohy and Higham (2011). They also present an approximate algorithm with complexity *O*(*T n*^3^) and report adequate accuracy. For the complexity of our procedure, the vector 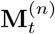 has dimension *O*(*n*^3^), and following Algorithm 4.1, the entries can be computed as linear combinations of entries in 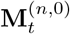 with dimension *O*(*n*^2^), resulting in complexity *O*(*n*^5^). Computing 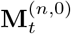 from *G*_*t*_(*a*_11_, *b*_1_, *c*_1_) with *a*_11_ +*b*_1_ +*c*_1_ = *n* using the inverse of the bijection (4.1) has complexity *O*(*n*^4^). To compute *G*_*t*_(*a*_11_, *b*_1_, *c*_1_) with *a*_11_ + *b*_1_ + *c*_1_ = *n*, note that {(*a, b, c*)|*a* + *b* + *c* = *n*} ⊂ *G*(*n*, 0, 0), and thus the entries of the vector 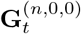 are sufficient to ultimately compute 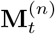. The vector 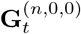 is of dimension *O*(*n*^4^) and under a piece-wise constant population size history the dynamics of this vector can also be computed using a chain of sparse matrix exponentials. Thus, the complexity to compute 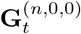 is *O*(*T*^′^*n*^4^), where the variable *T*^′^ reflects computation of the stationary distribution and evaluating the exponentials, similar to Kamm et al. (2016). Thus, the ODEs and algorithm presented here can be used to compute 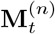 with complexity *O*(*T*^′^*n*^4^ + *n*^5^). We leave a detailed comparison of specific numerical implementations for future work.

## 5. Discussion

In this manuscript, we present a set of moments for the Wright-Fisher diffusion which can be computed using a system of ODEs that is closed under the dynamics of genetic drift, mutation, and recombination. To this end, we transform the standard generator of the diffusion into a new coordinate system, and show that the canonical moments in this new coordinate system yield a closed system. We present extensive simulations to validate these ODEs. We show that the moments computed using the ODEs match the corresponding statistics computed from simulated trajectories. In addition, we show that the sampling probabilities computed using the ODEs align well with the corresponding probabilities obtained from genomic simulations using `msprime`.

The framework presented here can be used to accurately and efficiently compute the dynamics of specific moments, as well as the two-locus haplotype sampling probabilities or the two-locus SFS for contemporary genomic data, under variable population size histories. This naturally suggests applications of the framework to infer populations size histories from genomic data, by matching expected values for these moments to the corresponding statistics obtained from empirical data. As indicated in Section 1, there is a rich history of methods that infer population size histories from patterns of LD (Tenesa et al., 2007; Waples and Do, 2010; Santiago et al., 2020), and our framework would allow these approaches to potentially benefit from higher order LD statistics. Moreover, the two-locus sampling probabilities computed using our framework can be readily substituted into methods for fine-scale recombination rate inference (Auton and McVean, 2007; Chan et al., 2012; Spence and Song, 2019), potentially increasing accuracy and efficiency. Our framework also opens up exciting avenues for possible joint inference of recombination rates and population sizes. In addition, extending the framework to include gene-flow between sub-populations will benefit the development of methods to characterize population structure and migration in natural populations. A straightforward approach to a scenario with *K* sub-population would require the outer product of moments in each, resulting in a vector of |*G*(*a, b, c*)|^*K*^ moments, which grows fairly quickly with the number of populations and the moment-order. Nonetheless, there is potential to increase accuracy when going beyond order 4 (used by Ragsdale and Gravel, 2019), and perhaps room for further improvement using suitable approximations instead of exact moments to increase the efficiency.

The set of moments we present here yields a closed system of ODEs, but it is not unique in this property. Ragsdale and Gravel (2019) also present closed systems in the case of two alleles per locus, which can likely be readily extended to the case of multiple alleles. However, here we aimed at keeping the system of moments minimal and without redundancies, so that numerical solutions are feasible, as well as presenting an approach that can be more readily generalized. As noted in Section 1, Baake and Baake (2003) present certain multi-locus LD-coordinates and use these to provide analytic solutions for the deterministic dynamics. Canonical moments in these coordinates will likely close as well under the full stochastic dynamics including genetic drift. They potentially diagonalize the recombination dynamics, but it is unclear what form the dynamics of genetic drift will take. Future work will determine which set of moments is most efficient in certain scenarios of interest, and perhaps it is possible to derive a set of moments that jointly diagonalizes the dynamics of genetic drift, mutation, and recombination.

We believe that the framework presented here is amenable to several interesting future extensions. First, we use the recurrent mutation model in the work presented here. Another model that is commonly used to analyze genetic variation is the non-recurrent mutation model (Ragsdale and Gravel, 2019), and it would be natural to include this model in our framework. However, rigorous PDEs for the multi-locus non-recurrent mutation model have not been well characterized in the literature, see for example Evans et al. (2007) for the single-locus case, and extending our approach is thus not straightforward. The non-recurrent mutation model has been introduced in similar frameworks heuristically (Ragsdale and Gravel, 2019; Friedlander and Steinrücken, 2022), but a more formal characterization of the associated PDEs and moments would be of interest.

In addition, we believe that the approach presented here for a two-locus system can be readily extended to an arbitrary number of loci, contrary to a conjecture by Baake and Hustedt (2011) that the multi-locus moment system with genetic drift and recombination cannot be closed. In the present paper we transformed the standard generator in haplotype-frequency coordinates to new coordinates that included the correct number of marginal frequencies. We believe that for an arbitrary number of loci, one would need to identify the right set of higher order marginals that would need to be included. Similarly, the canonical moments in a coordinate system based on the transformation presented by Baake and Baake (2003) for the case without drift will likely also yield a closed system of ODEs. However, Baake and Baake (2003) only present an analytic solution for cross-over recombination using their coordinates, that is, a recombination process where the genetic material on both sides of a given single breakpoint is exchanged. The authors state (Remark after Theorem 2, Baake and Baake, 2003) that their coordinate system does not lead to an analytic solution for other recombination processes, like gene conversion, where genetic material in a given region within the genome is exchanged between two haplotypes. It is conceivable that similar restrictions will hold when applying an approach based on this coordinate system to the moment-closure problem for an arbitrary number of loci. However, Baake and Baake (2016) present a solution for the general recombination dynamics without drift, so an approach based on this solution might lead to a closed system in the case of an arbitrary number of loci with general recombination. It remains an open question whether any specific approach for an arbitrary number of loci is superior, but we believe an extension of the approach presented here using multi-locus marginals to be a promising avenue.

Lastly, deriving an appropriate system of ODEs for a scenario with three loci, where two are neutral and one is under selection is another possible extension with a wide range of applications. Previous work has investigated the deterministic dynamics of LD between two neutral loci in the vicinity of selective sweeps (Stephan et al., 2006), and one interesting result is that LD across a completed sweep is zero, but elevated surrounding the sweep, which has motivated statistics used to scan for genomic signatures of selective sweeps (Kim and Nielsen, 2004). Developing a framework that could describe LD and higher order LD statistics between two neutral loci in the presence of a selected variant under the full stochastic dynamics including variable genetic drift could thus be used to devise improved tests for selective sweeps. While a closed system for the three-locus dynamics under recombination seems possible, it is unlikely that selection can be closed. However, closing the selection dynamics using numerical approximations still has the potential to yield efficient, accurate, and flexible solutions that will enable a more exact characterization of signatures of selection. In addition, it is common to approximate adaptive forces that affect genome-wide variation using two-locus statistics, for example background selection (Murphy et al., 2022; Barroso and Ragsdale, 2026) or polygenic adaptation (Bulmer, 1971; Negm and Veller, 2026), and we believe that adding selection to the framework presented here has the potential to yield more accurate approximations in these scenarios as well.

## Acknowledgements

MS was supported in part by the National Institute of General Medical Sciences (NIGMS) of the National Institutes of Health under award R01GM146051 and by grants from the NSF (DMS-2235451) and Simons Foundation (MPS-NITMB-00005320) to the NSF-Simons National Institute for Theory and Mathematics in Biology (NITMB).

## APPENDIX A Computing THE TWO-LOCUS Site-Frequency-Spectrum

We begin by recalling that we have the following expression for the two-locus SFS in terms of the standard moments:

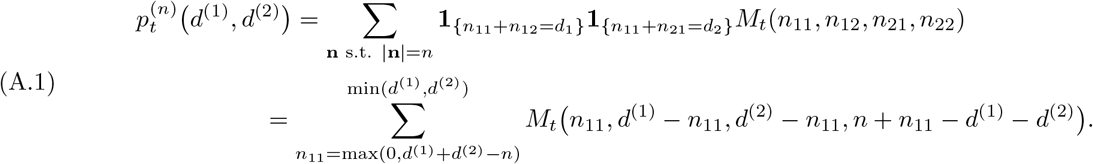

We now need some way to convert equation (A.1), which is expressed in terms of *M*_*t*_(*n*_11_, *n*_12_, *n*_21_, *n*_22_) into an expression in terms of the closed moments *G*_*t*_(*a*_11_, *b*_1_, *c*_1_). We can do so using the following observations.

*Remark* A.1. Recall the definition of the closed moments

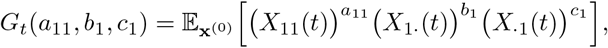

and re-write it in terms of variables used in the moments *M*_*t*_(**n**) by expanding the marginalization

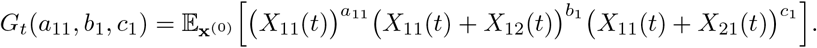

Then, we can expand this further to obtain the following linear combination of *M*_*t*_(**n**):

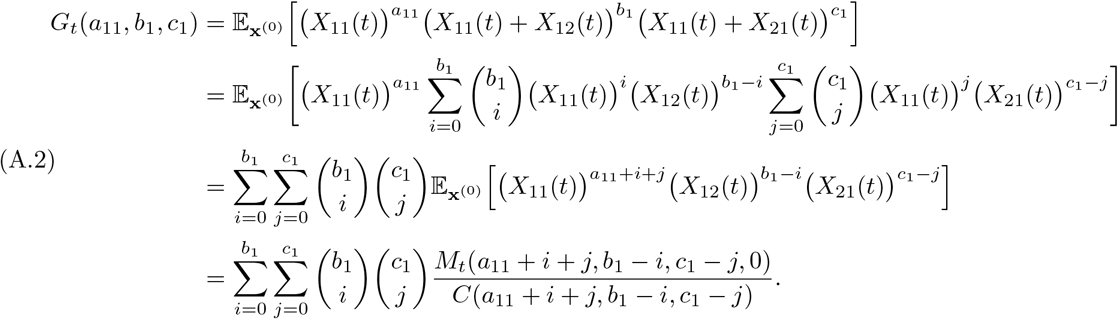

Now, if we denote *n* := *a*_11_ + *b*_1_ + *c*_1_, we observe that all moments in the last expression of equation (A.2) have order *n*. Thus, if we restrict to moments of the form *M*_*t*_(*n*_11_, *n*_12_, *n*_21_, 0), we can use equation (A.2) and arrange the coordinates appropriately to define an invertible triangular matrix *B*, used in Algorithm 4.1, such that

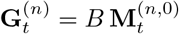

holds. Note that *B* defines a bijection between

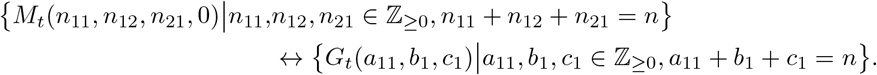

This remark shows that in order to rewrite equation (A.1) in terms of the moments *G*_*t*_(·), it is sufficient to re-write it in terms of moments *M*_*t*_(*n*_11_, *n*_12_, *n*_21_, 0). To achieve this, we employ Lemma 4.1, for which we provide a proof at the end of this section. We apply the relation in Lemma 4.1 to each term of the linear combination in equation (A.1) iteratively until it is written as a linear combination of moments *M*_*t*_(*n*_11_, *n*_12_, *n*_21_, 0). With all the terms of our expansion in this form, Remark A.1 provides a bijection that can be used to compute these moments from the respective closed moments *G*_*t*_(·). The exact bijection can be derived numerically, written as a matrix equation, and inverted to obtain 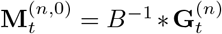. Together, these steps provide a procedure (Algorithm 4.1) to compute the two-locus SFS 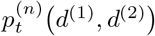 from the closed moments *G*_*t*_(·), which can be obtained from the closed system of ODEs.

*Proof of Lemma 4.1*. First, using the fact that the haplotype frequencies sum to 1, we obtain:

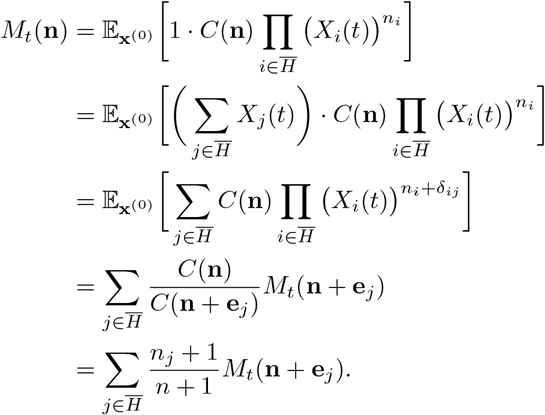

We can then re-write this relation as follows:

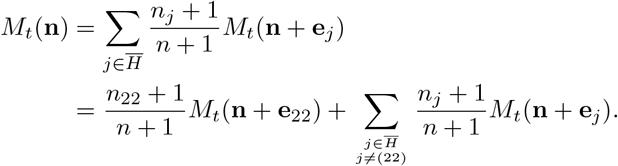

If *n*_22_ *>* 0, this allows us to write:

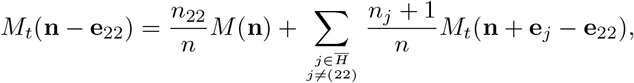

which yields the relation

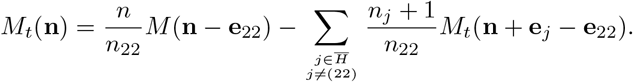

## APPENDIX B. Distance AND DIVERGENCE MEASURES BETWEEN ODE SOLUTIONS AND SIMULATIONS

Figure 10 shows that the two-locus SFS obtained using our ODEs visually match the corresponding SFS obtained from the `msprime` simulations well. Here, we compute explicit distance and divergence measures to support this. Since the two-locus SFS is a probability distribution, we can quantify this match further by computing the divergence between pairs of distributions in two ways: Jensen-Shannon Divergence (JSD) and Total Variation Distance (TVD). JSD is a measure of the difference in information between two distributions. It is a symmetrized and bounded version of the Kullback-Leibler (KL) Divergence. The latter measures the average extra information needed to transform one distribution into the other. JSD is bounded by 0 and 1, where a measure of 0 indicates that two distributions are identical and 1 indicates complete mismatch.

Similarly, TVD is the maximum probability with which any test can distinguish between two distributions, and is also bounded by 0 and 1, with 0 indicating identical distributions. In Table B.1, we present the JSD and TVD values between the two-locus SFS computed using our ODEs and each `msprime` approach as well as between the two `msprime` approaches for validation. From these values, we confirm that both simulations agree with one another and with the distribution computed from the ODEs using Algorithm 4.1. Thus, we affirm the ability of our ODEs to recapitulate the dynamics and stationary distribution of the two-locus system across a variety of scenarios.

**TABLE B1.**
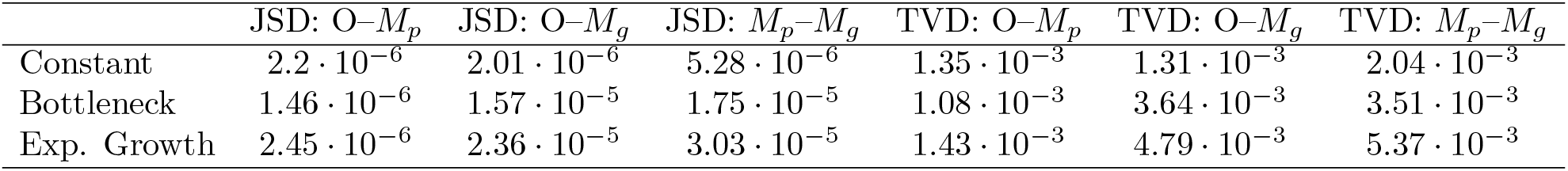
Values of the Jensen-Shannon Divergence (JSD) and Total Variation Distance (TVD) between two-locus SFS obtained from different approaches: O ≡ ODE solutions. *M*_*p*_ ≡ pairwise `msprime` simulations, and *M*_*g*_ ≡ genomic `msprime` simulations.

| | JSD: O-$M_p$ | JSD: O-$M_g$ | JSD: $M_p$-$M_g$ | TVD: O-$M_p$ | TVD: O-$M_g$ | TVD: $M_p$-$M_g$ |
| --- | --- | --- | --- | --- | --- | --- |
| Constant | $2.2 \cdot 10^{-6}$ | $2.01 \cdot 10^{-6}$ | $5.28 \cdot 10^{-6}$ | $1.35 \cdot 10^{-3}$ | $1.31 \cdot 10^{-3}$ | $2.04 \cdot 10^{-3}$ |
| Bottleneck | $1.46 \cdot 10^{-6}$ | $1.57 \cdot 10^{-5}$ | $1.75 \cdot 10^{-5}$ | $1.08 \cdot 10^{-3}$ | $3.64 \cdot 10^{-3}$ | $3.51 \cdot 10^{-3}$ |
| Exp. Growth | $2.45 \cdot 10^{-6}$ | $2.36 \cdot 10^{-5}$ | $3.03 \cdot 10^{-5}$ | $1.43 \cdot 10^{-3}$ | $4.79 \cdot 10^{-3}$ | $5.37 \cdot 10^{-3}$ |

## Notes

https://github.com/steinrue/two_locus_closure

### Competing Interest Statement

The authors have declared no competing interest.

### Summary of Updates

Main text revised/restructured. Some Figures revised.

