## Supplemental Material for "General moment closure for the neutral two-locus wright-fisher dynamics"

### S.1. Proofs of theorems in the main text

S.1.1. **Differential operators in coordinates z.** Recall from equation (3.1) in the main text that

$$\frac{\partial}{\partial x_{ij}} = \begin{cases} \frac{\partial}{\partial z_{ij}} + \frac{\partial}{\partial z_{i*}} + \frac{\partial}{\partial z_{*j}} & \text{if } 1 \le i \le K-1 \text{ and } 1 \le j \le L-1 \\ \frac{\partial}{\partial z_{i*}} & \text{if } 1 \le i \le K-1 \text{ and } j = L \\ \frac{\partial}{\partial z_{*j}} & \text{if } i = K \text{ and } 1 \le j \le J-1. \end{cases} \tag{S.1.1}$$

holds. To obtain the differential operators in the $\mathbf{z}$ coordinates, we substitute these expressions into the differential operators in the $\mathbf{x}$ coordinates.

*Proof of Lemma 3.1 in the main text.* We can rewrite the drift operator defined in equation (2.2) in the main text using the total derivative applied to the chain rule. First, we replace the term $\frac{\partial}{\partial x_{ij}}$ using relation (S.1.1):

$$\begin{aligned} &\frac{1}{4N_e(t)} \sum_{i=1}^{K-1} \sum_{j=1}^{L-1} \Big[ \sum_{k=1}^{K} \sum_{l=1}^{L} \mathbf{1}_{\{k \neq K \text{ or } l \neq L\}} x_{ij} (\delta_{ik}\delta_{jl} - x_{kl}) \frac{\partial}{\partial x_{kl}} \Big] \Big( \frac{\partial}{\partial z_{ij}} + \frac{\partial}{\partial z_{i*}} + \frac{\partial}{\partial z_{*j}} \Big) \\ &+ \frac{1}{4N_e(t)} \sum_{i=1}^{K-1} \Big[ \sum_{k=1}^{K} \sum_{l=1}^{L} \mathbf{1}_{\{k \neq K \text{ or } l \neq L\}} x_{iL} (\delta_{ik}\delta_{Ll} - x_{kl}) \frac{\partial}{\partial x_{kl}} \Big] \frac{\partial}{\partial z_{i*}} \\ &+ \frac{1}{4N_e(t)} \sum_{j=1}^{L-1} \Big[ \sum_{k=1}^{K} \sum_{l=1}^{L} \mathbf{1}_{\{k \neq K \text{ or } l \neq L\}} x_{Kj} (\delta_{Kk}\delta_{jl} - x_{kl}) \frac{\partial}{\partial x_{kl}} \Big] \frac{\partial}{\partial z_{*j}}. \end{aligned}$$

We can now substitute relation (S.1.1) for $\frac{\partial}{\partial x_{kl}}$ as well to obtain

$$\begin{aligned} \sum_{k=1}^{K} \sum_{l=1}^{L} \mathbf{1}_{\{k \neq K \text{ or } l \neq L\}} x_{ij} (\delta_{ik}\delta_{jl} - x_{kl}) \frac{\partial}{\partial x_{kl}} &= \sum_{k=1}^{K-1} \sum_{L=1}^{L-1} x_{ij} (\delta_{ik}\delta_{jl} - x_{kl}) \Big( \frac{\partial}{\partial z_{kl}} + \frac{\partial}{\partial z_{k*}} + \frac{\partial}{\partial z_{*l}} \Big) \\ &+ \sum_{k=1}^{K-1} x_{ij} (\delta_{ik}\delta_{jL} - x_{kL}) \frac{\partial}{\partial z_{k*}} + \sum_{l=1}^{L-1} x_{ij} (\delta_{iK}\delta_{jl} - x_{Kl}) \frac{\partial}{\partial z_{*l}}; \end{aligned}$$

---

[1]Program in Biophysics, Harvard University, Boston, Massachusetts, USA.
[2]Department of Ecology and Evolution, University of Chicago, Chicago, Illinois, USA.
[3]Committee on Computational and Applied Mathematics, University of Chicago, Chicago, Illinois, USA.
[4]Department of Human Genetics, University of Chicago, Chicago, Illinois, USA.
[5]National Institute for Theory and Mathematics in Biology, Northwestern University and University of Chicago, Chicago, Illinois, USA.
[*]Contributed equally.
*Date*: August 28, 2026.

thus, combining these two expansions gives the full drift operator

$$\frac{1}{4N_e(t)}\sum_{i=1}^{K-1}\sum_{j=1}^{L-1}\Bigg[\sum_{k=1}^{K-1}\sum_{l=1}^{L-1}x_{ij}(\delta_{ik}\delta_{jl}-x_{kl})(\frac{\partial^2}{\partial z_{ij}z_{kl}}+\frac{\partial^2}{\partial z_{i*}z_{k*}}+\frac{\partial^2}{\partial z_{*j}z_{*l}}+\frac{\partial^2}{\partial z_{ij}z_{k*}}+\frac{\partial^2}{\partial z_{ij}z_{*l}}$$
$$+\frac{\partial^2}{\partial z_{i*}z_{kl}}+\frac{\partial^2}{\partial z_{i*}z_{*l}}+\frac{\partial^2}{\partial z_{*j}z_{kl}}+\frac{\partial^2}{\partial z_{*j}z_{k*}})$$
$$+\sum_{k=1}^{K-1}x_{ij}(\delta_{ik}\delta_{jL}-x_{kL})\frac{\partial}{\partial z_{k*}}(\frac{\partial}{\partial z_{ij}}+\frac{\partial}{\partial z_{i*}}+\frac{\partial}{\partial z_{*j}})$$
$$+\sum_{l=1}^{L-1}x_{ij}(\delta_{iK}\delta_{jl}-x_{Kl})\frac{\partial}{\partial z_{*l}}(\frac{\partial}{\partial z_{ij}}+\frac{\partial}{\partial z_{i*}}+\frac{\partial}{\partial z_{*j}})\Bigg]$$
$$+\frac{1}{4N_e(t)}\sum_{i=1}^{K-1}\Bigg[\sum_{k=1}^{K-1}\sum_{l=1}^{L-1}x_{iL}(\delta_{ik}\delta_{Ll}-x_{kl})(\frac{\partial}{\partial z_{kl}}+\frac{\partial}{\partial z_{k*}}+\frac{\partial}{\partial z_{*l}})\frac{\partial}{\partial z_{i*}}$$
$$+\sum_{k=1}^{K-1}x_{iL}(\delta_{ik}-x_{kL})\frac{\partial^2}{\partial z_{i*}z_{k*}}+\sum_{l=1}^{L-1}x_{iL}(\delta_{iK}\delta_{Ll}-x_{Kl})\frac{\partial^2}{\partial z_{i*}z_{*l}}\Bigg]$$
$$+\frac{1}{4N_e(t)}\sum_{j=1}^{L-1}\Bigg[\sum_{k=1}^{K-1}\sum_{l=1}^{L-1}x_{Kj}(\delta_{Kk}\delta_{jl}-x_{kl})(\frac{\partial}{\partial z_{kl}}+\frac{\partial}{\partial z_{k*}}+\frac{\partial}{\partial z_{*l}})\frac{\partial}{\partial z_{*j}}$$
$$+\sum_{k=1}^{K-1}x_{Kj}(\delta_{Kk}\delta_{jL}-x_{kL})\frac{\partial^2}{\partial z_{*j}z_{k*}}+\sum_{l=1}^{L-1}x_{Kj}(\delta_{jl}-x_{Kl})\frac{\partial^2}{\partial z_{*j}z_{*l}}\Bigg].$$

Then, we can rearrange by grouping the common partials together, which yields nine terms:

$$\sum_{i=1}^{K-1}\sum_{j=1}^{L-1}\Bigg[\sum_{k=1}^{K-1}\sum_{l=1}^{L-1}x_{ij}(\delta_{ik}\delta_{jl}-x_{kl})\frac{\partial^2}{\partial z_{ij}z_{kl}}\Bigg]+\sum_{i=1}^{K-1}\sum_{j=1}^{L}\Bigg[\sum_{k=1}^{K-1}\sum_{l=1}^{L}x_{ij}(\delta_{ik}\delta_{jl}-x_{kl})\frac{\partial^2}{\partial z_{i*}z_{k*}}\Bigg]$$
$$+\sum_{i=1}^{K}\sum_{j=1}^{L-1}\Bigg[\sum_{k=1}^{K}\sum_{l=1}^{L-1}x_{ij}(\delta_{ik}\delta_{jl}-x_{kl})\frac{\partial^2}{\partial z_{*j}z_{*l}}\Bigg]+\sum_{i=1}^{K-1}\sum_{j=1}^{L-1}\Bigg[\sum_{k=1}^{K-1}\sum_{l=1}^{L}x_{ij}(\delta_{ik}\delta_{jl}-x_{kl})\frac{\partial^2}{\partial z_{ij}z_{k*}}\Bigg]$$
$$(S.1.2)\qquad +\sum_{i=1}^{K-1}\sum_{j=1}^{L-1}\Bigg[\sum_{k=1}^{K}\sum_{l=1}^{L-1}x_{ij}(\delta_{ik}\delta_{jl}-x_{kl})\frac{\partial^2}{\partial z_{ij}z_{*l}}\Bigg]+\sum_{i=1}^{K-1}\sum_{j=1}^{L}\Bigg[\sum_{k=1}^{K-1}\sum_{l=1}^{L-1}x_{ij}(\delta_{ik}\delta_{jl}-x_{kl})\frac{\partial^2}{\partial z_{i*}z_{kl}}\Bigg]$$
$$+\sum_{i=1}^{K-1}\sum_{j=1}^{L}\Bigg[\sum_{k=1}^{K}\sum_{l=1}^{L-1}x_{ij}(\delta_{ik}\delta_{jl}-x_{kl})\frac{\partial^2}{\partial z_{i*}z_{*l}}\Bigg]+\sum_{i=1}^{K}\sum_{j=1}^{L-1}\Bigg[\sum_{k=1}^{K-1}\sum_{l=1}^{L-1}x_{ij}(\delta_{ik}\delta_{jl}-x_{kl})\frac{\partial^2}{\partial z_{*j}z_{kl}}\Bigg]$$
$$+\sum_{i=1}^{K}\sum_{j=1}^{L-1}\Bigg[\sum_{k=1}^{K-1}\sum_{l=1}^{L}x_{ij}(\delta_{ik}\delta_{jl}-x_{kl})\frac{\partial^2}{\partial z_{*j}z_{k*}}\Bigg].$$

For now, we omit the leading $\frac{1}{4N_e(t)}$ for convenience.

In order to write this operator in terms of the coordinates $\mathbf{z}$, we write $x_{ij}=x_{ij}(\mathbf{z})$ and $x_{kl}=x_{kl}(\mathbf{z})$ explicitly as functions of $\mathbf{z}$ and simplify. For the first term in equation (S.1.2), we have that $x_{ij}(\mathbf{z})=z_{ij}$ for $1\le i\le K-1, 1\le j\le L-1$ so we can simply rewrite the first term as

$$\sum_{i=1}^{K-1}\sum_{j=1}^{L-1}\Bigg[\sum_{k=1}^{K-1}\sum_{l=1}^{L-1}z_{ij}(\delta_{ik}\delta_{jl}-z_{kl})\frac{\partial^2}{\partial z_{ij}z_{kl}}\Bigg].$$

Substituting the $x$ variables in the remaining eight terms of equation (S.1.2) is slightly more involved; we will now do so one term and a time. Focusing on the second term in equation (S.1.2),

$$\sum_{i=1}^{K-1}\sum_{j=1}^{L}\left[\sum_{k=1}^{K-1}\sum_{l=1}^{L} x_{ij}(\delta_{ik}\delta_{jl}-x_{kl})\frac{\partial^2}{\partial z_{i*}z_{k*}}\right]=\sum_{i=1}^{K-1}\sum_{j=1}^{L}\left[\sum_{k=1}^{K-1}\sum_{l=1}^{L} x_{ij}(\mathbf{z})(\delta_{ik}\delta_{jl}-x_{kl}(\mathbf{z}))\frac{\partial^2}{\partial z_{i*}z_{k*}}\right],$$

we can rearrange the order of summation and expand the product to obtain

$$\begin{aligned}
&=\sum_{i=1}^{K-1}\sum_{k=1}^{K-1}\left[\sum_{j=1}^{L}\sum_{l=1}^{L} x_{ij}(\mathbf{z})(\delta_{ik}\delta_{jl}-x_{kl}(\mathbf{z}))\right]\frac{\partial^2}{\partial z_{i*}z_{k*}}\\
&=\sum_{i=1}^{K-1}\sum_{k=1}^{K-1}\left[\sum_{j=1}^{L}\sum_{l=1}^{L} x_{ij}(\mathbf{z})\delta_{ik}\delta_{jl}-\sum_{j=1}^{L}\sum_{l=1}^{L} x_{ij}(\mathbf{z})x_{kl}(\mathbf{z})\right]\frac{\partial^2}{\partial z_{i*}z_{k*}}\\
&=\sum_{i=1}^{K-1}\sum_{k=1}^{K-1}\left[\sum_{j=1}^{L} x_{ij}(\mathbf{z})\delta_{ik}-\sum_{j=1}^{L} x_{ij}(\mathbf{z})\sum_{l=1}^{L} x_{kl}(\mathbf{z})\right]\frac{\partial^2}{\partial z_{i*}z_{k*}}\\
&=\sum_{i=1}^{K-1}\sum_{k=1}^{K-1}\left[z_{i*}\delta_{ik}-z_{i*}z_{k*}\right]\frac{\partial^2}{\partial z_{i*}z_{k*}}\\
&=\sum_{i=1}^{K-1}\sum_{k=1}^{K-1}\left[z_{i*}(\delta_{ik}-z_{k*})\right]\frac{\partial^2}{\partial z_{i*}z_{k*}},
\end{aligned}$$

where we used $\sum_{j=1}^{L} x_{ij}(\mathbf{z})=z_{i*}$. Now, focusing on the third term of equation (S.1.2), we similarly expand the product in the summand and rearrange terms, ultimately using $\sum_{i=1}^{K} x_{ij}(\mathbf{z})=z_{*j}$ to arrive at

$$\begin{aligned}
&\sum_{i=1}^{K}\sum_{j=1}^{L-1}\sum_{k=1}^{K}\sum_{l=1}^{L-1} x_{ij}(\mathbf{z})(\delta_{ik}\delta_{jl}-x_{kl}(\mathbf{z}))\frac{\partial^2}{\partial z_{*j}z_{*l}}\\
&\quad=\sum_{j=1}^{L-1}\sum_{l=1}^{L-1}\left[\sum_{i=1}^{K}\sum_{k=1}^{K} x_{ij}(\mathbf{z})\delta_{ik}\delta_{jl}-\sum_{i=1}^{K}\sum_{k=1}^{K} x_{ij}(\mathbf{z})x_{kl}(\mathbf{z})\right]\frac{\partial^2}{\partial z_{*j}z_{*l}}\\
&\quad=\sum_{j=1}^{L-1}\sum_{l=1}^{L-1}\left[z_{*j}\delta_{jl}-z_{*j}z_{*l}\right]\frac{\partial^2}{\partial z_{*j}z_{*l}}\\
&\quad=\sum_{j=1}^{L-1}\sum_{l=1}^{L-1} z_{*j}(\delta_{jl}-z_{*l})\frac{\partial^2}{\partial z_{*j}z_{*l}}.
\end{aligned}$$

Expanding the fourth term of equation (S.1.2), applying both $x_{ij}(\mathbf{z})=z_{ij}$ for $1\le i\le K-1, 1\le j\le L-1$ and $\sum_{j=1}^{L} x_{ij}(\mathbf{z})=z_{i*}$, we obtain

$$\begin{aligned}
&\sum_{i=1}^{K-1}\sum_{j=1}^{L-1}\sum_{k=1}^{K-1}\sum_{l=1}^{L} x_{ij}(\mathbf{z})(\delta_{ik}\delta_{jl}-x_{kl}(\mathbf{z}))\frac{\partial^2}{\partial z_{ij}z_{k*}}\\
&\quad=\sum_{i=1}^{K-1}\sum_{j=1}^{L-1}\sum_{k=1}^{K-1}\sum_{l=1}^{L} x_{ij}(\mathbf{z})\delta_{ik}\delta_{jl}\frac{\partial^2}{\partial z_{ij}z_{k*}}-\sum_{i=1}^{K-1}\sum_{j=1}^{L-1}\sum_{k=1}^{K-1}\sum_{l=1}^{L} x_{ij}(\mathbf{z})x_{kl}(\mathbf{z})\frac{\partial^2}{\partial z_{ij}z_{k*}}\\
&\quad=\sum_{i=1}^{K-1}\sum_{j=1}^{L-1}\sum_{k=1}^{K-1} z_{ij}\delta_{ik}\frac{\partial^2}{\partial z_{ij}z_{k*}}-\sum_{i=1}^{K-1}\sum_{j=1}^{L-1}\sum_{k=1}^{K-1} z_{ij}z_{k*}\frac{\partial^2}{\partial z_{ij}z_{k*}}\\
&\quad=\sum_{i=1}^{K-1}\sum_{j=1}^{L-1}\sum_{k=1}^{K-1} z_{ij}(\delta_{ik}-z_{k*})\frac{\partial^2}{\partial z_{ij}z_{k*}}.
\end{aligned}$$

We can repeat this same arithmetic logic for the fifth term of equation (S.1.2) to get

$$\sum_{i=1}^{K-1}\sum_{j=1}^{L-1}\sum_{k=1}^{K}\sum_{l=1}^{L-1} x_{ij}(\mathbf{z})(\delta_{ik}\delta_{jl} - x_{kl}(\mathbf{z}))\frac{\partial^2}{\partial z_{ij}z_{*l}}$$
$$= \sum_{i=1}^{K-1}\sum_{j=1}^{L-1}\sum_{k=1}^{K}\sum_{l=1}^{L-1} x_{ij}(\mathbf{z})\delta_{ik}\delta_{jl}\frac{\partial^2}{\partial z_{ij}z_{*l}} - \sum_{i=1}^{K-1}\sum_{j=1}^{L-1}\sum_{k=1}^{K}\sum_{l=1}^{L-1} x_{ij}(\mathbf{z})x_{kl}(\mathbf{z})\frac{\partial^2}{\partial z_{ij}z_{*l}}$$
$$= \sum_{i=1}^{K-1}\sum_{j=1}^{L-1}\sum_{l=1}^{L-1} z_{ij}\delta_{jl}\frac{\partial^2}{\partial z_{ij}z_{*l}} - \sum_{i=1}^{K-1}\sum_{j=1}^{L-1}\sum_{l=1}^{L-1} z_{ij}z_{*l}\frac{\partial^2}{\partial z_{ij}z_{*l}}$$
$$= \sum_{i=1}^{K-1}\sum_{j=1}^{L-1}\sum_{l=1}^{L-1} z_{ij}(\delta_{jl} - z_{*l})\frac{\partial^2}{\partial z_{ij}z_{*l}}.$$

For the sixth term in equation (S.1.2), we expand the same product, then simplify sums that involve the Kronecker delta function to rewrite the term as

$$\sum_{i=1}^{K-1}\sum_{j=1}^{L}\sum_{k=1}^{K-1}\sum_{l=1}^{L-1} x_{ij}(\mathbf{z})(\delta_{ik}\delta_{jl} - x_{kl}(\mathbf{z}))\frac{\partial^2}{\partial z_{i*}z_{kl}}$$
$$= \sum_{i=1}^{K-1}\sum_{j=1}^{L}\sum_{k=1}^{K-1}\sum_{l=1}^{L-1} x_{ij}(\mathbf{z})\delta_{ik}\delta_{jl}\frac{\partial^2}{\partial z_{i*}z_{kl}} - \sum_{i=1}^{K-1}\sum_{j=1}^{L}\sum_{k=1}^{K-1}\sum_{l=1}^{L-1} x_{ij}(\mathbf{z})x_{kl}(\mathbf{z})\frac{\partial^2}{\partial z_{i*}z_{kl}}$$
$$= \sum_{i=1}^{K-1}\sum_{k=1}^{K-1}\sum_{l=1}^{L-1} x_{il}(\mathbf{z})\delta_{ik}\frac{\partial^2}{\partial z_{i*}z_{kl}} - \sum_{i=1}^{K-1}\sum_{k=1}^{K-1}\sum_{l=1}^{L-1} z_{i*}z_{kl}\frac{\partial^2}{\partial z_{i*}z_{kl}}$$
$$= \sum_{i=1}^{K-1}\sum_{k=1}^{K-1}\sum_{l=1}^{L-1} z_{kl}(\delta_{ik} - z_{i*})\frac{\partial^2}{\partial z_{i*}z_{kl}}.$$

Similarly, for the seventh term of equation (S.1.2), we again simplify sums with the Kronecker delta function to obtain

$$\sum_{i=1}^{K-1}\sum_{j=1}^{L}\sum_{k=1}^{K}\sum_{l=1}^{L-1} x_{ij}(\mathbf{z})(\delta_{ik}\delta_{jl} - x_{kl}(\mathbf{z}))\frac{\partial^2}{\partial z_{i*}z_{*l}}$$
$$= \sum_{i=1}^{K-1}\sum_{j=1}^{L}\sum_{k=1}^{K}\sum_{l=1}^{L-1} x_{ij}(\mathbf{z})\delta_{ik}\delta_{jl}\frac{\partial^2}{\partial z_{i*}z_{*l}} - \sum_{i=1}^{K-1}\sum_{j=1}^{L}\sum_{k=1}^{K}\sum_{l=1}^{L-1} x_{ij}(\mathbf{z})x_{kl}(\mathbf{z})\frac{\partial^2}{\partial z_{i*}z_{*l}}$$
$$= \sum_{i=1}^{K-1}\sum_{j=1}^{L}\sum_{l=1}^{L-1} x_{ij}(\mathbf{z})\delta_{jl}\frac{\partial^2}{\partial z_{i*}z_{*l}} - \sum_{i=1}^{K-1}\sum_{l=1}^{L-1} z_{i*}z_{*l}\frac{\partial^2}{\partial z_{i*}z_{*l}}$$
$$= \sum_{i=1}^{K-1}\sum_{l=1}^{L-1} z_{il}\frac{\partial^2}{\partial z_{i*}z_{*l}} - \sum_{i=1}^{K-1}\sum_{l=1}^{L-1} z_{i*}z_{*l}\frac{\partial^2}{\partial z_{i*}z_{*l}}.$$

For to the eighth term of equation (S.1.2), we use the same arithmetic steps as for the sixth term, although applied to $z_{*j}$ rather than $z_{i*}$ to get

$$\sum_{i=1}^{K}\sum_{j=1}^{L-1}\sum_{k=1}^{K-1}\sum_{l=1}^{L-1} x_{ij}(\mathbf{z})(\delta_{ik}\delta_{jl} - x_{kl}(\mathbf{z}))\frac{\partial^2}{\partial z_{*j}z_{kl}}$$
$$= \sum_{i=1}^{K}\sum_{j=1}^{L-1}\sum_{k=1}^{K-1}\sum_{l=1}^{L-1} x_{ij}(\mathbf{z})\delta_{ik}\delta_{jl}\frac{\partial^2}{\partial z_{*j}z_{kl}} - \sum_{i=1}^{K}\sum_{j=1}^{L-1}\sum_{k=1}^{K-1}\sum_{l=1}^{L-1} x_{ij}(\mathbf{z})x_{kl}(\mathbf{z})\frac{\partial^2}{\partial z_{*j}z_{kl}}$$

$$= \sum_{j=1}^{L-1}\sum_{k=1}^{K-1}\sum_{l=1}^{L-1} x_{kj}(\mathbf{z})\delta_{jl}\frac{\partial^2}{\partial z_{*j} z_{kl}} - \sum_{j=1}^{L-1}\sum_{k=1}^{K-1}\sum_{l=1}^{L-1} z_{*j} z_{kl}\frac{\partial^2}{\partial z_{*j} z_{kl}}$$
$$= \sum_{j=1}^{L-1}\sum_{k=1}^{K-1}\sum_{l=1}^{L-1} z_{kl}(\delta_{jl} - z_{*j})\frac{\partial^2}{\partial z_{*j} z_{kl}}.$$

Finally, for to the ninth term of equation (S.1.2), expanding the product and simplifying yields

$$\sum_{i=1}^{K}\sum_{j=1}^{L-1}\sum_{k=1}^{K-1}\sum_{l=1}^{L} x_{ij}(\mathbf{z})(\delta_{ik}\delta_{jl} - x_{kl}(\mathbf{z}))\frac{\partial^2}{\partial z_{*j} z_{k*}}$$
$$= \sum_{i=1}^{K}\sum_{j=1}^{L-1}\sum_{k=1}^{K-1}\sum_{l=1}^{L} x_{ij}(\mathbf{z})\delta_{ik}\delta_{jl}\frac{\partial^2}{\partial z_{*j} z_{k*}} - \sum_{i=1}^{K}\sum_{j=1}^{L-1}\sum_{k=1}^{K-1}\sum_{l=1}^{L} x_{ij}(\mathbf{z})x_{kl}(\mathbf{z})\frac{\partial^2}{\partial z_{*j} z_{k*}}$$
$$= \sum_{i=1}^{K}\sum_{j=1}^{L-1}\sum_{k=1}^{K-1} x_{ij}\delta_{ik}(\mathbf{z})\frac{\partial^2}{\partial z_{*j} z_{k*}} - \sum_{i=1}^{K}\sum_{j=1}^{L-1}\sum_{k=1}^{K-1} x_{ij} z_{k*}\frac{\partial^2}{\partial z_{*j} z_{k*}}$$
$$= \sum_{j=1}^{L-1}\sum_{k=1}^{K-1} z_{kj}\frac{\partial^2}{\partial z_{*j} z_{k*}} - \sum_{j=1}^{L-1}\sum_{k=1}^{K-1} z_{*j} z_{k*}\frac{\partial^2}{\partial z_{*j} z_{k*}}.$$

**To summarize**, we get the drift operator

$$\mathcal{L}_{\mathbf{z}}^{\text{drift}}(t) = \frac{1}{4N_e(t)}\Bigg[\sum_{i=1}^{K-1}\sum_{j=1}^{L-1}\sum_{k=1}^{K-1}\sum_{l=1}^{L-1} z_{ij}(\delta_{ik}\delta_{jl} - z_{kl})\frac{\partial^2}{\partial z_{ij} z_{kl}}$$
$$+ \sum_{i=1}^{K-1}\sum_{k=1}^{K-1} z_{i*}(\delta_{ik} - z_{k*})\frac{\partial^2}{\partial z_{i*} z_{k*}} + \sum_{j=1}^{L-1}\sum_{l=1}^{L-1} z_{*j}(\delta_{jl} - z_{*l})\frac{\partial^2}{\partial z_{*j} z_{*l}}$$
$$+ \sum_{i=1}^{K-1}\sum_{j=1}^{L-1}\sum_{k=1}^{K-1} z_{ij}(\delta_{ik} - z_{k*})\frac{\partial^2}{\partial z_{ij} z_{k*}} + \sum_{i=1}^{K-1}\sum_{j=1}^{L-1}\sum_{l=1}^{L-1} z_{ij}(\delta_{jl} - z_{*l})\frac{\partial^2}{\partial z_{ij} z_{*l}}$$
$$+ \sum_{i=1}^{K-1}\sum_{k=1}^{K-1}\sum_{l=1}^{L-1} z_{kl}(\delta_{ik} - z_{i*})\frac{\partial^2}{\partial z_{i*} z_{kl}} + \sum_{i=1}^{K-1}\sum_{l=1}^{L-1}(z_{il} - z_{i*} z_{*l})\frac{\partial^2}{\partial z_{i*} z_{*l}}$$
$$+ \sum_{j=1}^{L-1}\sum_{k=1}^{K-1}\sum_{l=1}^{L-1} z_{kl}(\delta_{jl} - z_{*j})\frac{\partial^2}{\partial z_{*j} z_{kl}} + \sum_{j=1}^{L-1}\sum_{k=1}^{K-1}(z_{kj} - z_{*j} z_{k*})\frac{\partial^2}{\partial z_{*j} z_{k*}}\Bigg]$$

in the coordinates $\mathbf{z}$. □

*Proof of Lemma 3.2 in the main text.* Before proving the form of the mutation operator in the $\mathbf{z}$ coordinates, it is helpful to note that the relations

(S.1.3) $$x_{Kl} = x_{\cdot l} - \sum_{k=1}^{K-1} x_{kl} \qquad \text{and} \qquad x_{kL} = x_{k\cdot} - \sum_{l=1}^{L-1} x_{kl},$$

and

$$x_{KL} = 1 - \sum_{i=1}^{K}\sum_{j=1}^{L} \mathbf{1}_{\{(ij)\neq(KL)\}} x_{kl}$$
$$= 1 - \sum_{k=1}^{K-1}\sum_{l=1}^{L} x_{kl} - \sum_{l=1}^{L-1} x_{Kl}$$

$$= 1 - \sum_{k=1}^{K-1} x_{k\cdot} - \sum_{l=1}^{L-1} x_{\cdot l} + \sum_{l=1}^{L-1}\sum_{k=1}^{K-1} x_{kl}$$

hold. Thus,

$$\begin{aligned}
\sum_{l=1}^{L} x_{Kl} &= \sum_{l=1}^{L-1} x_{Kl} + x_{KL} \\
&= \sum_{l=1}^{L-1} x_{\cdot l} - \sum_{l=1}^{L-1}\sum_{k=1}^{K-1} x_{kl} + 1 - \sum_{k=1}^{K-1} x_{k\cdot} - \sum_{l=1}^{L-1} x_{\cdot l} + \sum_{l=1}^{L-1}\sum_{k=1}^{K-1} x_{kl} \qquad \text{(S.1.4)} \\
&= 1 - \sum_{k=1}^{K-1} x_{k\cdot}
\end{aligned}$$

holds, and similarly

$$\sum_{k=1}^{K} x_{kL} = 1 - \sum_{l=1}^{L-1} x_{\cdot l}.$$

Now, by following our method from the section on drift, we change coordinates for the mutation operator defined in equation (2.4) in the main text using the chain rule:

$$\begin{aligned}
\mathcal{L}_{\mathbf{x}}^{\text{mut}} = &\sum_{i=1}^{K-1}\sum_{j=1}^{L-1}\left[\sum_{l=1}^{L} x_{il}(\mathbf{z}) m_{l,j}^{(2)} + \sum_{k=1}^{K} x_{kj}(\mathbf{z}) m_{k,i}^{(1)}\right]\frac{\partial}{\partial z_{ij}} \\
&+ \sum_{i=1}^{K-1}\sum_{j=1}^{L}\left[\sum_{l=1}^{L} x_{il}(\mathbf{z}) m_{l,j}^{(2)} + \sum_{k=1}^{K} x_{kj}(\mathbf{z}) m_{k,i}^{(1)}\right]\frac{\partial}{\partial z_{i*}} \qquad \text{(S.1.5)} \\
&+ \sum_{i=1}^{K}\sum_{j=1}^{L-1}\left[\sum_{l=1}^{L} x_{il}(\mathbf{z}) m_{l,j}^{(2)} + \sum_{k=1}^{K} x_{kj}(\mathbf{z}) m_{k,i}^{(1)}\right]\frac{\partial}{\partial z_{*j}},
\end{aligned}$$

where $m_{i,j}^{(p)}$ is the per generation mutation probability defined in Lemma 3.2 in the main text. This is a sum with three terms, which we will simplify term-wise. The first term in equation (S.1.5) can be modified as follows:

$$\begin{aligned}
&\sum_{i=1}^{K-1}\sum_{j=1}^{L-1}\left[\sum_{l=1}^{L} x_{il}(\mathbf{z}) m_{l,j}^{(2)} + \sum_{k=1}^{K} x_{kj}(\mathbf{z}) m_{k,i}^{(1)}\right]\frac{\partial}{\partial z_{ij}} \\
&= \sum_{i=1}^{K-1}\sum_{j=1}^{L-1}\left[\sum_{l=1}^{L-1} m_{l,j}^{(2)} x_{il}(\mathbf{z}) + m_{L,j}^{(2)} x_{i\cdot}(\mathbf{z}) - m_{L,j}^{(2)} \sum_{l=1}^{L-1} x_{il}(\mathbf{z}) \right. \\
&\qquad \left. + \sum_{k=1}^{K-1} m_{k,i}^{(1)} x_{kj}(\mathbf{z}) + m_{K,i}^{(1)} x_{\cdot j}(\mathbf{z}) - m_{K,i}^{(1)} \sum_{k=1}^{K-1} x_{kj}(\mathbf{z})\right]\frac{\partial}{\partial z_{ij}},
\end{aligned}$$

where we substituted equation (S.1.3) for $x_{Kj}(\mathbf{z})$ and $x_{iL}(\mathbf{z})$.

Furthermore, the second term in equation (S.1.5) yields

$$\begin{aligned}
&\sum_{i=1}^{K-1}\sum_{j=1}^{L}\left[\sum_{l=1}^{L} x_{il}(\mathbf{z}) m_{l,j}^{(2)} + \sum_{k=1}^{K} x_{kj}(\mathbf{z}) m_{k,i}^{(1)}\right]\frac{\partial}{\partial z_{i*}} \\
&= \sum_{i=1}^{K-1}\left[\sum_{l=1}^{L} x_{il}(\mathbf{z}) \sum_{j=1}^{L} m_{l,j}^{(2)} + \sum_{k=1}^{K} m_{k,i}^{(1)} \sum_{j=1}^{L} x_{kj}(\mathbf{z})\right]\frac{\partial}{\partial z_{i*}}
\end{aligned}$$

$$= \sum_{i=1}^{K-1} \left[ \sum_{k=1}^{K-1} m_{k,i}^{(1)} x_{k\cdot}(\mathbf{z}) + m_{K,i}^{(1)} \sum_{j=1}^{L} x_{Kj}(\mathbf{z}) \right] \frac{\partial}{\partial z_{i*}}$$

$$= \sum_{i=1}^{K-1} \left[ \sum_{k=1}^{K-1} m_{k,i}^{(1)} x_{k\cdot}(\mathbf{z}) + m_{K,i}^{(1)} - m_{K,i}^{(1)} \sum_{k=1}^{K-1} x_{k\cdot}(\mathbf{z}) \right] \frac{\partial}{\partial z_{i*}},$$

where we used the fact that $\sum_{j=1}^{L} m_{l,j}^{(2)} = 0$, and substituted equation (S.1.4) for $\sum_{j=1}^{L} x_{Kj}(\mathbf{z})$.

The third term of equation (S.1.5) follows similarly. With $x_{ij}(\mathbf{z}) = z_{ij}$ for $i < K$ and $j < L$, $x_{i\cdot}(\mathbf{z}) = z_{i*}$ for $i < K$, and $x_{\cdot j}(\mathbf{z}) = z_{*j}$ for $j < L$, we can then write the fully simplified mutation operator in $\mathbf{z}$ coordinates as

$$\mathcal{L}_{\mathbf{z}}^{\text{mut}} = \sum_{i=1}^{K-1} \sum_{j=1}^{L-1} \left[ \sum_{l=1}^{L-1} (m_{l,j}^{(2)} - m_{L,j}^{(2)}) z_{il} + m_{L,j}^{(2)} z_{i*} + \sum_{k=1}^{K-1} (m_{k,i}^{(1)} - m_{K,i}^{(1)}) z_{kj} + m_{K,i}^{(1)} z_{*j} \right] \frac{\partial}{\partial z_{ij}}$$
$$+ \sum_{i=1}^{K-1} \left[ \sum_{k=1}^{K-1} (m_{k,i}^{(1)} - m_{K,i}^{(1)}) z_{k*} + m_{K,i}^{(1)} \right] \frac{\partial}{\partial z_{i*}} + \sum_{j=1}^{L-1} \left[ \sum_{l=1}^{L-1} (m_{l,j}^{(2)} - m_{L,j}^{(2)}) z_{*l} + m_{L,j}^{(2)} \right] \frac{\partial}{\partial z_{*j}}.$$

□

*Proof of Lemma 3.3 in the main text.* We can rewrite the recombination generator for $\mathbf{x}$ in the same way we modified the drift and the mutation operator: Applying the chain rule with respect to $x_{ij}$,

$$\sum_{i=1}^{K-1} \sum_{j=1}^{L-1} r(x_{i\cdot} x_{\cdot j} - x_{ij}) \left( \frac{\partial}{\partial z_{ij}} + \frac{\partial}{\partial z_{i*}} + \frac{\partial}{\partial z_{*j}} \right)$$
$$+ \sum_{i=1}^{K-1} r(x_{i\cdot} x_{\cdot L} - x_{iL}) \frac{\partial}{\partial z_{i*}} + \sum_{j=1}^{L-1} r(x_{K\cdot} x_{\cdot j} - x_{Kj}) \frac{\partial}{\partial z_{*j}},$$

and then grouping the common partials together

$$\sum_{i=1}^{K-1} \sum_{j=1}^{L-1} r(x_{i\cdot} x_{\cdot j} - x_{ij}) \frac{\partial}{\partial z_{ij}} + \sum_{i=1}^{K-1} \sum_{j=1}^{L} r(x_{i\cdot} x_{\cdot j} - x_{ij}) \frac{\partial}{\partial z_{i*}} + \sum_{i=1}^{K} \sum_{j=1}^{L-1} r(x_{i\cdot} x_{\cdot j} - x_{ij}) \frac{\partial}{\partial z_{*j}}.$$

We now can notice that the second and third terms are both zero, so, in the $\mathbf{z}$ coordinates, we write

$$\mathcal{L}_{z}^{\text{reco}} = \sum_{i=1}^{K-1} \sum_{j=1}^{L-1} r(z_{i*} z_{*j} - z_{ij}) \frac{\partial}{\partial z_{ij}}.$$

□

S.1.2. **Differential equations for closed moments.** The following proofs detail the derivation of the ODEs for $G_t(\mathbf{a}, \mathbf{b}, \mathbf{c})$ by applying the differential operators and using Dynkin's formula.

*Proof of Theorem 3.5 in the main text.* We will now derive the ODE by applying the drift generator $\mathcal{L}_{\mathbf{z}}^{\text{drift}}(t)$ to the closed moments $G_t(\mathbf{a}, \mathbf{b}, \mathbf{c})$. Again, we omit the coefficient $\frac{1}{4N_e(t)}$ for convenience. Beginning with the first term of the generator (S.1.2), we can take the derivative of the moment function and then absorb the variables $z_{ij}$ and $z_{kl}$ again. This allows us to rearrange the summands and obtain:

$$\sum_{i=1}^{K-1} \sum_{j=1}^{L-1} \left[ \sum_{k=1}^{K-1} \sum_{l=1}^{L-1} z_{ij} (\delta_{ik} \delta_{jl} - z_{kl}) \frac{\partial^2}{\partial z_{ij} z_{kl}} \right] G_t(\mathbf{a}, \mathbf{b}, \mathbf{c})$$
$$= \sum_{i=1}^{K-1} \sum_{j=1}^{L-1} \sum_{k=1}^{K-1} \sum_{l=1}^{L-1} z_{ij} (\delta_{ik} \delta_{jl} - z_{kl}) a_{ij} (a_{kl} - \delta_{ik} \delta_{jl}) G_t(\mathbf{a} - \mathbf{e}_{ij} - \mathbf{e}_{kl}, \mathbf{b}, \mathbf{c})$$

$$= \sum_{i=1}^{K-1}\sum_{j=1}^{L-1} a_{ij}(a_{ij}-1)G_t(\mathbf{a}-\mathbf{e}_{ij},\mathbf{b},\mathbf{c}) - \sum_{i=1}^{K-1}\sum_{j=1}^{L-1}\sum_{k=1}^{K-1}\sum_{l=1}^{L-1} a_{ij}(a_{kl}-\delta_{ik}\delta_{jl})G_t(\mathbf{a},\mathbf{b},\mathbf{c})$$

$$= \sum_{i=1}^{K-1}\sum_{j=1}^{L-1}\Big[a_{ij}(a_{ij}-1)G_t(\mathbf{a}-\mathbf{e}_{ij},\mathbf{b},\mathbf{c}) - \sum_{k=1}^{K-1}\sum_{l=1}^{L-1} a_{ij}(a_{kl}-\delta_{ik}\delta_{jl})G_t(\mathbf{a},\mathbf{b},\mathbf{c})\Big].$$

The second and third term of the generator (S.1.2) applied to the moment function can be resolved in a similar way. We will now look at the fourth term of the generator (3.2) in the main text acting on the moment function, again taking the derivative, re-absorbing $\mathbf{z}$ coordinates in the moments, and rearranging terms yields

$$\begin{aligned}
&\sum_{i=1}^{K-1}\sum_{j=1}^{L-1}\sum_{k=1}^{K-1} z_{ij}(\delta_{ik}-z_{k*})\frac{\partial^2}{\partial z_{ij}z_{k*}}G_t(\mathbf{a},\mathbf{b},\mathbf{c})\\
&= \sum_{i=1}^{K-1}\sum_{j=1}^{L-1}\sum_{k=1}^{K-1} z_{ij}(\delta_{ik}-z_{k*})a_{ij}b_kG_t(\mathbf{a}-\mathbf{e}_{ij},\mathbf{b}-\mathbf{e}_k,\mathbf{c})\\
&= \sum_{i=1}^{K-1}\sum_{j=1}^{L-1} a_{ij}b_iG_t(\mathbf{a},\mathbf{b}-\mathbf{e}_i,\mathbf{c}) - \sum_{i=1}^{K-1}\sum_{j=1}^{L-1}\sum_{k=1}^{K-1} a_{ij}b_kG_t(\mathbf{a},\mathbf{b},\mathbf{c})\\
&= \sum_{i=1}^{K-1}\sum_{j=1}^{L-1}\left[a_{ij}b_iG_t(\mathbf{a},\mathbf{b}-\mathbf{e}_i,\mathbf{c}) - \sum_{k=1}^{K-1} a_{ij}b_kG_t(\mathbf{a},\mathbf{b},\mathbf{c})\right].
\end{aligned} \tag{S.1.6}$$

With the sixth term of the generator (S.1.2), we take the derivative and rearrange the summands to obtain

$$\begin{aligned}
&\sum_{i=1}^{K-1}\sum_{k=1}^{K-1}\sum_{l=1}^{L-1} z_{kl}(\delta_{ik}-z_{i*})\frac{\partial^2}{\partial z_{i*}z_{kl}}G_t(\mathbf{a},\mathbf{b},\mathbf{c})\\
&= \sum_{i=1}^{K-1}\sum_{k=1}^{K-1}\sum_{l=1}^{L-1} z_{kl}(\delta_{ik}-z_{i*})a_{kl}b_iG_t(\mathbf{a}-\mathbf{e}_{kl},\mathbf{b}-\mathbf{e}_i,\mathbf{c})\\
&= \sum_{k=1}^{K-1}\sum_{l=1}^{L-1} a_{kl}b_kG_t(\mathbf{a},\mathbf{b}-\mathbf{e}_k,\mathbf{c}) - \sum_{i=1}^{K-1}\sum_{k=1}^{K-1}\sum_{l=1}^{L-1} a_{kl}b_iG_t(\mathbf{a},\mathbf{b},\mathbf{c})\\
&= \sum_{i=1}^{K-1}\sum_{j=1}^{L-1}\left[a_{ij}b_iG_t(\mathbf{a},\mathbf{b}-\mathbf{e}_i,\mathbf{c}) - \sum_{k=1}^{K-1} a_{ij}b_kG_t(\mathbf{a},\mathbf{b},\mathbf{c})\right],
\end{aligned} \tag{S.1.7}$$

where the last line simply involves a relabeling of $i$ to $k$, and $l$ to $j$, and vice-versa. We now notice that the last lines of (S.1.6) and (S.1.7) are equal, and thus can be combined into

$$2\sum_{i=1}^{K-1}\sum_{j=1}^{L-1}\Big[a_{ij}b_iG_t(\mathbf{a},\mathbf{b}-\mathbf{e}_i,\mathbf{c}) - \sum_{k=1}^{K-1} a_{ij}b_kG_t(\mathbf{a},\mathbf{b},\mathbf{c})\Big].$$

The fifth and eighth term of the generator (S.1.2) applied to the moment function behave similarly to the fourth and and sixth, respectively; thus, they can be combined into

$$2\sum_{i=1}^{K-1}\sum_{j=1}^{L-1}\Big[a_{ij}c_jG_t(\mathbf{a},\mathbf{b},\mathbf{c}-\mathbf{e}_j) - \sum_{l=1}^{L-1} a_{ij}c_lG_t(\mathbf{a},\mathbf{b},\mathbf{c})\Big].$$

There are now only two remaining terms in the generator (S.1.2) to consider, the seventh and the ninth. For the seventh term, taking the derivative and expanding the product yields

$$\begin{aligned}
&\sum_{i=1}^{K-1}\sum_{l=1}^{L-1}(z_{il}-z_{i*}z_{*l})\frac{\partial^2}{\partial z_{i*}z_{*l}}G_t(\mathbf{a},\mathbf{b},\mathbf{c})\\
&=\sum_{i=1}^{K-1}\sum_{l=1}^{L-1}(z_{il}-z_{i*}z_{*l})b_ic_lG_t(\mathbf{a},\mathbf{b}-\mathbf{e}_i,\mathbf{c}-\mathbf{e}_l)\\
&=\sum_{i=1}^{K-1}\sum_{l=1}^{L-1}b_ic_lG_t(\mathbf{a}+\mathbf{e}_{il},\mathbf{b}-\mathbf{e}_i,\mathbf{c}-\mathbf{e}_l)-b_ic_lG(\mathbf{a},\mathbf{b},\mathbf{c}),
\end{aligned}$$

and we observe that the ninth term can be simplified similarly.

**To summarize**, the drift operator in $\mathbf{z}$ coordinates applied to the moment function takes the form

$$\begin{aligned}
&\sum_{i=1}^{K-1}\sum_{j=1}^{L-1}\Big[a_{ij}(a_{ij}-1)G_t(\mathbf{a}-\mathbf{e}_{ij},\mathbf{b},\mathbf{c})-\sum_{k=1}^{K-1}\sum_{l=1}^{L-1}a_{ij}(a_{kl}-\delta_{ij}\delta_{kl})G_t(\mathbf{a},\mathbf{b},\mathbf{c})\Big]\\
&+\sum_{i=1}^{K-1}\Big[b_i(b_i-1)G_t(\mathbf{a},\mathbf{b}-\mathbf{e}_i,\mathbf{c})-\sum_{k=1}^{K-1}b_i(b_k-\delta_{ik})G_t(\mathbf{a},\mathbf{b},\mathbf{c})\Big]\\
&+\sum_{j=1}^{L-1}\Big[c_j(c_j-1)G_t(\mathbf{a},\mathbf{b},\mathbf{c}-\mathbf{e}_j)-\sum_{l=1}^{L-1}c_j(c_l-\delta_{jl})G_t(\mathbf{a},\mathbf{b},\mathbf{c})\Big]\\
&+2\sum_{i=1}^{K-1}\sum_{j=1}^{L-1}\Big[a_{ij}b_iG_t(\mathbf{a},\mathbf{b}-\mathbf{e}_i,\mathbf{c})-\sum_{k=1}^{K-1}a_{ij}b_kG_t(\mathbf{a},\mathbf{b},\mathbf{c})\Big]\\
&+2\sum_{i=1}^{K-1}\sum_{j=1}^{L-1}\Big[a_{ij}c_jG_t(\mathbf{a},\mathbf{b},\mathbf{c}-\mathbf{e}_j)-\sum_{l=1}^{L-1}a_{ij}c_lG_t(\mathbf{a},\mathbf{b},\mathbf{c})\Big]\\
&+\sum_{i=1}^{K-1}\sum_{l=1}^{L-1}b_ic_lG_t(\mathbf{a}+\mathbf{e}_{il},\mathbf{b}-\mathbf{e}_i,\mathbf{c}-\mathbf{e}_l)-b_ic_lG_t(\mathbf{a},\mathbf{b},\mathbf{c})\\
&+\sum_{j=1}^{L-1}\sum_{k=1}^{K-1}b_kc_jG_t(\mathbf{a}+\mathbf{e}_{kj},\mathbf{b}-\mathbf{e}_k,\mathbf{c}-\mathbf{e}_j)-b_kc_jG_t(\mathbf{a},\mathbf{b},\mathbf{c}).
\end{aligned}$$

We can rearrange this by grouping terms to obtain:

$$\begin{aligned}
&-\Big[\sum_{i=1}^{K-1}\sum_{j=1}^{L-1}\sum_{k=1}^{K-1}\sum_{l=1}^{L-1}a_{ij}(a_{kl}-\delta_{ij}\delta_{kl})+\sum_{i=1}^{K-1}\sum_{k=1}^{K-1}b_i(b_k-\delta_{ik})+\sum_{j=1}^{L-1}\sum_{l=1}^{L-1}c_j(c_l-\delta_{jl})\\
&+2\sum_{i=1}^{K-1}\sum_{j=1}^{L-1}\sum_{k=1}^{K-1}a_{ij}b_k+2\sum_{i=1}^{K-1}\sum_{j=1}^{L-1}\sum_{l=1}^{L-1}a_{ij}c_l+2\sum_{i=1}^{K-1}\sum_{l=1}^{L-1}b_ic_l\Big]G_t(\mathbf{a},\mathbf{b},\mathbf{c})\\
&+\sum_{i=1}^{K-1}\sum_{j=1}^{L-1}a_{ij}(a_{ij}-1)G_t(\mathbf{a}-\mathbf{e}_{ij},\mathbf{b},\mathbf{c})+\Big[\sum_{i=1}^{K-1}b_i(b_i-1)+2\sum_{i=1}^{K-1}\sum_{j=1}^{L-1}a_{ij}b_i\Big]G_t(\mathbf{a},\mathbf{b}-\mathbf{e}_i,\mathbf{c})\\
&+\Big[\sum_{j=1}^{L-1}c_j(c_j-1)+2\sum_{i=1}^{K-1}\sum_{j=1}^{L-1}a_{ij}c_j\Big]G_t(\mathbf{a},\mathbf{b},\mathbf{c}-\mathbf{e}_j)+2\sum_{i=1}^{K-1}\sum_{l=1}^{L-1}b_ic_lG_t(\mathbf{a}+\mathbf{e}_{il},\mathbf{b}-\mathbf{e}_i,\mathbf{c}-\mathbf{e}_l).
\end{aligned}$$

□

*Proof of Theorem 3.6 in the main text.* We can now apply the converted mutation operator to the moments $G_t(\mathbf{a},\mathbf{b},\mathbf{c})$, taking the respective derivatives and re-absorbing coordinates, to obtain

$$\begin{aligned}\mathcal{L}^{\mathrm{mut}}_{\mathbf{z}} G_t(\mathbf{a},\mathbf{b},\mathbf{c}) = \sum_{i=1}^{K-1}\sum_{j=1}^{L-1} a_{ij}\Bigg[&\sum_{l=1}^{L-1}(m^{(2)}_{l,j}-m^{(2)}_{L,j})G_t(\mathbf{a}-\mathbf{e}_{ij}+\mathbf{e}_{il},\mathbf{b},\mathbf{c}) + m^{(2)}_{L,j}G_t(\mathbf{a}-\mathbf{e}_{ij},\mathbf{b}+\mathbf{e}_i,\mathbf{c})\\ &+\sum_{k=1}^{K-1}(m^{(1)}_{k,i}-m^{(1)}_{K,i})G_t(\mathbf{a}-\mathbf{e}_{ij}+\mathbf{e}_{kj},\mathbf{b},\mathbf{c}) + m^{(1)}_{K,i}G_t(\mathbf{a}-\mathbf{e}_{ij},\mathbf{b},\mathbf{c}+\mathbf{e}_j)\Bigg]\\ +\sum_{i=1}^{K-1} b_i\Bigg[&\sum_{k=1}^{K-1}(m^{(1)}_{k,i}-m^{(1)}_{K,i})G_t(\mathbf{a},\mathbf{b}-\mathbf{e}_i+\mathbf{e}_k,\mathbf{c}) + m^{(1)}_{K,i}G_t(\mathbf{a},\mathbf{b}-\mathbf{e}_i,\mathbf{c})\Bigg]\\ +\sum_{j=1}^{L-1} c_j\Bigg[&\sum_{l=1}^{L-1}(m^{(2)}_{l,j}-m^{(2)}_{L,j})G_t(\mathbf{a},\mathbf{b},\mathbf{c}-\mathbf{e}_j+\mathbf{e}_l) + m^{(2)}_{L,j}G_t(\mathbf{a},\mathbf{b},\mathbf{c}-\mathbf{e}_j)\Bigg],\end{aligned}$$

which can be expanded to obtain

$$\begin{aligned}\mathcal{L}^{\mathrm{mut}}_{\mathbf{z}} G_t(\mathbf{a},\mathbf{b},\mathbf{c}) = &\sum_{i=1}^{K-1}\sum_{j=1}^{L-1}\sum_{\substack{l=1\\ l\neq j}}^{L-1} a_{ij}(m^{(2)}_{l,j}-m^{(2)}_{L,j})G_t(\mathbf{a}-\mathbf{e}_{ij}+\mathbf{e}_{il},\mathbf{b},\mathbf{c})\\ &+\sum_{i=1}^{K-1}\sum_{j=1}^{L-1}\sum_{\substack{k=1\\ k\neq i}}^{K-1} a_{ij}(m^{(1)}_{k,i}-m^{(1)}_{K,i})G_t(\mathbf{a}-\mathbf{e}_{ij}+\mathbf{e}_{kj},\mathbf{b},\mathbf{c})\\ &+\sum_{i=1}^{K-1}\sum_{j=1}^{L-1} a_{ij}m^{(2)}_{L,j}G_t(\mathbf{a}-\mathbf{e}_{ij},\mathbf{b}+\mathbf{e}_i,\mathbf{c}) + \sum_{i=1}^{K-1}\sum_{j=1}^{L-1} a_{ij}m^{(1)}_{K,i}G_t(\mathbf{a}-\mathbf{e}_{ij},\mathbf{b},\mathbf{c}+\mathbf{e}_j)\\ &+\sum_{i=1}^{K-1}\sum_{\substack{k=1\\ k\neq i}}^{K-1} b_i(m^{(1)}_{k,i}-m^{(1)}_{K,i})G_t(\mathbf{a},\mathbf{b}-\mathbf{e}_i+\mathbf{e}_k,\mathbf{c}) + \sum_{i=1}^{K-1} b_i m^{(1)}_{K,i}G_t(\mathbf{a},\mathbf{b}-\mathbf{e}_i,\mathbf{c})\\ &+\sum_{j=1}^{L-1}\sum_{\substack{l=1\\ \neq j}}^{L-1} c_j(m^{(2)}_{l,j}-m^{(2)}_{L,j})G_t(\mathbf{a},\mathbf{b},\mathbf{c}-\mathbf{e}_j+\mathbf{e}_l) + \sum_{j=1}^{L-1} c_j m^{(2)}_{L,j}G_t(\mathbf{a},\mathbf{b},\mathbf{c}-\mathbf{e}_j)\\ &+\Bigg[\sum_{i=1}^{K-1}(m^{(1)}_{i,i}-m^{(1)}_{K,i})\Big(b_i+\sum_{j=1}^{L-1}a_{ij}\Big)\\ &\quad+\sum_{j=1}^{L-1}(m^{(2)}_{j,j}-m^{(2)}_{L,j})\Big(c_j+\sum_{i=1}^{K-1}a_{ij}\Big)\Bigg]G_t(\mathbf{a},\mathbf{b},\mathbf{c}).\end{aligned} \tag{S.1.8}$$

□

*Proof of Corollary 3.7 in the main text.* We can simplify equation (S.1.8) further by substituting the general mutation model with a parent-independent mutation model. Under this parent-independent mutation model, we have $m^{(1)}_{i,k} = m^{(1)}_k$ for all $i,k \in \{1,\dots,K\}$ with $i \neq k$. For $i = k$, we have

$$m^{(1)}_{i,i} = -\sum_{\substack{k=1\\ k\neq i}}^{K} m^{(1)}_{i,k} = -\sum_{\substack{k=1\\ k\neq i}}^{K} m^{(1)}_{k}.$$

Similar relations hold for $m^{(2)}$. Substituting this into equation (S.1.8), we can simplify to

$$\begin{aligned}\mathcal{L}_{\mathbf{z}}^{\text{mut}} G_t(\mathbf{a},\mathbf{b},\mathbf{c}) = &\sum_{j=1}^{L-1} m_j^{(2)} \sum_{i=1}^{K-1} a_{ij} G_t(\mathbf{a}-\mathbf{e}_{ij},\mathbf{b}+\mathbf{e}_i,\mathbf{c}) + \sum_{i=1}^{K-1} m_i^{(1)} \sum_{j=1}^{L-1} a_{ij} G_t(\mathbf{a}-\mathbf{e}_{ij},\mathbf{b},\mathbf{c}+\mathbf{e}_j) \\ &+ \sum_{i=1}^{K-1} b_i m_i^{(1)} G_t(\mathbf{a},\mathbf{b}-\mathbf{e}_i,\mathbf{c}) + \sum_{j=1}^{L-1} c_j m_j^{(2)} G_t(\mathbf{a},\mathbf{b},\mathbf{c}-\mathbf{e}_j) \\ &- \Big[\Big(\sum_{k=1}^{K} m_k^{(1)}\Big) \sum_{i=1}^{K-1} \Big(b_i + \sum_{j=1}^{L-1} a_{ij}\Big) + \Big(\sum_{l=1}^{L} m_l^{(2)}\Big) \sum_{j=1}^{L-1} \Big(c_j + \sum_{i=1}^{K-1} a_{ij}\Big)\Big] G_t(\mathbf{a},\mathbf{b},\mathbf{c}).\end{aligned}$$

□

### S.2. Equivalence of differential operators for different dimensions

As detailed in Section 2.1 in the main text, the Wright-Fisher diffusion can be described as a stochastic process on $\Delta_H$, positive haplotype frequencies for $h \in H$ that sum to 1 or less, where implicitly $x_{KL} = 1 - \sum_{h \in H} x_h$. Alternatively, the Wright-Fisher diffusion can also be introduced as a stochastic process on $\partial\Delta_{\overline{H}}$, the boundary of the simplex $\Delta_{\overline{H}}$; that is, positive haplotype frequencies for all $h \in \overline{H}$ that sum to exactly 1. In the following sections, we show that the diffusion generator for genetic drift, recurrent mutation, and recombination, respectively, can be defined in either representation, and that the two definitions are equivalent.

#### S.2.1. Genetic Drift.

For genetic drift, the fact that we have two loci is actually not important, and we can just work on the level of the genetic types $\bar{H} = \{1, \ldots, |\bar{H}|\}$ ordered in a way that $\bar{H}$ includes the last genetic type and $H$ excludes it. The two possible ways to formulate the drift generator are

$$\mathcal{L}_H^{\text{drift}} := \frac{1}{4N_e(t)} \sum_{h=1}^{|H|} \sum_{g=1}^{|H|} x_h(\delta_{hg} - x_g) \frac{\partial}{\partial x_h} \frac{\partial}{\partial x_g}$$

on the full simplex $\Delta_H$, see equation (2.2) in the main text, and

$$\mathcal{L}_{\bar{H}}^{\text{drift}} := \frac{1}{4N_e(t)} \sum_{h=1}^{|\bar{H}|} \sum_{g=1}^{|\bar{H}|} y_h(\delta_{hg} - y_g) \frac{\partial}{\partial y_h} \frac{\partial}{\partial y_g}$$

on the boundary of the simplex $\partial\Delta_{\overline{H}}$. We will now show that the generator $\mathcal{L}_{\bar{H}}^{\text{drift}}$ on $\partial\Delta_{\overline{H}}$ describes the same dynamics as the generator $\mathcal{L}_H^{\text{drift}}$ on $\Delta_H$. To this end, note that using the chain rule, we have

$$\begin{aligned}\frac{\partial}{\partial x_h} &= \sum_{g=1}^{|\bar{H}|} \frac{\partial y_g}{\partial x_h} \frac{\partial}{\partial y_g} \\ &= \frac{\partial}{\partial y_h} + \frac{\partial y_{|\bar{H}|}}{\partial y_h} \frac{\partial}{\partial y_{|\bar{H}|}} \qquad \text{(S.2.1)} \\ &= \frac{\partial}{\partial y_h} - \frac{\partial}{\partial y_{|\bar{H}|}},\end{aligned}$$

since $y_{|\bar{H}|} = 1 - \sum_{g\in H} y_g$. Then, the full simplex drift generator becomes

$$
\begin{aligned}
4N_e(t)\mathcal{L}_H^{\text{drift}} &= \sum_{h=1}^{|H|}\sum_{g=1}^{|H|} x_h(\delta_{hg} - x_g)\frac{\partial}{\partial x_h}\frac{\partial}{\partial x_g} \\
&= \sum_{h=1}^{|H|}\sum_{g=1}^{|H|} y_h(\delta_{hg} - y_g)\left(\frac{\partial}{\partial y_h} - \frac{\partial}{\partial y_{|\bar{H}|}}\right)\left(\frac{\partial}{\partial y_g} - \frac{\partial}{\partial y_{|\bar{H}|}}\right) \\
&= \sum_{h=1}^{|H|}\sum_{g=1}^{|H|} y_h(\delta_{hg} - y_g)\frac{\partial}{\partial y_h}\frac{\partial}{\partial y_g} - \sum_{h=1}^{|H|}\sum_{g=1}^{|H|} y_h(\delta_{hg} - y_g)\frac{\partial}{\partial y_{|\bar{H}|}}\frac{\partial}{\partial y_g} \\
&\quad - \sum_{h=1}^{|H|}\sum_{g=1}^{|H|} y_h(\delta_{hg} - y_g)\frac{\partial}{\partial y_h}\frac{\partial}{\partial y_{|\bar{H}|}} + \sum_{h=1}^{|H|}\sum_{g=1}^{|H|} y_h(\delta_{hg} - y_g)\frac{\partial}{\partial y_{|\bar{H}|}}\frac{\partial}{\partial y_{|\bar{H}|}}.
\end{aligned} \tag{S.2.2}
$$

The first term can be left for now. For the second term, we get

$$
\begin{aligned}
&-\sum_{h=1}^{|H|}\sum_{g=1}^{|H|} y_h(\delta_{hg} - y_g)\frac{\partial}{\partial y_{|\bar{H}|}}\frac{\partial}{\partial y_g} \\
&\qquad = -\sum_{g=1}^{|H|}\left[\sum_{h=1}^{|H|} y_h(\delta_{hg} - y_g)\right]\frac{\partial}{\partial y_{|\bar{H}|}}\frac{\partial}{\partial y_g} \\
&\qquad = -\sum_{g=1}^{|H|}\left[y_g - \sum_{h=1}^{|H|} y_h y_g\right]\frac{\partial}{\partial y_{|\bar{H}|}}\frac{\partial}{\partial y_g} \\
&\qquad = -\sum_{g=1}^{|H|} y_{|\bar{H}|} y_g\frac{\partial}{\partial y_{|\bar{H}|}}\frac{\partial}{\partial y_g},
\end{aligned}
$$

and, by the same arithmetic, the third term yields

$$
-\sum_{h=1}^{|H|}\sum_{g=1}^{|H|} y_h(\delta_{hg} - y_g)\frac{\partial}{\partial y_h}\frac{\partial}{\partial y_{|\bar{H}|}} = -\sum_{h=1}^{|H|} y_h y_{|\bar{H}|}\frac{\partial}{\partial y_h}\frac{\partial}{\partial y_{|\bar{H}|}},
$$

and the fourth term yields

$$
\begin{aligned}
&\sum_{h=1}^{|H|}\sum_{g=1}^{|H|} y_h(\delta_{hg} - y_g)\frac{\partial}{\partial y_{|\bar{H}|}}\frac{\partial}{\partial y_{|\bar{H}|}} \\
&\qquad = \sum_{h=1}^{|H|} y_h y_{|\bar{H}|}\frac{\partial}{\partial y_{|\bar{H}|}}\frac{\partial}{\partial y_{|\bar{H}|}} \\
&\qquad = y_{|\bar{H}|}(1 - y_{|\bar{H}|})\frac{\partial}{\partial y_{|\bar{H}|}}\frac{\partial}{\partial y_{|\bar{H}|}}.
\end{aligned}
$$

Substituting these equations back into equation (S.2.2), we obtain

$$
\begin{aligned}
4N_e(t)\mathcal{L}_H^{\text{drift}} &= \sum_{h=1}^{|\bar{H}|}\sum_{g=1}^{|\bar{H}|} y_h(\delta_{hg} - y_g)\frac{\partial}{\partial y_h}\frac{\partial}{\partial y_g} \\
&= 4N_e(t)\mathcal{L}_{\bar{H}}^{\text{drift}},
\end{aligned}
$$

which shows that the two version of the drift generator describe the same dynamics.

S.2.2. **Recurrent Mutation.** Next, we are interested in the two formulations for the generator of the mutation process. For the general mutation case, the two-locus structure of our genetic types can again be ignored, and we just need to consider the type space $\bar{H} = \{1, \ldots, |\bar{H}|\}$. Further, for $i, j \in \bar{H}$ and $i \neq j$, let $m_{i,j}$ be the per generation probability that genetic type $i$ mutates into genetic type $j$. Moreover, define $m_{i,i} := -\sum_{\substack{j=1 \\ j\neq i}}^{|\bar{H}|} m_{i,j}$. Then, the full simplex version of the mutation generator is given by

$$
\begin{aligned}
\mathcal{L}_H^{\text{mut}} &:= \sum_{i=1}^{|H|} \sum_{\substack{j=1 \\ j\neq i}}^{|\bar{H}|} \left[x_j m_{j,i} - x_i m_{i,j}\right] \frac{\partial}{\partial x_i} \\
&= \sum_{i=1}^{|H|} \sum_{j=1}^{|\bar{H}|} x_j m_{j,i} \frac{\partial}{\partial x_i},
\end{aligned} \tag{S.2.3}
$$

see equation (2.3) in the main text. Furthermore, the simplex boundary version is given by

$$
\begin{aligned}
\mathcal{L}_{\bar{H}}^{\text{mut}} &:= \sum_{i=1}^{|\bar{H}|} \sum_{\substack{j=1 \\ j\neq i}}^{|\bar{H}|} \left[y_j m_{j,i} - y_i m_{i,j}\right] \frac{\partial}{\partial y_i} \\
&= \sum_{i=1}^{|\bar{H}|} \sum_{j=1}^{|\bar{H}|} y_j m_{j,i} \frac{\partial}{\partial y_i}.
\end{aligned}
$$

Again, to show that the two versions are equivalent, we substitute the relation (S.2.1) into definition (S.2.3) and obtain

$$
\begin{aligned}
\mathcal{L}_H^{\text{mut}} &= \sum_{i=1}^{|H|} \sum_{\substack{j=1 \\ j\neq i}}^{|\bar{H}|} \left[y_j m_{j,i} - y_i m_{i,j}\right] \left(\frac{\partial}{\partial y_i} - \frac{\partial}{\partial y_{|\bar{H}|}}\right) \\
&= \sum_{i=1}^{|H|} \sum_{\substack{j=1 \\ j\neq i}}^{|\bar{H}|} \left[y_j m_{j,i} - y_i m_{i,j}\right] \frac{\partial}{\partial y_i} - \sum_{i=1}^{|H|} \sum_{\substack{j=1 \\ j\neq i}}^{|\bar{H}|} \left[y_j m_{j,i} - y_i m_{i,j}\right] \frac{\partial}{\partial y_{|\bar{H}|}}.
\end{aligned} \tag{S.2.4}
$$

Focusing on the second summand, ignoring the partial derivative, we obtain

$$
\begin{aligned}
&- \sum_{i=1}^{|H|} \sum_{\substack{j=1 \\ j\neq i}}^{|\bar{H}|} \left[y_j m_{j,i} - y_i m_{i,j}\right] \\
&= - \sum_{i=1}^{|\bar{H}|} \sum_{j=1}^{|\bar{H}|} \mathbf{1}_{\{i\neq|\bar{H}|\}} \mathbf{1}_{\{i\neq j\}} \left[y_j m_{j,i} - y_i m_{i,j}\right] \\
&= - \left(\sum_{i=1}^{|\bar{H}|} \sum_{j=1}^{|\bar{H}|} \mathbf{1}_{\{i\neq|\bar{H}|\}} \mathbf{1}_{\{i\neq j\}} y_j m_{j,i}\right) + \left(\sum_{i=1}^{|\bar{H}|} \sum_{j=1}^{|\bar{H}|} \mathbf{1}_{\{i\neq|\bar{H}|\}} \mathbf{1}_{\{i\neq j\}} y_i m_{i,j}\right) \\
&= - \left(\sum_{i=1}^{|\bar{H}|} \sum_{j=1}^{|\bar{H}|} \mathbf{1}_{\{i\neq|\bar{H}|\}} \mathbf{1}_{\{i\neq j\}} y_j m_{j,i}\right) + \left(\sum_{j=1}^{|\bar{H}|} \sum_{i=1}^{|\bar{H}|} \mathbf{1}_{\{j\neq|\bar{H}|\}} \mathbf{1}_{\{j\neq i\}} y_j m_{j,i}\right)
\end{aligned}
$$

$$
\begin{aligned}
&= -\Big(\sum_{i=1}^{|H|}\sum_{\substack{j=1\\ j\neq i}}^{|\bar H|} y_j m_{j,i}\Big) + \Big(\sum_{i=1}^{|\bar H|}\sum_{\substack{j=1\\ j\neq i}}^{|H|} y_j m_{j,i}\Big)\\
&= -\Big(\sum_{i=1}^{|H|}\sum_{\substack{j=1\\ j\neq i}}^{|H|} y_j m_{j,i}\Big) + \Big(\sum_{i=1}^{|H|}\sum_{\substack{j=1\\ j\neq i}}^{|H|} y_j m_{j,i}\Big) - \sum_{i=1}^{|H|} y_{|\bar H|} m_{|\bar H|,i} + \sum_{j=1}^{|H|} y_j m_{j,|\bar H|}\\
&= \sum_{j=1}^{|H|} \big[y_j m_{j,|\bar H|} - y_{|\bar H|} m_{|\bar H|,j}\big].
\end{aligned}
$$

Substituting this expression back into equation (S.2.4) shows that

$$
\begin{aligned}
\mathcal{L}_H^{\text{mut}} &= \sum_{i=1}^{|\bar H|}\sum_{\substack{j=1\\ j\neq i}}^{|\bar H|} \big[y_j m_{j,i} - y_i m_{i,j}\big]\frac{\partial}{\partial y_i}\\
&= \mathcal{L}_{\bar H}^{\text{mut}}
\end{aligned}
$$

holds.

S.2.3. **Recombination.** For recombination, the fact that the a genetic type consists of two loci with a certain set of alleles has to be taken into account. Thus, the generator will have sums over the alleles at the first locus $E^{(1)} = \{1,\dots,K\}$, and the alleles at the second locus $E^{(2)} = \{1,\dots,L\}$. Denote by $r$ the per generation recombination probability between the two loci. Then the two formulations for the generator are

$$
\mathcal{L}_H^{\text{reco}} := r\sum_{i=1}^{K}\sum_{j=1}^{L} \mathbf{1}_{\{(ij)\neq(KL)\}}(x_{i\cdot}x_{\cdot j} - x_{ij})\frac{\partial}{\partial x_{ij}}, \tag{S.2.5}
$$

see equation (2.5) in the main text, and

$$
\mathcal{L}_{\bar H}^{\text{reco}} := r\sum_{i=1}^{K}\sum_{j=1}^{L} (y_{i\cdot}y_{\cdot j} - y_{ij})\frac{\partial}{\partial y_{ij}},
$$

Similar to the cases of genetic drift and mutation, we show $\mathcal{L}_H^{\text{reco}}$ defines the same dynamics as $\mathcal{L}_{\bar H}^{\text{reco}}$ by substituting relation (S.2.1) into definition (S.2.5) to obtain

$$
\begin{aligned}
\frac{1}{r}\mathcal{L}_H^{\text{reco}} &= \sum_{i=1}^{K}\sum_{j=1}^{L} \mathbf{1}_{\{(ij)\neq(KL)\}}(x_{i\cdot}x_{\cdot j} - x_{ij})\frac{\partial}{\partial x_{ij}}\\
&= \sum_{i=1}^{K}\sum_{j=1}^{L} \mathbf{1}_{\{(ij)\neq(KL)\}}(y_{i\cdot}y_{\cdot j} - y_{ij})\Big(\frac{\partial}{\partial y_{ij}} - \frac{\partial}{\partial y_{KL}}\Big)\\
&= \sum_{i=1}^{K}\sum_{j=1}^{L} \mathbf{1}_{\{(ij)\neq(KL)\}}(y_{i\cdot}y_{\cdot j} - y_{ij})\frac{\partial}{\partial y_{ij}} - \sum_{i=1}^{K}\sum_{j=1}^{L} \mathbf{1}_{\{(ij)\neq(KL)\}} y_{i\cdot}y_{\cdot j}\frac{\partial}{\partial y_{KL}}\\
&\quad + \sum_{i=1}^{K}\sum_{j=1}^{L} \mathbf{1}_{\{(ij)\neq(KL)\}} y_{ij}\frac{\partial}{\partial y_{KL}}.
\end{aligned} \tag{S.2.6}
$$

Focusing on the second summand, we obtain

$$
\begin{aligned}
&-\sum_{i=1}^{K}\sum_{j=1}^{L} \mathbf{1}_{\{(ij)\neq(KL)\}} y_{i\cdot}y_{\cdot j}\frac{\partial}{\partial y_{KL}}\\
&\qquad = -\sum_{i=1}^{K}\sum_{j=1}^{L} y_{i\cdot}y_{\cdot j}\frac{\partial}{\partial y_{KL}} + y_{K\cdot}y_{\cdot L}\frac{\partial}{\partial y_{KL}}\\
&\qquad = -1\cdot\frac{\partial}{\partial y_{KL}} + y_{K\cdot}y_{\cdot L}\frac{\partial}{\partial y_{KL}},
\end{aligned}
$$

and the third summand can be simplified to

$$\sum_{i=1}^{K}\sum_{j=1}^{L}\mathbf{1}_{\{(ij)\neq(KL)\}}y_{ij}\frac{\partial}{\partial y_{KL}} = (1-y_{KL})\frac{\partial}{\partial y_{KL}}.$$

Again, substituting these relations into equation (S.2.6) yields

$$\begin{aligned}\frac{1}{r}\mathcal{L}_{H}^{\text{reco}} &= \sum_{i=1}^{K}\sum_{j=1}^{L}(y_{i\cdot}y_{\cdot j}-y_{ij})\frac{\partial}{\partial y_{ij}}\\ &= \frac{1}{r}\mathcal{L}_{\tilde{H}}^{\text{reco}},\end{aligned}$$

demonstrating that these two formulations of the generator for recombination are equivalent. With that, we have demonstrated that the two definitions of the complete generator are equivalent.
